# Topographic reference modelling of white matter hyperintensities across the adult lifespan

**DOI:** 10.64898/2026.10.28.760476

**Authors:** Mohamad J. Alshikho, Jeffrey D. Pyne, Natalie C. Edwards, Christiane Hale, Dejania Cotton-Samuel, Clarissa D. Morales, Natalie Morel, Rafael Vignoli Lippert, Derek B. Archer, Michael L. Cuccaro, Arthur W. Toga, Andrew J. Saykin, Shannon L. Risacher, Michael E. Belloy, Walter Kukull, Sarah Biber, Jesse Mez, Paul K. Crane, Shubhabrata Mukherjee, Guray Erus, Reisa A. Sperling, David A. Bennett, Lisa L. Barnes, Julie A. Schneider, Sterling C. Johnson, Annie J. Lee, Christos Davatzikos, Sid O’Bryant, Robert C. Barber, Carlos Cruchaga, Tammie L.S. Benzinger, Dong Young Lee, Dahyun Yi, A4 Study team, HABS-HD Study Team, Alzheimer’s Disease Neuroimaging Initiative (ADNI), Alzheimer’s Disease Sequencing Project (ADSP), Timothy J. Hohman, Jennifer J. Manly, Richard Mayeux, Mohamad Habes, Patrick J. Lao, Adam M. Brickman

## Abstract

White matter hyperintensities (WMH) are among the most common MRI findings in older adults. They are widely regarded as markers of brain aging and cerebral small-vessel disease, and associated with risk for cognitive decline, Alzheimer’s disease and related dementias (ADRD), and stroke. White matter hyperintensities (WMH) are typically interpreted by total lesion burden, but lesions with similar volumes can occupy different anatomical locations, potentially reflecting distinct pathological processes. Here we present voxelwise topographic reference modelling (vTRM), a framework for characterizing individual WMH masks against age- and sex-stratified frequency maps derived from cognitively unimpaired individuals. Using 42,469 FLAIR-derived lesion masks from twelve adult cohorts, we mapped population-level WMH spatial vulnerability, characterized age-related changes in its distribution across periventricular, lobar and arterial-territory compartments, and constructed paired percentile charts for lesion burden and spatial typicality. A quadrant-based classification defined by these two percentile dimensions revealed graded relationships with ADRD pathophysiology, neurodegeneration, and cognitive impairment, while identifying WMH patterns differentially associated with ageing-related, vascular and neurodegenerative processes among individuals with similar lesion volumes. These reference charts establish a benchmark for normative WMH ageing, enabling quantification of individual deviations from typical age-, sex- and burden-related patterns and identification of WMH phenotypes associated with vascular, neurodegenerative or mixed etiologies. The framework, together with a deployable 3D U-Net model trained on these data for automated WMH segmentation, is available through the Brain Lesion Analytics platform (https://brainlesionanalytics.org/).

## Introduction

White matter hyperintensities (WMH) are among the most common findings on magnetic resonance imaging (MRI) in older adults [^1,2^]. They are associated with vascular risk factors, cognitive decline and dementia, and are widely studied in research on cognitive ageing and Alzheimer’s disease (AD) as markers of cerebral small-vessel disease [^3^]. The severity and clinical relevance of WMH are assessed primarily by lesion burden, most often expressed as total lesion volume.

This volume-centred view is simple, reproducible and clinically intuitive, but it is incomplete because WMH are spatially heterogeneous. Lesions preferentially involve periventricular white matter, extend into ventricle-distal regions at older ages and vary across lobar and vascular territories [^4^]. Two individuals with similar WMH burden may therefore have markedly different lesion topographies. Treating these patterns as equivalent may obscure biologically distinct processes and limits interpretation of spatial vulnerability [^5^]. Depending on their anatomical distribution, WMH have been linked to ageing [^6^], hypertension [^7^], diabetes [^8^], small-vessel disease [^9^], amyloid and tau pathology [^10^], atrophy and impaired brain clearance, and regional tissue vulnerability [^11,12^]. Previous work suggests that WMH location carries regionally distinct associations with vascular risk and AD pathology [^13^]. Global lesion volume therefore collapses potentially informative spatial signals into a single scalar measure, limiting the study of mixed vascular and neurodegenerative processes.

Despite their ubiquity with increasing age, the radiologic assessment of WMH burden remains largely dependent on subjective visual interpretation, with findings frequently described clinically as “mild”, “moderate”, or “severe” in the absence of standardized operational definitions or quantitative severity anchors. Many biological measures, however, are most informative when interpreted formally against an expected population distribution. Paediatric growth charts, for example, allow clinicians to determine whether a child’s height or weight is typical for age and sex, rather than interpreting the raw value in isolation. Brain charts have extended this principle to structural brain measures across the lifespan [^14^], and burden-based WMH reference charts now allow total lesion volume to be expressed relative to age- and sex-specific expectations [^15^]. However, these charts address lesion quantity, not lesion location; they do not additionally indicate whether the spatial distribution of an individual’s lesions is atypical for their age, sex and total burden [^16^]. Building such a spatial reference requires voxelwise lesion maps in a common space from large samples, which heterogeneous acquisition and segmentation protocols have so far made difficult.

Here, we develop voxelwise topographic reference modelling (vTRM), a population-scale framework for characterizing the spatial distribution of WMH relative to age-, sex- and burden-specific expectations. Using lesion masks derived from 42,469 fluid-attenuated inversion recovery (FLAIR) scans from the Alzheimer’s Disease Sequencing Project Phenotype Harmonization Consortium (ADSP-PHC) [^17,18^], we define reference maps of WMH spatial frequency and age-related topographic variation across periventricular, ventricle-distal, lobar and vascular-territory compartments. We further construct paired percentile charts for total lesion burden and spatial typicality, defined as the degree to which an individual’s lesion distribution matches that expected for their age, sex and burden. Together, these charts enable: (1) standardized quantification of whether an individual’s lesion burden is typical for age and sex; (2) assessment of whether an individual’s lesion location follows the expected topographic pattern for their burden; (3) identification of atypical lesion topographies that may suggest mixed vascular and neurodegenerative processes; and (4) a quadrant-based stratification of individuals according to WMH burden and spatial typicality. We do not claim that spatial typicality alone provides a definitive diagnostic classifier. Rather, this work establishes a reference framework for interpreting WMH as both a volumetric and a spatial phenotype, providing a foundation for investigating vascular brain injury and mixed pathophysiology in ageing, AD and related dementias.

## Results

### Mapping population-level WMH burden across the adult lifespan

We analysed 42,469 standardized FLAIR-derived WMH lesion masks from 22,084 participants across twelve contributing cohorts, spanning 21–103 years of age (mean 72.5 ± 8.9 years; median 72.8 years, interquartile range 67.1–78.4). The sample comprised 16,807 scans from men (39.6%) and 25,662 from women (60.4%). All lesion masks were represented in MNI152 2 mm standard space. The largest contributing cohorts were NACC/SCAN (30.9%), ADNI (17.0%), A4 (14.0%) and HABS-HD (13.8%), with additional data from MAP, WRAP, WHICAP, Knight ADRC, KBASE, LEARN, MARS and ROS. Of the full sample, 27,247 scans from cognitively unimpaired participants were used to establish normative WMH reference models (Fig. 1a, Extended Data Table 1, Supplementary Fig. S1.F1 and Supplementary Table S1.T2). ComBat harmonisation, evaluated as a sensitivity analysis, left all associations unchanged in direction and significance (Supplementary Fig. S1.F3).

**Figure 1.**
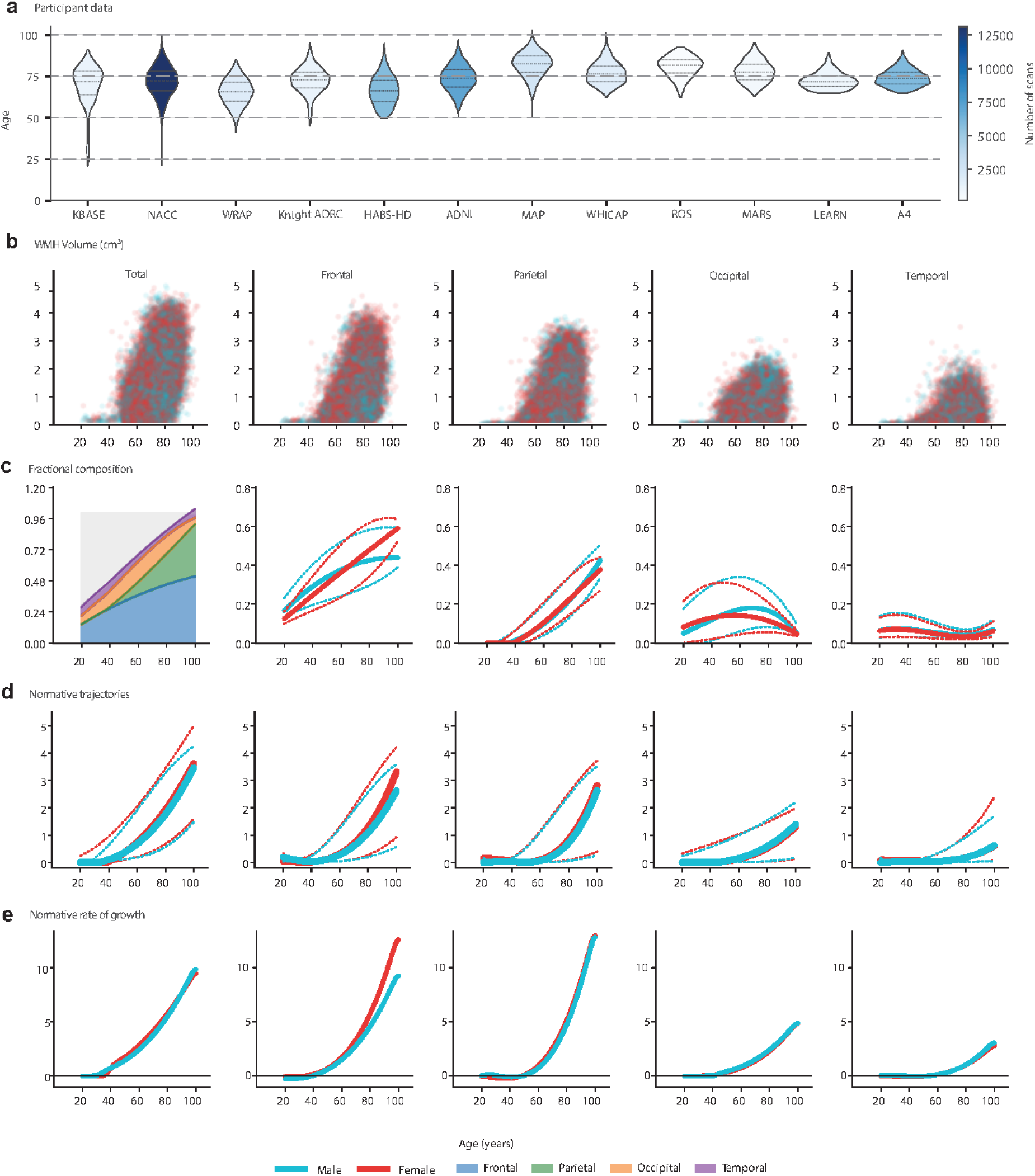
Total and lobar WMH volume charts across the adult lifespan. a, Data from 12 imaging cohorts within the ADSP-PHC consortium, comprising 42,469 FLAIR-derived WMH lesion masks from participants aged 21–103 years. Violin plots show cohort-specific age distributions, coloured by the number of cognitively unimpaired participants. b, Observed total and lobar WMH volumes across the full analytic sample (42,469 scans). Each point represents one scan, including repeated scans from the same participant. Volumes are shown as log(1 + volume in cm³), separately for male (cyan) and female (red) participants. Columns represent total, frontal, parietal, occipital and temporal WMH volume. c, Age-related lobar composition of WMH in cognitively unimpaired scans (male, n = 9,818; female, n = 17,429). The first panel shows median fractions across frontal, temporal, parietal, occipital and other compartments, summarized across sexes. Remaining panels show sex-stratified fractional reference curves estimated using quantile regression, with medians (solid lines) and 25th and 75th percentiles (dotted lines). d, Sex-stratified normative trajectories of total and lobar WMH volume in cognitively unimpaired scans, estimated using cubic quantile regression in age without cohort covariates. Solid lines indicate medians and dotted lines indicate the 5th and 95th percentiles. Shared y axes facilitate comparison of WMH burden across lobes. e, Age-related slopes of fitted median log-transformed WMH volume curves across lobar compartments. Slopes characterize differences between ages in the cross-sectional normative reference, rather than within-participant lesion accumulation. The horizontal line indicates zero slope; shared y axes facilitate comparison across lobes.

The distribution of WMH volume was strongly right-skewed. Sex-stratified quantile regression (QR) was used to estimate age-related reference curves for total and regional WMH burden. Compared with Gaussian-family generalized additive models (GAMs), QR provided similar scan rank-ordering but better agreement with the observed upper-tail quantiles and fewer implausible curve features at the age extremes. Comparisons with generalized additive models for location, scale and shape (GAMLSS) supported these findings. QR reference curves closely matched empirical decade-specific quantiles, particularly in the distribution tails (Supplementary Figs. S2.F1–S2.F4 and Supplementary Tables S2.T1–S2.T4).

Cohort-adjusted models indicated predominantly intercept-level differences, supporting pooled reference estimation without cohort indicators. Fitted quantiles remained ordered across the principal reference age range, with crossings restricted to sparsely sampled extremes. Bootstrap confidence bands were narrower in densely sampled age ranges and wider at the extremes. Refitting the reference using one scan per participant produced nearly identical percentile curves, although confidence bands were wider when observations were independent. Among cognitively unimpaired participants with repeated scans, regional WMH burden percentiles showed moderate-to-strong longitudinal correspondence despite within-participant variation, with no systematic age-related drift. Leave-one-cohort-out analyses indicated that between-cohort differences predominantly affected reference levels rather than age-related curve shapes. Percentile assignments were largely insensitive to the choice of log transformation, with differences concentrated in regional compartments containing exact zero volumes. Subsampling analyses showed that deviations from the full-sample reference decreased approximately with the inverse square root of sample size, reaching 0.5–1.9% in volume terms at ages 60–80 years, with larger deviations at sparsely sampled age extremes (Supplementary Figs. S2.F5–S2.F12 and Supplementary Tables S2.T5–S2.T12).

### Regional WMH phenotypes across the adult lifespan

Regional WMH burden was characterized across periventricular, ventricle-distal, lobar and arterial-territory compartments using QR (Figs. 1b–e and 2b–d). Across the full sample, WMH were predominantly periventricular, with 56.3% of lesion volume within 10 mm of the ventricular system, 41.1% in ventricle-distal white matter and 2.6% within the ventricular mask, potentially reflecting choroid plexus or CSF flow artifacts. Regional composition, quantified as pooled lesion-volume fractions across scans within each age group, differed across age groups. Among individuals younger than 50 years, periventricular and ventricle-distal fractions were 58% and 33%, respectively, compared with 43% and 56% at age 90 years and older. The periventricular fraction was highest in the sixth decade (68%), whereas ventricle-distal WMH accounted for a larger fraction than periventricular WMH after age 80. Arterial-territory distributions showed similar age-related differences: MCA-territory fractions were 30% and 53% in the youngest and oldest age groups, respectively, whereas PCA-territory fractions were 27% and 11%.

**Figure 2.**
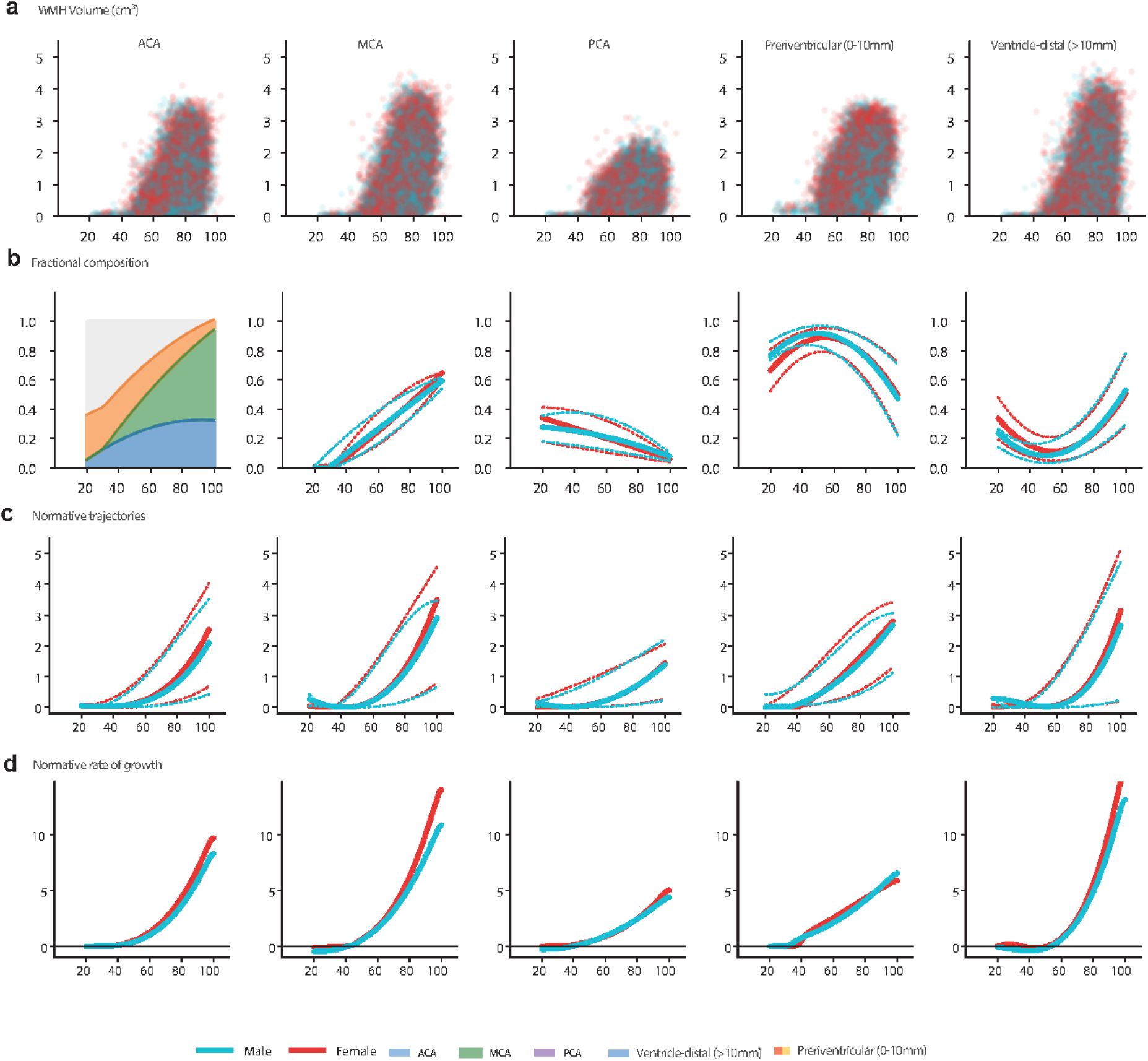
Depth- and vascular-territory WMH charts across the adult lifespan. a, Raw data points show periventricular, ventricle-distal, anterior cerebral artery (ACA), middle cerebral artery (MCA) and posterior cerebral artery (PCA) territory WMH volumes across the adult lifespan for male (cyan) and female (red) participants (n = 42,469 scans). Volumes are displayed as log(1 + volume in cm³). b, Regional WMH composition in cognitively unimpaired scans (male, n = 9,818; female, n = 17,429). The first panel shows median regional fractions summarized across sexes. Remaining panels show sex-stratified fractional trajectories estimated using quantile regression, with the median (solid lines) and 25th and 75th percentiles (dotted lines). c, Sex-stratified normative trajectories for periventricular, ventricle-distal, ACA-, MCA- and PCA-territory WMH volume, estimated using cubic quantile regression in age. Median (solid lines) and 5th and 95th percentiles (dotted lines) are shown. Shared y axes enable comparison of regional WMH burden. d, Normative age-related slopes of the fitted median log-transformed WMH volume trajectories, calculated with respect to log(age). The horizontal line (y = 0) denotes points where the fitted slope changes sign. Shared y axes enable comparison of slope magnitudes across regions.

Lobar distributions showed corresponding age-related differences. Frontal and parietal WMH fractions were 33% and 20%, respectively, among individuals younger than 50 years, compared with 52% and 32% at age 90 years and older. Conversely, occipital fractions were 24% and 10%, and temporal fractions were 8% and 5%, respectively. Despite their smaller fractions at older ages, absolute posterior and PCA-territory WMH burdens were higher, indicating that fractional differences did not reflect lower absolute posterior lesion burden.

Relative to anatomical compartment size, occipital and PCA-territory WMH were 2.9-fold and 1.8-fold over-represented, respectively, among individuals younger than 50 years, whereas MCA-territory WMH were represented at 0.6 times the proportion expected from compartment size. Across older age groups, greater fractions of WMH were observed in ventricle-distal white matter, frontal and parietal lobes, and MCA territory. By the seventh decade, anterior regions accounted for a greater proportion of WMH than posterior regions (Supplementary Fig. S3.F3 and Supplementary Table S3.T3).

### Age-related milestones of regional WMH burden

We characterized regional age-related slopes as derivatives of sex-stratified median QR curves across six age strata. These slopes represent cross-sectional differences in fitted WMH burden rather than observed within-individual growth. Periventricular white matter was the only compartment with a positive slope in the youngest age stratum (0.66, 95% CI 0.49–0.86). Positive slopes were also evident in frontal, MCA- and ACA-territory regions during the sixth decade, whereas ventricle-distal slopes remained near zero. At ages 60–70 years, parietal and ventricle-distal slopes were steeper than occipital, PCA-territory and temporal slopes. At older ages, MCA-territory slopes exceeded periventricular slopes, with relatively steep ventricle-distal slopes (Fig. 3a, b).

**Figure 3.**
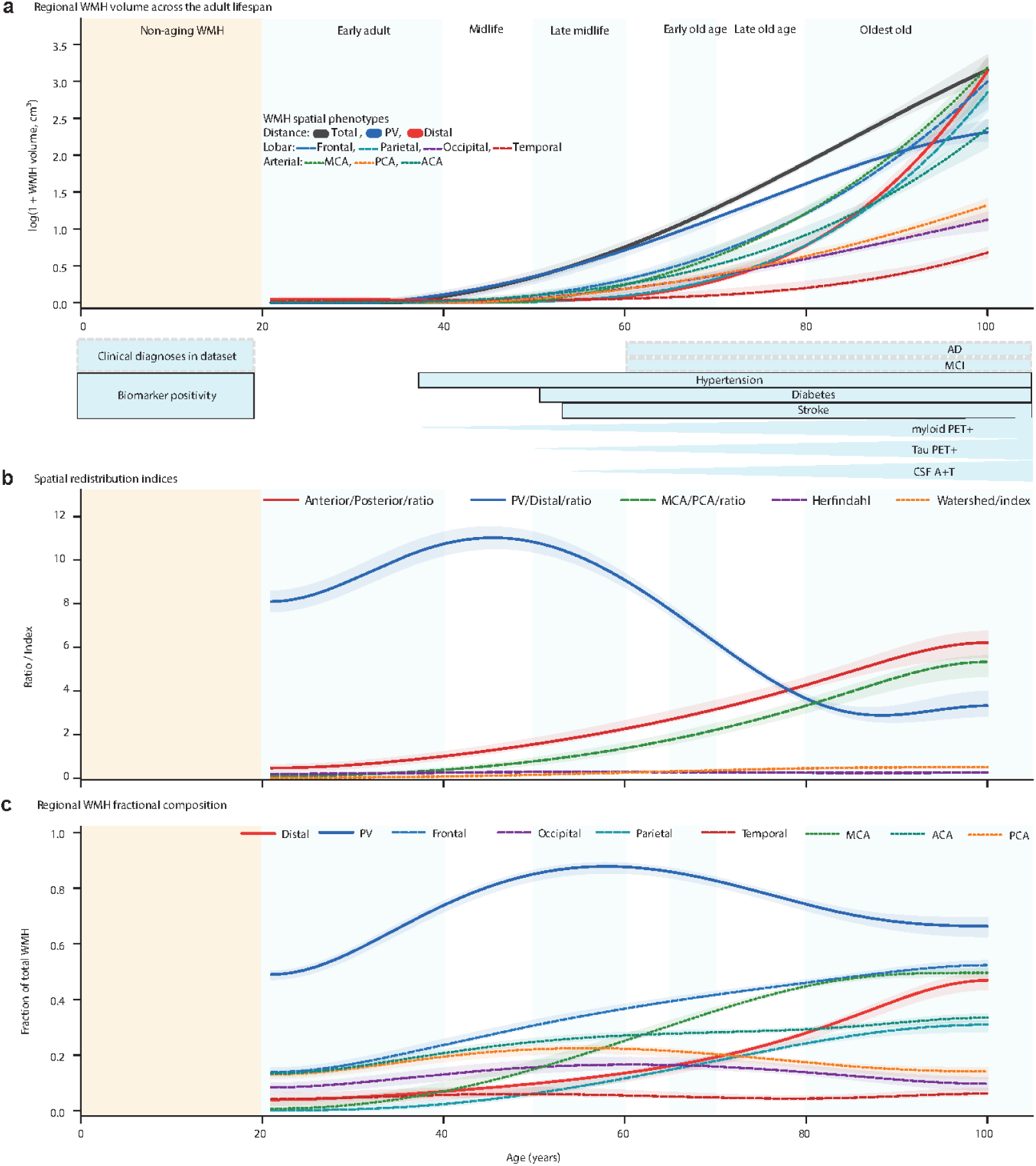
WMH spatial phenotype trajectories and age-related milestones across the adult lifespan. a, Median WMH volume trajectories for ten spatial compartments across 27,247 cognitively unimpaired scans, modelled using fourth-degree polynomial quantile regression of log-transformed WMH volume. Solid lines denote total, periventricular (PV) and ventricle-distal WMH; dashed lines denote frontal, temporal, parietal and occipital regions; and dotted lines denote middle cerebral artery (MCA), posterior cerebral artery (PCA) and anterior cerebral artery (ACA) territories. Shaded bands represent 95% bootstrap confidence intervals from 200 iterations. Annotations below the trajectories summarize age ranges associated with clinical diagnoses, biomarker positivity and vascular risk factors. Tapered bars indicate higher estimated prevalence at older ages. Light orange shading marks ages 0–20 years, outside the analysed age range, and light grey vertical lines delimit the labelled lifespan epochs. b, Age-related trajectories of five spatial redistribution indices: anterior-to-posterior WMH ratio, PV-to-distal ratio, MCA-to-PCA ratio, Herfindahl territorial concentration index and MCA-territory fraction. Higher Herfindahl values indicate greater concentration of WMH within fewer arterial territories. c, Age-related fractional composition of WMH across the spatial compartments, expressed as the proportion of total lesion volume within each compartment.

Model-estimated milestones were defined by zero crossings of regional slope curves (Fig. 4a–c). The periventricular zero crossing was estimated at 23.9 years (95% CI 20.8–29.7) but could not be reliably resolved near the lower age boundary, occurring within the evaluated range in only 132 of 200 bootstrap replicates. Other regional milestones were more consistently estimated: PCA, occipital, frontal, ACA and MCA territories at 39.6, 40.1, 43.5, 43.7 and 45.4 years, respectively, followed by parietal, temporal and ventricle-distal compartments at 52.5, 54.1 and 54.8 years. MCA-territory slopes first exceeded periventricular slopes at 68.0 years (95% CI 66.9–69.0), followed by ventricle-distal slopes at 78.7 years (95% CI 76.4–81.9). Leave-one-study-out analyses supported the stability of most milestone estimates, with greater uncertainty for the periventricular boundary estimate (Supplementary Figs. S3.F1–S3.F2 and Supplementary Tables S3.T1–S3.T2).

**Figure 4.**
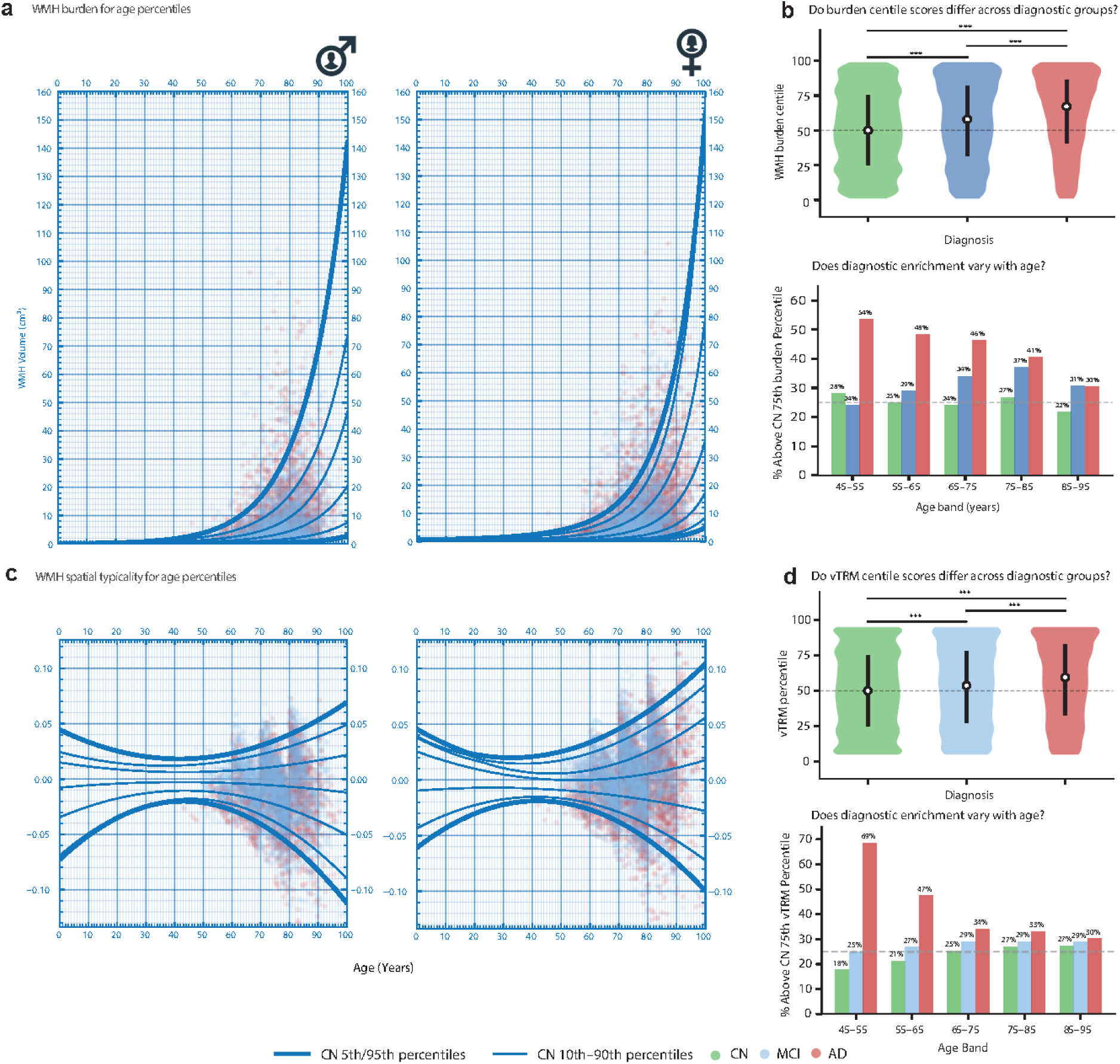
WMH burden and spatial typicality percentile charts with diagnostic validation. a, Normative WMH volume-for-age percentile charts for male (left) and female (right) cognitively unimpaired participants, estimated using quadratic quantile regression in age. The 5^th^, 10^th^, 25^th^, 50^th^, 75^th^, 90^th^ and 95^th^ percentile trajectories are displayed on the original volume scale (cm³), with the 5^th^ and 95^th^ percentiles highlighted. Grid lines follow CDC growth-chart conventions. b, Diagnostic differences in WMH burden percentiles. Violin plots show percentile distributions among cognitively unimpaired, mild cognitive impairment (MCI) and Alzheimer’s disease (AD) scans, with white dots indicating medians. Bar plots show the percentage of scans exceeding the 75^th^ percentile across age groups; the dashed line indicates the expected 25% in the cognitively unimpaired reference. Asterisks denote P < 0.001 (Mann–Whitney U-test). c, Sex-stratified normative percentile charts for voxelwise topographic reference modelling (vTRM) scores in cognitively unimpaired participants, estimated using quadratic quantile regression in age. Higher vTRM percentiles indicate greater lesion-location atypicality relative to the normative reference. Fitted percentile-band widths followed a U-shaped pattern, narrowing to a minimum around ages 46–47 years before widening at older ages. Empirical score dispersion confirmed the older-age widening but showed no clear increase below age 50, where relatively few scans were available. The younger-age portion of the fitted U-shape should therefore be interpreted cautiously. d, Diagnostic differences in vTRM percentiles. Violin plots show percentile distributions across scans from cognitively unimpaired individuals, individuals with MCI, and individuals with AD. Bar plots show the percentage exceeding the 75^th^ percentile across age groups, with the dashed line indicating the expected 25% reference. The highest proportion of scans exceeding the 75th percentile among individuals with AD is observed at ages 45–55 years.

To contextualize these milestones, we examined concurrent vascular, molecular, and diagnostic measures. Earlier milestones could not be contextualized because fewer than 50 participants contributed corresponding measures below age 43 years. Across the parietal milestone (52.5 years) and the MCA and ventricle-distal crossovers (68.0 and 78.7 years), estimated amyloid positivity was 8.1%, 24.2% and 40.6%, respectively, with corresponding tau Braak ≥3 positivity of 3.3%, 10.6% and 17.2%. Hypertension prevalence was also high at these ages (59.9%, 75.5% and 85.8%), alongside higher prevalence of MCI and AD dementia at older milestone ages. These estimates were standardized within measure-specific study sets and should not be interpreted as direct comparisons between different clinical or biological measures (Supplementary Fig. S3.F4 and Supplementary Table S3.T4).

Spatial indices provided convergent evidence of age-related differences in regional WMH composition (Fig. 3b). The anterior-to-posterior ratio was higher at age 90 than at age 30 (5.59 versus 0.52), as was the MCA-to-PCA ratio (4.42 versus 0.07). The periventricular-to-distal ratio was highest at age 47 (9.30, 95% CI 8.91–9.74) and was lower at age 90 (1.69, 95% CI 1.53–1.86). Similarly, the Herfindahl–Hirschman index was lower at age 90 than at age 50 (0.21 versus 0.27), consistent with a more spatially distributed WMH pattern at older ages (Supplementary Table S3.T3b).

### Burden percentile charts for individual WMH scoring

To provide an age- and sex-specific reference for WMH burden, we constructed percentile charts from cognitively unimpaired scans using QR (Fig. 4a). The charts covered the 5^th^–95^th^ percentiles of log-transformed WMH volume and were well calibrated: the median reference percentile was 50.0, with 25.0% and 10.0% of scans exceeding the 75^th^ and 90^th^ percentiles, respectively. Percentile assignments from a GAM comparator were closely concordant with those from QR. Across both sexes, WMH burden was higher at older ages, accompanied by wider inter-percentile variation (Supplementary Figs. S4.F1–S4.F2 and Supplementary Tables S4.T1–S4.T2). Scans from participants with mild cognitive impairment (MCI) or Alzheimer’s disease (AD) dementia were scored against the cognitively unimpaired reference (Fig. 4b). The median burden percentile was 67 in individuals with AD, with 41.3% of scans exceeding the 75^th^ percentile, compared with the reference values of 50.0 and 25.0%, respectively. Individuals with MCI showed an intermediate distribution (median 58.0; 34.0% above the 75^th^ percentile). Among participants with AD, men had higher burden percentiles than women (median 70 versus 64), with 45% versus 38% exceeding the 75^th^ percentile. Percentile distributions were similar between sexes in MCI. Diagnostic differences were more pronounced at younger ages. The proportion of scans from people with AD exceeding the 75^th^ percentile was 54% at ages 45–55 years compared with 30% at ages 85–95 years, indicating greater overlap with the cognitively unimpaired reference at older ages (Fig. 4b).

Higher burden percentiles were associated with poorer memory, executive function, language and visuospatial performance, and with higher SPARE-BA, SPARE-AD, amyloid PET Centiloids, plasma p-tau217, hypertension, ASCVD risk and systolic blood pressure. Age- and sex-adjusted analyses showed similar cognitive and SPARE-AD associations, a stronger association with SPARE-BA, and generally attenuated molecular and vascular associations. Diagnosis-stratified analyses showed weaker cognitive associations in cognitively unimpaired participants than in AD, whereas SPARE-AD was associated with burden percentiles across all three diagnostic groups (r = 0.094, 0.125 and 0.219 for cognitively unimpaired, MCI and AD, respectively). Hypertension was associated with burden percentiles in cognitively unimpaired participants (r = 0.132, 95% CI 0.106–0.158) and MCI (r = 0.098, 95% CI 0.058–0.138), but not AD (r = 0.049, 95% CI −0.010 to 0.108). Correlations, sample sizes and sensitivity to cohort adjustment are detailed in (Supplementary Figs. S4.F3–S4.F4 and Supplementary Tables S4.T3–T4.)

The exported reference curves enabled individual WMH burden scoring. For a 72-year-old woman, the median reference WMH volume was 3.50 cm³, whereas volumes of 5.0 and 12.5 cm³ corresponded to the 63.0^th^ and 90.4^th^ percentiles, respectively. These examples illustrate how individual lesion burden can be evaluated against an age- and sex-specific normative distribution (Supplementary Fig. S4.F3).

### Population WMH frequency maps and spatial vulnerability

We generated population WMH frequency maps to characterize the spatial distribution of lesions across the full analytic sample. Voxelwise frequency, defined as the proportion of scans in which each location was labelled as lesional, was highly non-uniform, with greatest involvement in periventricular and central white matter and comparatively low frequencies across much of the remaining white matter. Across the full sample, the maximum voxelwise WMH frequency was 25.2%.

Age-stratified maps suggested higher WMH frequency and greater spatial extent at older ages (Fig. 5a). Peak voxelwise frequency was 15.2% before age 50, 29.6% during the 70s and 54.2% at age 90 years and older. At older ages, WMH involvement was more widespread beyond the periventricular zone into ventricle-distal white matter, indicating more diffuse spatial vulnerability. Sex-stratified maps showed no obvious differences in spatial distribution between men and women. One-scan-per-participant sensitivity maps were highly concordant with the full scan-level maps (Pearson r = 0.998, Spearman ρ = 0.969), with comparable peak frequencies (23.8% versus 25.2%), indicating that repeated scans did not materially alter the observed spatial pattern (Supplementary Fig. S5.F1 and Supplementary Table S5.T1).

**Figure 5.**
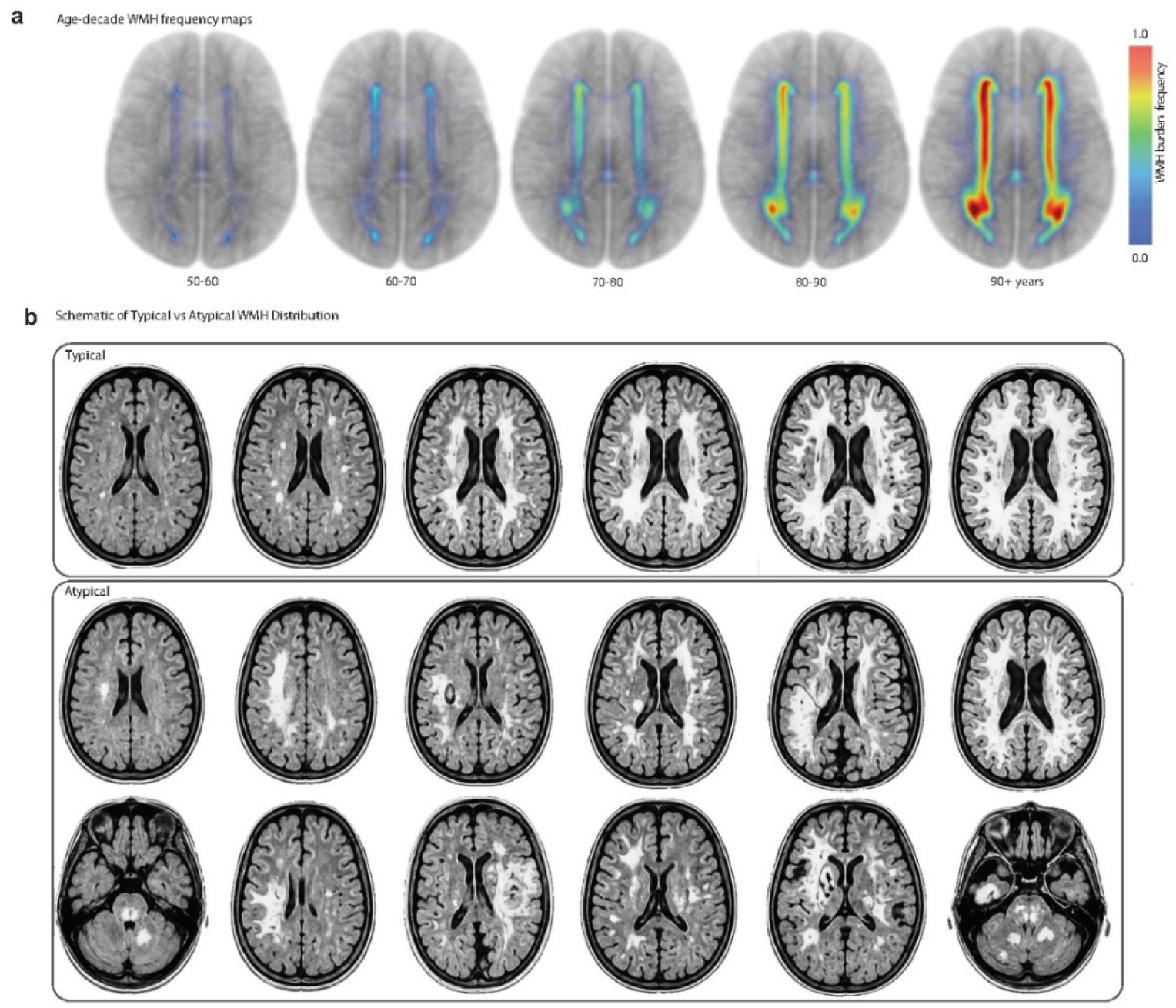
Age-related WMH frequency maps and schematic representation of typical and atypical WMH topography. a, WMH frequency maps from the full analytic sample, shown as glass-brain projections across successive age groups (50–59, 60–69, 70–79, 80–89 and ≥90 years). Voxelwise frequency denotes the proportion of scans within each age group containing WMH at a given location, illustrating age-related differences in lesion frequency and spatial distribution. b, Schematic representation of typical and atypical WMH spatial distributions. Typical patterns illustrate lesions concentrated in commonly affected periventricular regions, whereas atypical examples show asymmetric, unilateral or less commonly affected white-matter locations. Spatial typicality is defined relative to the age-, sex- and burden-conditioned population reference rather than anatomical location alone. The examples illustrate differences in lesion topography without assigning specific underlying disease mechanisms.

We then examined whether this spatial organization was preserved across individual cohorts. Cohort-specific maps showed broad consistency with the full-sample map, with spatial correlations ranging from r = 0.81 for WRAP to r = 0.99 for NACC, while retaining variation in the relative contribution of individual regions. Across cohorts, WMH distributions remained predominantly periventricular and frontal, with comparatively low temporal involvement. Regional composition nevertheless varied, with WRAP showing the most posterior-weighted distribution, including the highest mean occipital and PCA-territory fractions. Together, these spatial patterns provided the population reference for subsequent lesion-location scoring (Supplementary Table S5.T1).

### Voxelwise topographic reference modelling

Voxelwise topographic reference modelling (vTRM) quantified individual lesion-location typicality relative to age- and sex-matched frequency maps from cognitively unimpaired participants, conditioned on age and total WMH burden. The model explained 57% of the variance in mean lesion-location frequency, compared with 11% for a linear model of burden alone. Age interpolation provided continuous reference estimates across age bands. Because the score is a residual, three properties were established before use. Conditioning removed burden rather than leaving it in the residual: within every age band the residual was uncorrelated with log WMH volume, with no band exceeding r = 0.04. Within-participant reproducibility among participants with repeated scans gave an intraclass correlation of 0.79, against 0.86 for directly measured WMH volume, placing most residual variance between individuals rather than between repeat scans of the same person. Exact leave-one-out correction showed negligible self-inclusion bias in cognitively unimpaired scans (median absolute vTRM difference = 1.4 × 10⁻⁴; Supplementary Figs. S5.F2–S5.F4 and Supplementary Tables S5.T2–S5.T4).

Sex-stratified vTRM percentile charts were constructed from cognitively unimpaired scans (Fig. 4c). The fitted reference showed a U-shaped pattern of variability, with percentile bands narrowest around ages 46–47 years and wider at younger and older ages. Observed score dispersion supported widening at older ages, whereas the younger-age widening was less certain because of sparse observations and the quadratic model specification (Supplementary Fig. S5.F7 and Supplementary Table S5.T7).

vTRM percentiles differed across diagnostic groups (Fig. 4d). Median percentiles were 50 in cognitively unimpaired scans, 53.7 in MCI and 59.4 in AD (Mann–Whitney U, all pairwise P < 0.001). Among scans from individuals with AD, 34.9% and 16.5% exceeded the 75^th^ and 90^th^ normative percentiles, respectively, scans from individuals with MCI showed an intermediate distribution, with 28.8% exceeding the 75^th^ percentile.

Diagnostic differences were most pronounced at younger ages (Fig. 4d). Among scans from individuals with AD, 68.5% exceeded the 75^th^ vTRM percentile at ages 45–55 years, compared with 47% at 55–65 years, 34% at 65–75 years and 30–33% at 75–95 years. Thus, spatial atypicality was most strongly over-represented in AD at ages 45–55 years, with greater overlap with the cognitively unimpaired reference at older ages.

After adjustment for age, sex and total WMH volume, vTRM percentile was associated with higher SPARE-AD and SPARE-BA, poorer memory, executive function, language and visuospatial performance, and amyloid PET Centiloids and CSF Aβ42. No significant association was observed with the composite vascular burden score (partial r = 0.010, P = 0.35). PCA-territory WMH fraction was positively associated with amyloid PET Centiloids (partial r = 0.107, P < 10⁻⁴⁴) and CSF p-tau181 (partial r = 0.064, P < 10⁻⁵) after adjustment for the same covariates (Supplementary Fig. S5.F6 and Supplementary Table S5.T6).

### vTRM characterizes WMH heterogeneity

To assess whether lesion-location atypicality captured clinical and biological differences beyond total WMH burden, scans were stratified into four quadrants based on WMH burden and vTRM percentiles, using the 75th percentile as the threshold for each dimension. The AD-to-cognitively-unimpaired ratios were 0.14, 0.20, 0.27 and 0.48 in the low-burden/low-vTRM, low-burden/high-vTRM, high-burden/low-vTRM and high-burden/high-vTRM groups, respectively. The distribution differed significantly across quadrants (χ² = 849.7, P = 7.16 × 10⁻¹⁸⁴), with a 3.5-fold higher ratio in the high-burden/high-vTRM group than in the low-burden/low-vTRM group. Logistic regression further identified a positive burden × vTRM interaction on the log-odds scale (OR = 1.60, 95% CI 1.40–1.83, P = 1.28 × 10⁻¹¹; Supplementary Fig. S6.F1 and Supplementary Table S6.T1).

Among participants with AD, the four quadrants showed distinct clinical, biological and spatial profiles. Notably, among scans with high WMH burden, those with high vTRM percentiles showed greater amyloid burden, higher SPARE-AD, poorer memory performance and more ventricle-distal WMH, whereas those with low vTRM percentiles had the highest hypertension prevalence (73%) and predominantly periventricular WMH. The high-burden/high-vTRM group had the lowest median age among the four AD quadrants (73 years), with median memory performance of −0.87, SPARE-AD of 0.79, amyloid burden of 80 Centiloids and APOE-ε4 carrier prevalence of 60%. Periventricular WMH accounted for 45.7% of lesions in this group, compared with 77.8% in the low-burden/low-vTRM group; these compartments are defined by anatomical atlas and are independent of the frequency maps used to compute vTRM. Among the remaining quadrants, the low-burden/low-vTRM group was oldest (median 78 years), with the highest frequency of heart disease (28%) and diabetes (21%), whereas the low-burden/high-vTRM group had the highest frequency of stroke (16%) and temporal WMH fraction (8.8%). Cardiovascular frequency estimates were based on participants with available records. Comparable quadrant profiles were not observed among cognitively unimpaired participants (Supplementary Fig. S6.F2 and Supplementary Tables S6.T2–T4).

We next examined whether burden and vTRM showed distinct clinical and biological associations. In analyses adjusting vTRM percentiles for age, sex and log-transformed WMH volume, and burden percentiles for age and sex, vTRM showed stronger associations with SPARE-AD and SPARE-BA, whereas burden showed stronger associations with memory, executive function, and language. Associations with amyloid PET Centiloids also differed between the two measures. vTRM associations were less sensitive to cohort adjustment than burden-related cognitive associations. Correlation estimates, confidence intervals and sample sizes are provided in Supplementary Table S6.T2.

A one-scan-per-participant sensitivity analysis comprising 21,465 baseline scans with available vTRM scores supported the primary findings. AD-to-cognitively-unimpaired ratios across the four quadrants were 0.16, 0.24, 0.32 and 0.64, corresponding to a 4.0-fold contrast between the high-burden/high-vTRM and low-burden/low-vTRM groups. The burden × vTRM interaction remained positive (OR = 1.34, 95% CI 1.10–1.63, P = 0.003; Supplementary Table S6.T1a-d).

Together, these findings indicate that WMH burden and lesion-location atypicality capture partly distinct clinical and biological associations, with spatially atypical WMH providing information beyond total lesion volume.

### Scoring new lesion masks using the vTRM reference package

To enable application of the normative framework to new WMH scans, we assembled a vTRM reference package comprising twelve age- and sex-stratified WMH frequency maps derived from cognitively unimpaired participants, a fitted model for age- and burden-conditioning, normative percentile references and a single-scan scoring tool. The framework was integrated into Brain Lesion Analytics (https://brainlesionanalytics.org/), enabling automated vTRM scoring of WMH masks generated by the deployed 3D U-Net in approximately 45-60 seconds.

For each binary WMH mask in MNI152 2 mm space, the scoring tool uses age-interpolated, sex-specific frequency maps to estimate lesion-location typicality and assigns a vTRM percentile after conditioning on age and total WMH volume (Fig. 6). The output includes mean and median lesion-location frequency, the proportion of lesion voxels in rare (<1%) and common (>5%) locations, the vTRM raw score and percentile, and an interpretation relative to the cognitively unimpaired reference. The tool also provides descriptive lobar and lesion-depth labels, based on deviations in regional WMH composition from age- and burden-conditioned reference patterns.

**Figure 6.**
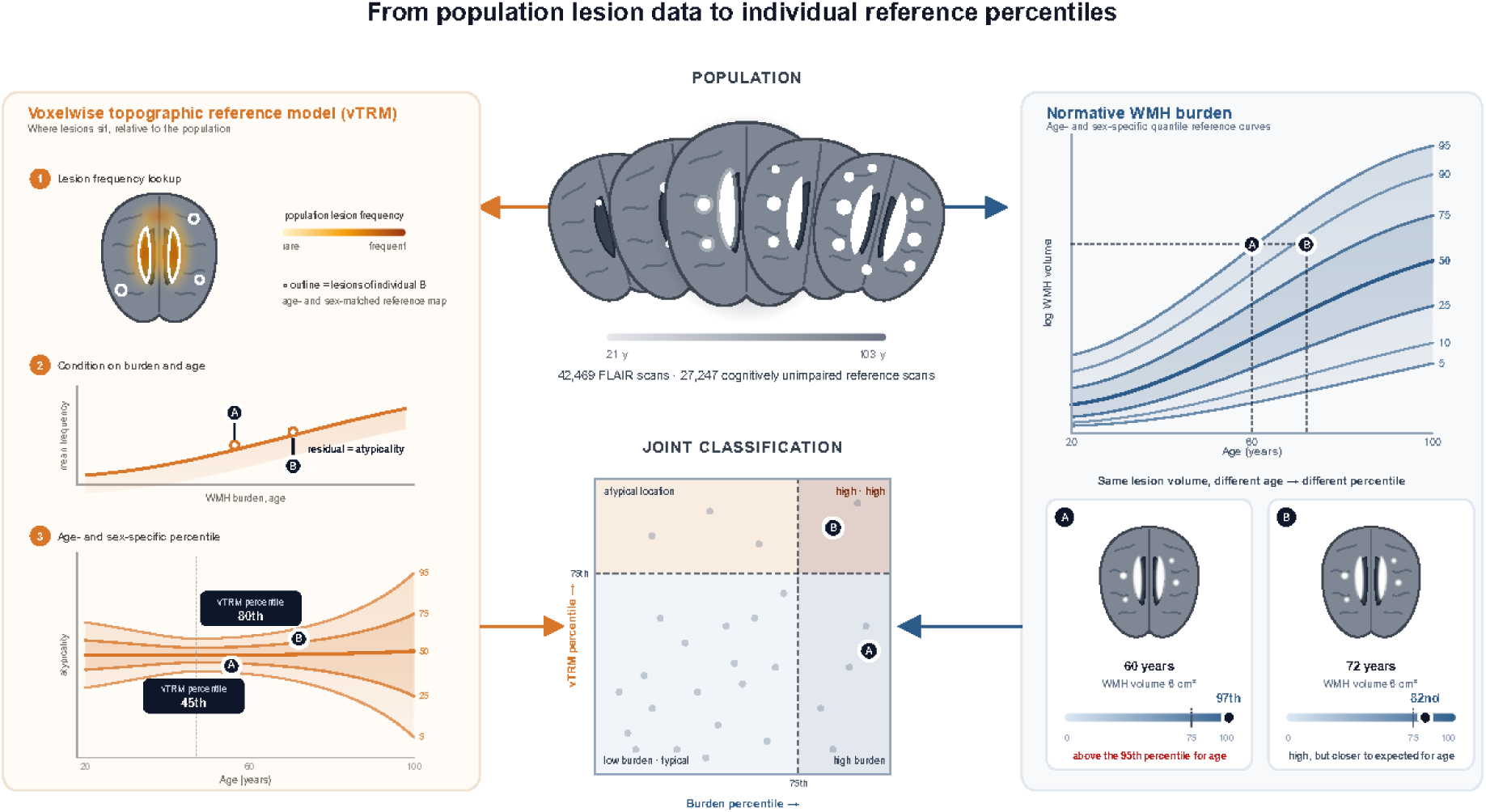
Schematic overview of WMH burden and vTRM percentile scoring for new WMH lesion masks. The normative reference framework scores an individual WMH mask on two independent dimensions, lesion burden and lesion location, each expressed as an age- and sex-specific percentile, using a large population sample (top centre; 42,469 FLAIR scans, of which 27,247 from cognitively unimpaired participants form the reference set, spanning 21–103 years). Normative WMH burden (right). Total WMH volume is evaluated against age- and sex-specific quantile reference curves (5^th^–95^th^ percentiles of log WMH volume as a function of age) to assign a burden percentile. Two example individuals illustrate the age dependence of this score: with an identical WMH volume of 6 cm³, individual A (60 years) falls at the 97^th^ percentile, above the 95^th^ percentile for age, whereas individual B (72 years) falls at the 82^nd^ percentile, high but closer to the volume expected for age. Voxelwise topographic reference model (vTRM; left). Lesion location is evaluated in three steps, shown for individual B. (1) Reference lesion frequencies are sampled at each lesion voxel from an age-interpolated, sex-specific WMH frequency map derived from cognitively unimpaired participants (heat map; white outlines mark the lesions of individual B); their mean provides an observed measure of lesion-location typicality. (2) The expected mean frequency is estimated with a conditioning model incorporating age and log-transformed WMH volume; the difference between observed and expected mean frequency defines the vTRM residual, a measure of spatial atypicality. (3) The residual is converted to an age- and sex-specific percentile, with higher values indicating greater spatial atypicality (individual B: 80th percentile). Joint classification (bottom centre). Burden and vTRM percentiles are combined in a two-dimensional chart characterising individuals by lesion quantity and spatial distribution. The 75th percentile on each axis defines four quadrants: low burden/typical location, high burden/typical location, low burden/atypical location and high burden/atypical location, where high denotes a percentile at or above 75. Individual B (82nd burden, 80th vTRM percentile) falls in the high burden/atypical location quadrant. Grey points represent reference individuals. Orange and blue arrows indicate the flow from population data through the topographic and burden branches, respectively, to the joint classification.

To assess applicability without study-specific cohort information, we compared cohort-free scoring with the original cohort-adjusted reference. Percentiles generally differed by only a few points, although discrepancies were larger below age 50, where cognitively unimpaired reference data were sparse. These findings support the feasibility of applying the reference framework to new lesion masks without cohort-specific adjustment or model refitting (Supplementary Fig. S5.F5 and Supplementary Table S5.T5 and Supplementary Table S7.T1).

## Discussion

Here we present voxelwise topographic reference modelling (vTRM), a population-scale framework for mapping WMH spatial vulnerability and quantifying individual lesion-location typicality using 42,469 standardized FLAIR WMH lesion masks. The central advance is to extend normative WMH assessment beyond lesion quantity to lesion topography. Burden percentiles quantify whether total WMH volume is high or low for age and sex, whereas vTRM quantifies whether that burden occurs in locations that are typical or atypical given age, sex and total WMH burden. These two dimensions define a quadrant-based classification framework, an approach used in cardiology, critical care and neuroimaging to distinguish physiological or biological profiles [^19^,^20^,^21^,^22^,^23^]. This framework addresses an important gap: individuals with similar lesion volumes can exhibit markedly different spatial patterns, potentially implicating distinct vascular, neurodegenerative or mixed processes [^24–27^]. The limitation of burden alone becomes particularly apparent with ageing; among individuals with AD dementia, the proportion exceeding the normative 75^th^ burden percentile was 54% at ages 45–55 but was substantially lower, 30%, at ages 85–95, suggesting increasing overlap between normative and disease-related burden in older age groups. vTRM addresses this limitation by defining lesion-location typicality empirically from cognitively unimpaired population frequency maps, conditioned on age and burden to isolate spatial atypicality from volume-driven effects. This extends percentile-based brain-chart principles from how much WMH is present to where it occurs, providing an additional reference axis for resolving heterogeneity obscured by global lesion volume alone.

First, the voxelwise frequency maps establish a population reference for WMH spatial vulnerability, suggesting that ageing is associated not only with greater lesion burden but also with systematic differences in lesion topography. WMH at younger ages and lower burden were predominantly periventricular, whereas older individuals exhibited greater ventricle-distal, frontal, parietal and MCA-territory involvement. Regional growth-rate models suggested measurable growth first in periventricular white matter before age 50, followed by frontal and MCA territories in the sixth decade and parietal and ventricle-distal regions in the seventh decade. After age 70, estimated MCA-territory growth rates exceeded those in periventricular white matter. These findings argue against uniform lesion accumulation or simple radial expansion from the ventricles, instead supporting age-related differences in WMH distribution across ventricular-distance, lobar and vascular-territory axes. The earliest estimated regional onsets, from periventricular white matter in early adulthood to posterior compartments around midlife, preceded the age range covered by concurrent vascular, molecular and diagnostic measures, leaving the biological context of WMH initiation unresolved. Later redistribution of growth away from the ventricular surface coincided with ages at which vascular risk was common and indicators of amyloid and tau pathophysiology were higher in the contributing cohorts, suggesting a more complex cerebrovascular and neurodegenerative background rather than a uniform continuation of earlier periventricular processes. These relationships are temporal rather than mechanistic, and differences in cohort composition preclude direct comparisons of absolute prevalence or relative contributions across measures. Importantly, absolute burden and fractional topography were not equivalent. PCA-territory WMH burden was higher at older ages in absolute terms but proportionally lower, alongside greater frontal, ventricle-distal and MCA involvement. Nevertheless, PCA-territory topography was associated with amyloid PET and plasma p-tau217 after adjustment for age and total WMH volume. These findings suggest that age-related redistribution and AD-related topographic vulnerability represent distinct but convergent signals: broader ventricle-distal and MCA involvement may reflect ageing-related vascular injury, whereas relatively greater posterior/PCA involvement may be relevant to AD-related pathophysiology and neurodegeneration [28,29].

Second, individualized vTRM percentiles provide information beyond conventional lesion burden. Among scans from individuals with clinical AD, 35% exceeded the normative 75^th^ vTRM percentile, and this higher proportion was observed in older age groups, though it was less pronounced in older age groups. When burden and vTRM were considered jointly, the AD-to-cognitively unimpaired ratio showed a 3.5-fold difference between the low-burden/low-vTRM and high-burden/high-vTRM groups. These findings establish spatial typicality as an additional dimension to lesion volume, with their combination providing greater separation of disease-related heterogeneity than either measure alone.

Third, diagnostic-stage analyses support a dimensional rather than categorical interpretation of spatial atypicality. Median vTRM percentiles were higher across cognitively unimpaired, MCI and AD dementia groups (50, 54 and 55, respectively), suggesting graded differences in WMH topography rather than a discrete disease-specific pattern. These cross-sectional associations do not establish within-person progression but suggest greater deviation from normative WMH organization with increasing clinical severity [^30–32^]. Thus, vTRM represents a dimensional marker of WMH heterogeneity rather than a diagnostic classifier, potentially supporting biologically informed stratification and endophenotyping.

Fourth, vTRM captured biological information beyond total lesion burden. After adjustment for age, sex and WMH volume, vTRM was associated with neurodegeneration (SPARE-AD and SPARE-BA) and with amyloid PET Centiloids, but not vascular risk. These findings suggest that lesion quantity and topography capture partially distinct biological dimensions, with burden more closely reflecting vascular and age-related injury and spatial atypicality signaling impact on or consequences of AD-associated neurodegeneration and amyloid pathology [^33^]. AD-specific quadrant analyses reinforced this distinction: individuals with high burden but low vTRM had the highest hypertension rate and predominantly periventricular WMH, whereas those with high burden and high vTRM were younger, with greater amyloid burden and APOE-ε4 frequency but fewer vascular comorbidities. A significant burden × vTRM interaction indicated that their combination was associated with greater odds of AD than predicted by either dimension alone, supporting distinct quadrant phenotypes rather than independent gradients. Matched scans further illustrated that similar WMH volumes can have markedly different spatial typicality, while even very high burden can follow an expected periventricular distribution. These findings support the two-axis framework for characterizing biologically heterogeneous WMH correlates within AD.

The translational value of vTRM lies in combining lesion burden and spatial typicality to characterize WMH phenotypes, potentially identifying endophenotypes relevant to AD that are obscured by burden alone [^34^]. The framework can be applied directly to external MNI-space lesion masks, requiring only a lesion mask, age, and sex to generate an individualized spatial-typicality percentile. By adding an interpretable spatial dimension to established WMH burden measures, vTRM provides a practical framework for investigating WMH heterogeneity in brain ageing and dementia. Cross-cohort variation presents a challenge for normative WMH modelling because differences in lesion burden may reflect both technical factors and genuine differences in underlying pathology. Cohort effects predominantly shifted reference burden levels rather than age-related trajectory shapes, although leave-one-cohort-out deviations were substantial in some studies (Supplementary Figs. S2.F5 and S2.F8). In contrast, voxelwise WMH frequency maps showed consistent spatial patterns across cohorts (r = 0.81–0.99; Supplementary Fig. S5.F1), and cohort-free vTRM percentiles closely agreed with cohort-adjusted estimates (r = 0.983; median absolute difference, 2.30 percentile points; Supplementary Fig. S5.F5). These findings demonstrate the consistency of the spatial reference across contributing cohorts and support its application without cohort-specific model refitting, while independent-site validation remains an important next step.

The etiology of WMH in ageing and AD remains debated [^12,35,36^]. Although WMH are widely attributed to cerebral small-vessel disease in ageing [^3,37,38^], their biological origins in AD are less clear. Some propose that AD-related WMH arise downstream of neurodegeneration through Wallerian degeneration [^39^], whereas evidence from temporal sequencing, spatial distribution, biomarker associations and preclinical models [^40,41^] supports vascular and/or inflammatory contributions potentially linked to AD risk and pathophysiology [^42^]. The AD-related WMH topography observed here is consistent with previous associations between regional WMH distribution and AD risk [^43^], suggesting a potential endogenous vascular or inflammatory contribution to disease susceptibility. Moreover, many AD genetic risk loci are expressed in the vasculature [^44,44^]. Regardless of the underlying mechanisms, our findings provide a spatial reference for interpreting WMH in normative and pathological brain ageing.

Several methodological considerations are relevant to interpreting these findings and guiding future refinement. The vTRM score summarizes lesion-location typicality as a single scalar without indicating the direction of atypicality. Although regional compartment analyses separately characterize dominant lobar involvement and lesion depth, future work should assess whether integrating these features improves biological or prognostic interpretation. Age-banded normative frequency maps also introduce discrete transitions that continuous age-conditioned models may resolve. Despite standardized processing, residual variability may reflect biological heterogeneity or differences in lesion segmentation and registration [^45^]. Subsampling analyses demonstrated that the normative reference curves were largely converged at the available sample size, particularly across well-sampled ages, although larger datasets may further improve precision at age extremes and upper percentiles. Reference estimates may nevertheless vary with cohort composition and population representation [^14,46^], underscoring the importance of evaluating generalizability across populations. Although WMH burden differences are often observed across race and ethnicity groups, race and ethnicity was not included as a covariate in the normative models, consistent with recommendations against routine adjustment for race or ethnicity without scientific justification [^47^]. Previous normative modelling research showed that sociodemographic variation in brain structure can remain detectable [^48^], supporting the importance of examining mediators of population differences rather than automatically adjusting them away. The small association with CSF Aβ42 requires replication, and the cross-sectional framework does not establish longitudinal progression. Diagnostic analyses provide secondary validation, but percentile thresholds require external and prospective validation before clinical use.

The reference models also included repeated scans, potentially overweighting participants with longer follow-up. However, repeat frequency was negligibly associated with age or WMH burden, and AD participants contributed fewer scans per person than cognitively unimpaired participants. Given the highly right-skewed WMH distribution, repeated cognitively unimpaired observations were concentrated near the low-burden floor, limiting their influence on upper-tail quantiles [^49,50^]. Baseline-only models produced comparable or stronger associations; the AD-to-cognitively unimpaired ratio across joint phenotypes was 4.0-fold versus 3.5-fold using all scans, with a significant burden × vTRM interaction in both analyses. Thus, repeated observations did not drive the findings and may have yielded more conservative estimates (Supplementary Fig. S2.F10 and Supplementary Table S2.T10).

In summary, vTRM extends normative brain-chart principles to WMH topography, providing an individualized measure of spatial typicality relative to age-, sex- and burden-matched population references. By distinguishing lesion quantity from expected or atypical spatial patterns, this framework reveals WMH heterogeneity obscured by global volume measures and supports more biologically informed characterization of brain ageing and dementia.

## Methods

### Ethics

This study used de-identified imaging and metadata from previously collected cohort studies. All participants provided informed consent, and primary data collection was conducted under Institutional Review Board-approved protocols and applicable data-use agreements at the contributing institutions. Participant compensation, when provided, was administered by the original cohort studies.

### Software reporting

Statistical analyses and visualizations were performed in Python (v3.11.15) using pandas (v3.0.2), NumPy (v1.26.4), seaborn (v0.13.0), matplotlib (v3.10.8), pyGAM (v0.12.0), SciPy (v1.16.3), statsmodels (v0.14.6), scikit-learn (v1.8.0), and NiBabel (v5.4.2). GAMLSS sensitivity models were fitted in R (v4.5.3) using the gamlss package (v5.5.0) via rpy2 (v3.6.7). Three-dimensional glass-brain visualizations of WMH frequency maps were generated using Vedo (v2025.5.4). Software used for MRI preprocessing and processing is detailed in the MRI processing pipeline section.

### Data

We compiled Fluid-Attenuated Inversion Recovery (FLAIR) MRI scans and harmonized scan-level metadata from the Alzheimer’s Disease Sequencing Project Phenotype Harmonization Consortium (ADSP-PHC) Release 2 (https://vmacdata.org/adsp-phc). Data originated from twelve contributing cohorts: Anti-Amyloid Treatment in Asymptomatic Alzheimer’s (A4) Study and Longitudinal Evaluation of Amyloid Risk and Neurodegeneration (LEARN) Study [^51^], Alzheimer’s Disease Neuroimaging Initiative (ADNI) (https://adni.loni.usc.edu/), National Alzheimer’s Coordinating Center (NACC) and the Standardized Centralized Alzheimer’s & Related Dementias Neuroimaging (SCAN), which include data contributed by more than 30 federally funded Alzheimer’s Disease Research Centers (ADRCs) across the United States [^52,53^], Health and Aging Brain Study-Health Disparities (HABS-HD) [^54^], Korean Brain Aging Study for the Early Diagnosis and Prediction of Alzheimer’s Disease (KBASE) [^55^], Religious Orders Study/Rush Memory and Aging Project/Minority Aging Research Study (ROS/MAP/MARS: www.radc.rush.edu) [^56,57^], Knight Alzheimer Disease Research Center (Knight-ADRC) (https://knightadrc.wustl.edu/), Washington Heights and Inwood Columbia Aging Project (WHICAP) [^58^], and Wisconsin Registry for Alzheimer’s Prevention (WRAP) [^59,60^]. Participants ranged from 21 to 103 years of age and included cognitively unimpaired individuals and those with mild cognitive impairment (MCI) or Alzheimer’s disease (AD) dementia.

ADSP-PHC provided harmonized demographic, imaging, diagnostic, cognitive, vascular, molecular biomarker, genetic, and MRI-derived neurodegeneration data. Cognitive domain scores were harmonized across cohorts using domain-specific models [^61^]. Data tables and dictionaries were integrated to link FLAIR scans with demographic and clinical metadata. Following FLAIR processing, WMH segmentation, spatial standardization, and image-level quality control, the final analytic sample comprised 42,469 WMH lesion masks with complete age and sex information.

### MRI processing and data selection

All FLAIR images underwent visual quality control to exclude unreadable scans, imaging artifacts, major orientation errors and incomplete brain coverage. Images passing quality control were processed using the Automatic Segmentation of White Matter Hyperintensities (ASWMH) pipeline, as previously described [^62^]. The pipeline performs image standardization, preprocessing, white matter segmentation and automated WMH labeling, with lesion masks transformed to MNI152 2 mm standard space. Segmentations were visually reviewed for lesion plausibility, image–mask alignment and segmentation artifacts. Scans passing imaging and segmentation quality control were retained if valid lesion masks and corresponding demographic metadata were available. No scans were excluded due to missing age or sex information. For vTRM analyses, 1,060 scans (2.5%) with minimal WMH burden were excluded because reliable spatial alignment to MNI152 space could not be achieved without manual correction. These scans remained eligible for analyses not requiring vTRM.

Total WMH volume was calculated from lesion voxel counts and voxel volume and expressed in cubic centimeters. Age was treated as a continuous variable, and sex was encoded as a binary variable according to the contributing study. Scans from cognitively unimpaired participants were used to establish normative WMH burden and voxelwise topographic reference models (vTRM). Scans from participants with MCI or AD dementia were evaluated against these references. The primary analysis included 42,469 scans from 22,084 participants, of which 27,247 scans from cognitively unimpaired individuals were used to establish the normative models. Sensitivity to repeated observations was assessed by refitting models and downstream analyses using one scan per participant (n = 13,303 for normative models), selecting baseline or otherwise the earliest FLAIR scan.

The reference framework is publicly available at https://brainlesionanalytics.org. The web application provides demographic summaries, age- and sex-specific WMH burden and regional reference charts, three-dimensional visualizations of age-related spatial WMH patterns, burden and vTRM percentile scoring, quadrant visualizations, and percentile reports for uploaded WMH masks. The platform also includes ADSP-Lite (beta), a deployable 3D U-Net model for automated WMH segmentation trained on ASWMH-derived lesion masks from ADSP-PHC imaging.

### Normative modelling of WMH volume

Age- and sex-specific normative reference curves for total and regional WMH burden were established using scans from cognitively unimpaired participants. WMH volumes were right-skewed and included near-zero values and were therefore log-transformed:

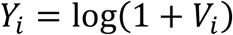

where *Y_i_* is log-transformed WMH volume and *V_i_* is WMH volume in cubic centimeters. This transformation accommodates zero values while preserving the ordering of WMH volumes. Sex-stratified quantile regression (QR) was used to estimate age-dependent reference percentiles without assuming a parametric outcome distribution. Models were fitted for total WMH burden and periventricular, ventricle-distal, lobar and arterial-territory compartments. Conditional quantiles were modelled as:

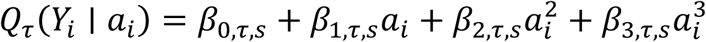

where *a_i_* denotes age, *s* denotes sex, and *β* represents sex- and quantile-specific coefficients. Reference curves were estimated at the 5^th^, 10^th^, 25^th^, 50^th^, 75^th^ and 95^th^ percentiles.

Primary models excluded cohort indicators to provide pooled reference curves independent of contributing study. Cohort-adjusted sensitivity models were also examined. Two alternative modelling frameworks were evaluated. Gaussian-family generalized additive models (GAMs), with age-dependent mean and variance, were fitted for total WMH burden. Regional reference curves were also estimated using generalized additive models for location, scale and shape (GAMLSS). Following Bethlehem et al. [^14^], generalized gamma distributions were fitted separately by sex using fractional polynomials in age, with the shape parameter specified as a global offset. Exact zero volumes were replaced with half the voxel volume to satisfy the distribution’s positive-support requirement. Model selection used the Bayesian information criterion, first optimizing the location parameter with scale restricted to a single term, followed by scale optimization conditional on the selected location specification. GAMLSS models were fitted with and without cohort indicators.

### Regional WMH phenotyping

To summarize WMH spatial distribution across anatomical compartments, lesion burden was extracted from periventricular, ventricle-distal, lobar and arterial-territory masks in MNI152 2 mm space. A binary ventricular mask was constructed from the lateral, inferior lateral, third and fourth ventricular labels of the MNI-space FreeSurfer-derived parcellation (WMPARC labels 4, 5, 14, 15, 43 and 44). Euclidean distance from this mask was computed in millimeters, accounting for 2 mm voxel spacing. Lesion voxels were classified as periventricular (greater than 0 and up to 10 mm) or ventricle-distal (greater than 10 mm), the latter extending from the outer periventricular boundary toward the cortical ribbon. For each scan and anatomical region, lesion volume, lesion presence and the region’s fraction of total WMH volume were computed:

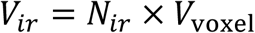

where *V_ir_* is WMH volume for scan *i* in region *r*, *N_ir_* is the number of lesion voxels in region *r*, and *V*_voxel_ is voxel volume.

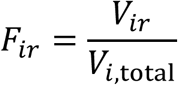

where *F_ir_* is the fraction of total WMH volume in region *r*, *V_ir_* is regional WMH volume, and *V_i_*_,total_ is total WMH volume. Regional WMH volume was defined as the number of lesion voxels in the region multiplied by voxel volume. Regional WMH fraction was defined as regional WMH volume divided by total WMH volume, where total WMH volume was the sum of WMH volume across the whole lesion mask. Regional fractions quantify WMH spatial composition, allowing regional redistribution to be analyzed separately from age-related differences in total lesion burden. Absolute regional burden captured how much lesion volume was present in a region, whereas fractional topography captured how each scan’s total WMH burden was distributed across regions. Summary measures included periventricular 0–10 mm fraction, ventricle-distal fraction, anterior fraction, posterior fraction, and PCA-territory fraction. Regional enrichment was calculated as the ratio of each compartment’s pooled WMH volume fraction to its fraction of total white matter volume, with compartment sizes derived from atlas voxel counts in MNI152 2 mm space. Raw ratio measures with unstable denominators were not used for primary inference; fraction-based measures were used for validation analyses.

### WMH accumulation milestones

To characterize age-related differences in regional WMH burden, normative slopes were derived from sex-stratified median QR curves. For each anatomical compartment, the slope was calculated as the derivative of median log-transformed WMH volume with respect to log age:

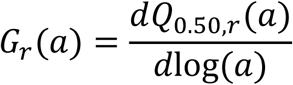

where *G_r_*(*a*) denotes the age-related slope (growth-rate analogue) for region *r*, and *Q*_0.50,*r*_(*a*) is the fitted median reference curve.

For this analysis, median curves were refitted using sex-stratified QR with linear and quadratic age terms, without cohort indicators, and evaluated between 20 and 100 years. The resulting slope curves were smoothed using a Gaussian kernel (σ = 8). The slopes represent changes in fitted log-transformed WMH burden per unit difference in log age, rather than annual changes in lesion volume. Slopes were averaged across sexes and evaluated at the midpoints of six age strata (<50, 50–59, 60–69, 70–79, 80–89 and ≥90 years). Ninety-five percent confidence intervals were estimated using 200 participant-clustered bootstrap replicates, with QR fitting, differentiation and smoothing repeated within each replicate.

Milestones were defined as the ages at which the smoothed, model-estimated slope curves first crossed zero. Two additional crossover milestones were defined as the ages at which MCA-territory and ventricle-distal slopes first exceeded the periventricular slope. Milestone ages were estimated by linear interpolation, with confidence intervals obtained from the same bootstrap procedure. Robustness to contributing studies was assessed through leave-one-study-out sensitivity analyses. Estimates were bounded by the evaluated age range when zero crossings occurred outside the modelled interval. Concurrent vascular, molecular and diagnostic measures were evaluated at the corresponding milestone ages to provide biological and clinical context.

Five indices were derived to characterize age-related differences in WMH spatial composition. The anterior-to-posterior ratio was defined as frontal plus parietal WMH volume divided by occipital plus temporal volume; the periventricular-to-distal ratio as periventricular 0–10 mm volume divided by ventricle-distal volume; and the MCA-to-PCA ratio as MCA-territory volume divided by PCA-territory volume. A small constant was added to denominators to avoid division by zero. Spatial concentration was quantified using the Herfindahl–Hirschman index (HHI), adapted from economics for biological applications [^63,64^], calculated as the sum of squared volume fractions across seven anatomical compartments (periventricular, ventricle-distal, frontal, parietal, occipital, temporal and MCA territory). Because these compartments overlap, HHI was interpreted as a composite spatial concentration measure rather than concentration across mutually exclusive regions. The fifth index, MCA-territory fraction, was calculated as MCA-territory WMH volume divided by total WMH volume. Age-dependent reference curves for each index were fitted using median QR with a fourth-degree polynomial in age, evaluated at 200 equally spaced ages between 21 and 100 years, with confidence intervals from 200 bootstrap replicates.

### Burden percentile scoring across diagnostic groups

Age- and sex-specific WMH burden percentile charts were established using scans from cognitively unimpaired participants. Total WMH volume was transformed as log(V + 10⁻⁶), and sex-stratified QR models were fitted at the 5^th^, 10^th^, 25^th^, 50^th^, 75^th^, 90^th^ and 95^th^ percentiles using linear and quadratic age terms:

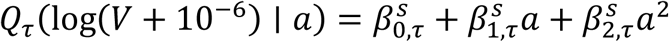

where *V* is total WMH volume in cm³, *a* is age, ***s*** denotes sex, and *τ* ∈ {0.05,0.10,0.25,0.50,0.75,0.90,0.95}. Models excluded cohort indicators to provide pooled reference percentiles independent of contributing study. Individual burden percentiles were assigned by interpolating observed log-transformed WMH volumes across the corresponding sex-specific conditional quantiles. These scoring models were distinct from the cubic QR models used to characterize normative WMH reference curves.

For comparison, a Gaussian-family generalized additive model (GAM) was fitted with the mean and variance modelled as smooth functions of age, and percentiles derived from the conditional Gaussian distribution. The spline basis dimensions were set to eight for the mean and four for the variance. QR and GAM percentile assignments were compared.

Scans from those with MCI or AD dementia were scored against the normative reference calculated from cognitively unimpaired individuals. The proportion exceeding the 75th percentile was calculated overall and within five age bands (45–55, 55–65, 65–75, 75–85 and 85–95 years). Model coefficients were exported as a portable scoring tool that estimates individual burden percentiles from age, sex and total WMH volume using the corresponding sex-specific quantile curves. The same scoring procedure was applied to internal and external scans without cohort-specific adjustment. Individual scoring examples and associations with cognitive, biomarker and vascular measures were examined, along with sensitivity to between-study variation.

### Clinical and biological context of WMH milestones

To contextualize model-estimated WMH milestone ages, age-specific prevalence was estimated for seven vascular, molecular and diagnostic measures: hypertension, diabetes, and stroke from cardiovascular records; amyloid positivity from PET; tau positivity defined as Braak stage ≥3; and MCI and AD dementia from diagnostic records. Cardiovascular and diagnostic measures were matched within one year of the scan to avoid attributing subsequent diagnoses to earlier scan ages. Participants with non-AD dementia or other cognitive diagnoses were retained in the denominators for diagnostic frequency estimates. Each measure was modelled as a binary outcome using generalized estimating equations (GEE) with a binomial family, logit link and exchangeable working correlation clustered by participant. Models included a natural cubic spline for age, sex and contributing study with sum-to-zero coding. This provided population-average age-specific probabilities with robust covariance estimation. Spline complexity was evaluated separately for each measure using three, four and five degrees of freedom and the uncorrected quasi-likelihood under the independence model criterion (QICu). The smallest number of degrees of freedom within two QICu points of the minimum was selected.

Predicted probabilities were standardized by evaluating the fitted model for each contributing study and sex at each age, transforming predictions to the probability scale, and averaging equally across studies and according to the measure-specific sample sex distribution. To account for differences in study age coverage, primary estimates were standardized over a fixed set of studies with adequate coverage at the relevant milestone ages; standardization using locally varying study sets was evaluated as a sensitivity analysis. Estimates were suppressed when fewer than 50 participants were within five years of the target age, fewer than two studies contributed, or the target age was outside the observed range. Ninety-five percent confidence intervals were obtained from 2,000 draws from the robust coefficient covariance matrix, propagated through the same standardization procedure. Study sets, excluded zero-event studies and sensitivity analyses were documented.

### Population WMH frequency maps

Population WMH frequency maps were generated by aggregating binary lesion masks in MNI152 2 mm standard space. Voxelwise WMH frequency was defined as the proportion of valid scans containing a lesion at each voxel. Maps were generated at the scan level, consistent with the reference framework. Frequency maps were generated for the full analytic sample and stratified by age, sex and cohort. Six age strata were defined: <50, 50–59, 60–69, 70–79, 80–89 and ≥90 years. Cohort-specific maps were used for quality control and assessment of spatial consistency across datasets. Consistency was quantified using voxelwise correlations between cohort-specific and full-sample maps, and between sex-specific and corresponding age-stratified maps, restricted to voxels containing lesions in at least one scan.

Sensitivity maps were generated using one scan per participant and compared with the full-sample maps.

### Voxelwise topographic reference modelling

Voxelwise topographic reference modelling (vTRM) quantifies lesion-location typicality by comparing individual WMH masks with population frequency maps derived from cognitively unimpaired participants. Lesions in locations with higher reference frequencies are considered spatially typical, whereas lesions in less frequently affected locations are considered atypical. The procedure comprises three steps: frequency lookup, burden-and-age conditioning and percentile assignment.

Step 1: Frequency lookup. For each sex and the six age strata defined above, binary WMH masks from cognitively unimpaired scans were averaged voxelwise to generate twelve reference frequency maps. To obtain age-specific references without discontinuities at age-band boundaries, maps were linearly interpolated between the two reference maps whose median ages bracketed the individual’s age. Scans outside the range of reference median ages were assigned the nearest reference map. The influence of interpolation was assessed. Potential self-inclusion bias in the reference maps was evaluated using exact leave-one-out correction. Mean lesion-location frequency was calculated by averaging the interpolated reference frequencies across all lesion voxels:

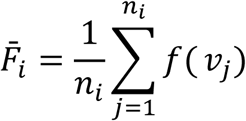

where 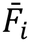 is the mean lesion-location frequency for scan *i*, *n_i_* is the number of lesion voxels, and *f*(*v_j_*) is the interpolated age- and sex-specific reference frequency at voxel *v_j_*. All scans included in vTRM scoring contained at least one lesion voxel in MNI152 space, requiring no special handling of empty lesion masks.

Step 2: Burden-and-age conditioning. Because mean lesion-location frequency depends on total WMH burden, the expected frequency was modelled as a function of WMH volume and age using tensor-product cubic B-spline regression fitted to cognitively unimpaired scans:

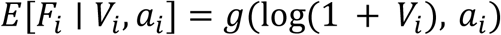

where *g* is a tensor-product B-spline function of four knots and degree 3 in each variable, *V_i_* is total WMH volume in cm³, and *a_i_* is age in years. Separate spline bases for log-transformed volume and age, together with their pairwise interactions, allowed the association between burden and mean frequency to vary with age. Comparisons with an additive specification were performed. The vTRM score was defined as the residual:

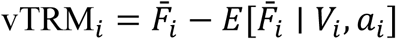

Negative residuals indicate less frequent lesion locations than expected for age and WMH burden, whereas positive residuals indicate more frequent locations. Because the vTRM score is a residual, it carries both lesion-location information and any variance not absorbed by the conditioning model. Three properties were therefore established before the score was used. Independence from burden was verified by testing the residual against log WMH volume within each age band, under both the additive and tensor specifications. Within-individual reproducibility was quantified by intraclass correlation among participants cognitively unimpaired at every visit with two or more scans, with directly measured WMH volume included as a benchmark. Independent spatial validity was assessed by comparing the regional composition of scans matched on burden but differing in vTRM, using compartments defined by anatomical atlas rather than by the reference frequency maps (Supplementary Table S6.T4).

Step 3: Percentile assignment. Sex-stratified vTRM percentile charts were fitted in cognitively unimpaired scans using QR at the 5^th^, 10^th^, 25^th^, 50^th^, 75^th^, 90^th^ and 95^th^ percentiles:

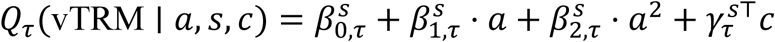

where *τ* ∈ {0.05, 0.10, 0.25, 0.50, 0.75, 0.90, 0.95}, *a* is age, *s* denotes sex, and c is a vector of cohort indicators with sum to zero coding, yielding an unweighted cohort-average reference. Individual typicality percentiles *P_i_* were obtained by interpolating observed vTRM scores across the corresponding sex-specific conditional quantiles. Percentiles were inverted so that higher values indicated greater spatial atypicality:

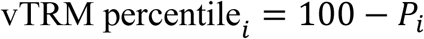

Thus, a vTRM percentile of 75 indicates greater spatial atypicality than 75% of the age- and sex-matched cognitively unimpaired reference. External scans were scored with cohort terms set to zero, corresponding to the unweighted cohort-average reference. QR provided age-conditional percentile estimates rather than age-independent global rankings. Age-related variability in vTRM scores was evaluated using the widths of the fitted 5th–95th and 25th–75th percentile bands and observed interquartile ranges calculated in sliding 10-year age windows. Fitted and empirical variability estimates were compared to assess the extent to which age-dependent dispersion reflected the quadratic reference model.

### Clinical, biological, and diagnostic analysis

Partial correlations were used to assess associations of vTRM and burden percentiles with cognitive, vascular, neurodegeneration, and molecular biomarker measures. For vTRM percentile, both the percentile and each outcome were residualized against age, sex and log-transformed total WMH volume using linear regression, followed by Pearson correlation between residuals. Burden percentile correlations were adjusted for age and sex only, because additional adjustment for WMH volume would remove the burden-related variation being assessed. Outcomes included harmonized cognitive domain scores (memory, executive function, language and visuospatial function), vascular burden (harmonized cardiovascular principal component), MRI-derived neurodegeneration measures (SPARE-AD, reflecting AD-like patterns of brain atrophy, and SPARE-BA, reflecting age-related patterns of brain atrophy)[^65,66^], and molecular AD biomarkers (amyloid PET Centiloids, plasma p-tau217, CSF p-tau181 and CSF Aβ42). Partial correlations between PCA-territory WMH fraction and molecular biomarkers were calculated using the same adjustment. Joint burden and spatial atypicality were assessed by classifying scans into four groups according to burden and vTRM percentiles, using the 75^th^ percentile as the upper-quartile cut point on both axes: low/low, high/low, low/high and high/high group. The AD-to-cognitively-unimpaired ratio was compared across groups using a chi-squared test. An interaction between high burden and high spatial atypicality was assessed by logistic regression:

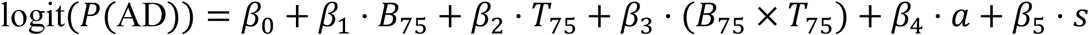

where *B*_75_ indicates burden percentile ≥ 75^th^ and *T*_75_ indicates vTRM percentile ≥ 75^th,^ *a* is age and *s* denotes sex. The interaction coefficient *β*_3_ was used to assess departure from additivity on the log-odds scale.

Quadrants were compared using Kruskal–Wallis tests for age, WMH volume, SPARE-AD, SPARE-BA, the four cognitive domain scores, amyloid PET Centiloids, plasma p-tau217, BMI, systolic and diastolic blood pressure, cardiovascular principal component and education. Chi-squared tests were used for hypertension, diabetes, heart disease, stroke, antihypertensive medication use, smoking status and APOE-ε4 carrier status. Analyses were conducted separately in AD and cognitively unimpaired scans to distinguish disease-related from normative differences. Regional WMH fractions across periventricular, ventricle-distal, four lobar and three arterial-territory compartments were also compared across quadrants.

## Reporting Summary

Further information on research design is available in the Reporting Summary linked to this article.

## Data availability

The imaging, demographic, clinical and biomarker data analysed in this study were obtained from 12 contributing cohorts through the Alzheimer’s Disease Sequencing Project Phenotype Harmonization Consortium (ADSP-PHC) Release 2 (https://adseq.org/). Access to participant-level data is governed by the respective cohort data-use agreements and institutional policies. These data may be requested from the original study investigators or designated data repositories, subject to applicable access requirements.

Data from the Alzheimer’s Disease Neuroimaging Initiative (ADNI) are available upon request from https://adni.loni.usc.edu/.

Data from the National Alzheimer’s Coordinating Center (NACC) and Standardized Centralized Alzheimer’s and Related Dementias Neuroimaging (SCAN) are available through https://naccdata.org/requesting-data/data-request-process. Data from the Health and Aging Brain Study–Health Disparities (HABS-HD) are available upon request from https://apps.unthsc.edu/itr/reports.

Data from the Anti-Amyloid Treatment in Asymptomatic Alzheimer’s Disease (A4) Study and the Longitudinal Evaluation of Amyloid Risk and Neurodegeneration (LEARN) Study are available upon registration and acceptance of the data-use agreement at https://www.a4studydata.org/.

Data from the Korean Brain Aging Study for the Early Diagnosis and Prediction of Alzheimer’s Disease (KBASE) are available through the LONI data repository, subject to study-specific access requirements. Data from the Religious Orders Study (ROS), Rush Memory and Aging Project (MAP) and Minority Aging Research Study (MARS) are available upon request from the Rush Alzheimer’s Disease Center at https://www.radc.rush.edu/.

Data from the Wisconsin Registry for Alzheimer’s Prevention (WRAP) are available upon request from https://wrap.wisc.edu/data-requests-2/.

Data from the Washington Heights–Inwood Columbia Aging Project (WHICAP) and the Knight Alzheimer Disease Research Center (Knight ADRC) are available upon reasonable request to the respective study investigators, subject to institutional approval and applicable data-use agreements.

## Code availability

The normative WMH reference package, including twelve age- and sex-stratified cognitively unimpaired WMH frequency maps in MNI152 2 mm space, fitted burden and vTRM percentile models, the spline-based conditioning model, normative reference distributions, regional atlas files and the single-scan scoring tool, will be made available through the Open Science Framework (OSF; https://osf.io/swzau/). The Brain Lesion Analytics platform (https://brainlesionanalytics.org/) provides interactive normative WMH scoring and automated WMH segmentation using the deployed 3D U-Net model developed and trained as part of this study data.

The package contains the model coefficients, regional parameters and constants required to reproduce individualized scoring and assign lobar and lesion-depth descriptors. The scoring tool accepts a binary WMH lesion mask in MNI152 2 mm space, together with age and sex, and generates WMH burden and vTRM percentiles, quadrant classification and a visual report. The web application also supports automated WMH segmentation using the deployed 3D U-Net. Supporting reference files and release documentation will be accessible through the same website. Software versions are provided in the Methods.

## Acknowledgements

The ADSP Phenotype Harmonization Consortium (ADSP-PHC) is funded by NIA (U24 AG074855, U01 AG068057 and R01 AG059716). The harmonized cohorts within the ADSP-PHC include: the Anti-Amyloid Treatment in Asymptomatic Alzheimer’s study (A4 Study), a secondary prevention trial in preclinical Alzheimer’s disease, aiming to slow cognitive decline associated with brain amyloid accumulation in clinically normal older individuals. The A4 Study is funded by a public-private-philanthropic partnership, including funding from the National Institutes of Health-National Institute on Aging, Eli Lilly and Company, Alzheimer’s Association, Accelerating Medicines Partnership, GHR Foundation, an anonymous foundation and additional private donors, with in-kind support from Avid and Cogstate. The companion observational Longitudinal Evaluation of Amyloid Risk and Neurodegeneration (LEARN) Study is funded by the Alzheimer’s Association and GHR Foundation. The A4 and A4-LEARN Studies are led by Dr. Reisa Sperling at Brigham and Women’s Hospital, Harvard Medical School and Dr. Paul Aisen at the Alzheimer’s Therapeutic Research Institute (ATRI), University of Southern California. The A4 and LEARN Studies are coordinated by ATRI at the University of Southern California, and the data are made available through the Laboratory for Neuro Imaging at the University of Southern California. The participants screening for the A4 Study provided permission to share their de-identified data in order to advance the quest to find a successful treatment for Alzheimer’s disease. We would like to acknowledge the dedication of all the participants, the site personnel, and all of the partnership team members who continue to make the A4 and LEARN Studies possible. The complete A4 Study Team list is available on: a4study.org/a4-study-team.; Alzheimer’s Disease Neuroimaging Initiative (ADNI): Data collection and sharing for this project was funded by the Alzheimer’s Disease Neuroimaging Initiative (ADNI) (National Institutes of Health Grant U01 AG024904) and DOD ADNI (Department of Defense award number W81XWH-12-2-0012). ADNI is funded by the National Institute on Aging, the National Institute of Biomedical Imaging and Bioengineering, and through generous contributions from the following: AbbVie, Alzheimer’s Association; Alzheimer’s Drug Discovery Foundation; Araclon Biotech; BioClinica, Inc.; Biogen; Bristol-Myers Squibb Company; CereSpir, Inc.; Cogstate; Eisai Inc.; Elan Pharmaceuticals, Inc.; Eli Lilly and Company; EuroImmun; F. Hoffmann-La Roche Ltd and its affiliated company Genentech, Inc.; Fujirebio; GE Healthcare; IXICO Ltd.;Janssen Alzheimer Immunotherapy Research & Development, LLC.; Johnson & Johnson Pharmaceutical Research & Development LLC.; Lumosity; Lundbeck; Merck & Co., Inc.;Meso Scale Diagnostics, LLC.; NeuroRx Research; Neurotrack Technologies; Novartis Pharmaceuticals Corporation; Pfizer Inc.; Piramal Imaging; Servier; Takeda Pharmaceutical Company; and Transition Therapeutics. The Canadian Institutes of Health Research is providing funds to support ADNI clinical sites in Canada. Private sector contributions are facilitated by the Foundation for the National Institutes of Health (www.fnih.org). The grantee organization is the Northern California Institute for Research and Education, and the study is coordinated by the Alzheimer’s Therapeutic Research Institute at the University of Southern California. ADNI data are disseminated by the Laboratory for Neuro Imaging at the University of Southern California; Brain tissue and additional biological samples were collected from individuals who were assessed and followed in the Case Western ADC which was funded by NIA P50AG008012; Estudio Familiar de Influencia Genetica en Alzheimer (EFIGA): 5R37AG015473, RF1AG015473, R56AG051876; the Health & Aging Brain Study – Health Disparities (HABS-HD), supported by the National Institute on Aging of the National Institutes of Health under Award Numbers R01AG054073, R01AG058533, R01AG070862, P41EB015922, and U19AG078109; brain tissues were obtained from the Brain Bank at the John P. Hussman Institute for Human Genomics at the University of Miami Miller School of Medicine, with funding provided by Louis D. Scientific Prize from the Institut de France; the Indiana Alzheimer’s Disease Research Center (IADRC) supported by National Institutes of Health (NIH) grant P30AG072976; the Korean Brain Aging Study for the Early Diagnosis and Prediction of Alzheimer’s disease (KBASE), which was supported by a grant from Ministry of Science, ICT and Future Planning (Grant No: NRF-2014M3C7A1046042, U01AG072177); Memory & Aging Project at Knight Alzheimer’s Disease Research Center (MAP at Knight ADRC): The Memory and Aging Project at the Knight-ADRC (Knight-ADRC). This work was supported by the National Institutes of Health (NIH) grants R01AG064614, R01AG044546, RF1AG053303, RF1AG058501, U01AG058922 and R01AG064877 to Carlos Cruchaga. The recruitment and clinical characterization of research participants at Washington University was supported by NIH grants P30AG066444, P01AG03991, and P01AG026276. Data collection and sharing for this project was supported by NIH grants RF1AG054080, P30AG066462, R01AG064614 and U01AG052410. We thank the contributors who collected samples used in this study, as well as patients and their families, whose help and participation made this work possible. This work was supported by access to equipment made possible by the Hope Center for Neurological Disorders, the Neurogenomics and Informatics Center (NGI: https://neurogenomics.wustl.edu/) and the Departments of Neurology and Psychiatry at Washington University School of Medicine; the University of Miami Brain Endowment Bank (Miami Brain Bank); National Alzheimer’s Coordinating Center (NACC): The NACC database is funded by NIA/NIH Grant U24 AG072122. SCAN is a multi-institutional project that was funded as a U24 grant (AG067418) by the National Institute on Aging in May 2020. Data collected by SCAN and shared by NACC are contributed by the NIA-funded ADRCs as follows: P30 AG062429 (PI James Brewer, MD, PhD), P30 AG066468 (PI Oscar Lopez, MD), P30 AG062421 (PI Bradley Hyman, MD, PhD), P30 AG066509 (PI Thomas Grabowski, MD), P30 AG066514 (PI Mary Sano, PhD), P30 AG066530 (PI Helena Chui, MD), P30 AG066507 (PI Marilyn Albert, PhD), P30 AG066444 (PI John Morris, MD), P30 AG066518 (PI Jeffrey Kaye, MD), P30 AG066512 (PI Thomas Wisniewski, MD), P30 AG066462 (PI Scott Small, MD), P30 AG072979 (PI David Wolk, MD), P30 AG072972 (PI Charles DeCarli, MD), P30 AG072976 (PI Andrew Saykin, PsyD), P30 AG072975 (PI David Bennett, MD), P30 AG072978 (PI Neil Kowall, MD), P30 AG072977 (PI Robert Vassar, PhD), P30 AG066519 (PI Frank LaFerla, PhD), P30 AG062677 (PI Ronald Petersen, MD, PhD), P30 AG079280 (PI Eric Reiman, MD), P30 AG062422 (PI Gil Rabinovici, MD), P30 AG066511 (PI Allan Levey, MD, PhD), P30 AG072946 (PI Linda Van Eldik, PhD), P30 AG062715 (PI Sanjay Asthana, MD, FRCP), P30 AG072973 (PI Russell Swerdlow, MD), P30 AG066506 (PI Todd Golde, MD, PhD), P30 AG066508 (PI Stephen Strittmatter, MD, PhD), P30 AG066515 (PI Victor Henderson, MD, MS), P30 AG072947 (PI Suzanne Craft, PhD), P30 AG072931 (PI Henry Paulson, MD, PhD), P30 AG066546 (PI Sudha Seshadri, MD), P20 AG068024 (PI Erik Roberson, MD, PhD), P20 AG068053 (PI Justin Miller, PhD), P20 AG068077 (PI Gary Rosenberg, MD), P20 AG068082 (PI Angela Jefferson, PhD), P30 AG072958 (PI Heather Whitson, MD), P30 AG072959 (PI James Leverenz, MD); National Institute on Aging Alzheimer’s Disease Family Based Study (NIA-AD FBS): U24 AG056270; Religious Orders Study (ROS)/ Memory and Aging Project (MAP - Rush)/ Minority Aging Research Study (MARS) is supported by P30AG10161, P30AG72975, R01AG17917, R01AG015819, R01AG22018, R01AG056405, R01AG064233, and UH3NS100599. Washington Heights/Inwood Columbia Aging Project (WHICAP): RF1 AG054023, R01 AG034189, UL1TR001873 and UM1 TR006153; this manuscript has been reviewed by WHICAP investigators for scientific content and consistency of data interpretation with previous WHICAP Study publications. We acknowledge the WHICAP study participants and the WHICAP research and support staff for their contributions to this study. Wisconsin Registry for Alzheimer’s Prevention (WRAP): R01AG027161 and R01AG054047. Additional acknowledgments include the National Institute on Aging Genetics of Alzheimer’s Disease Data Storage Site (NIAGADS, U24AG041689) at the University of Pennsylvania, funded by NIA.

## Notes

https://adseq.org/

https://www.radc.rush.edu/

https://www.a4studydata.org/

https://apps.unthsc.edu/itr/reports

https://adni.loni.usc.edu/

https://wrap.wisc.edu/data-requests-2/

### Competing Interest Statement

The authors have declared no competing interest.

