## supplementary information for "Topographic reference modelling of white matter hyperintensities across the adult lifespan"

### S1. Sample and data

#### F1. Sample construction and analytic subsets.

The analytic sample comprises 42,469 FLAIR-derived WMH lesion masks from 22,084 participants across twelve contributing studies, all represented in MNI152 2 mm standard space. Different analyses draw on different subsets of this sample, and the figure below sets out how each is derived. The normative reference is fitted on the 27,247 scans from participants who were cognitively unimpaired at every visit; the one-scan-per-participant refit uses the earliest scan from each of the 13,303 participants contributing to that reference. Two-axis analyses require both a burden percentile and a vTRM percentile, available for 41,409 scans, of which 21,465 are baseline scans used in the participant-level sensitivity analysis. Regional compartment analyses draw on the 41,438 scans with regional lesion counts.

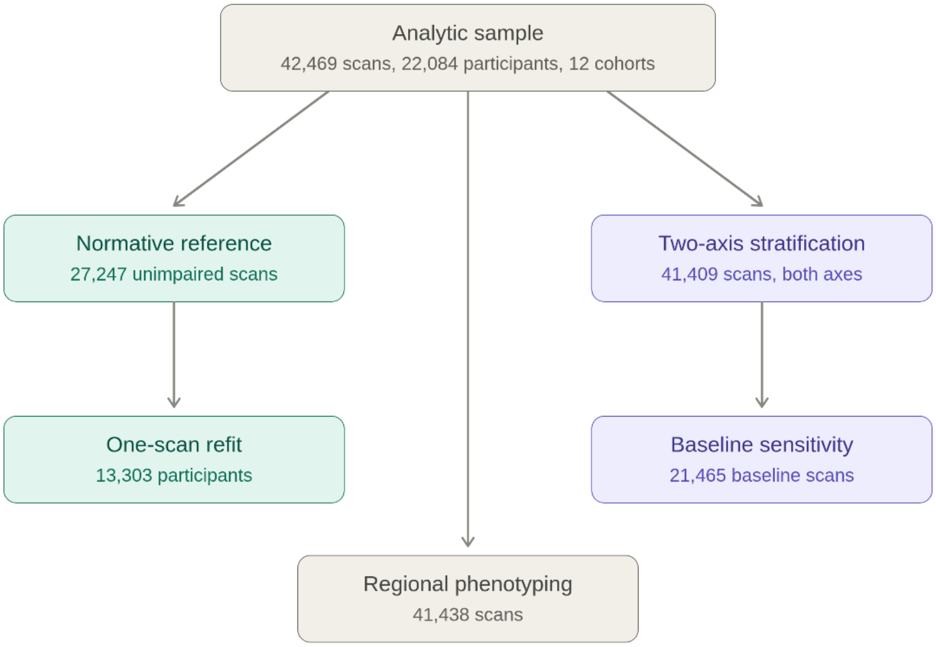

**Figure S1.F1 | Analytic sample and its subsets. Counts of scans and participants contributing to each analysis.** Teal indicates subsets of the cognitively unimpaired reference sample; purple indicates subsets requiring both burden and vTRM percentiles.

#### F2. Cohort composition, demographics, and diagnosis distribution.

Each contributing study differs in imaging density, follow-up, and participant characteristics. Supplementary Table S1.T2 gives scans, participants, scans per participant, median follow-up among participants with repeat imaging, mean years of education, APO-ε4 carrier percentage among those genotyped, and the proportion of scans acquired at 3T, for each study. Diagnosis and age distributions by study are given in Extended Data Table 1. Imaging density varies substantially: A4 contributes 4.83 scans per participant, whereas WHICAP contributes exactly one, and NACC and HABS-HD fewer than two. APO-ε4 carrier rates range from 17.5% in ROS to 60.3% in A4, reflecting the enrichment criteria of the contributing studies. Genotype was unavailable for KBASE. Most studies imaged exclusively at 3 T; WHICAP, MAP, ROS and NACC also include scans acquired at 1.5 T.

| Cohort | Scans | Participants | Scans per participant | Follow-up (median yr) | Education (mean) | APOE4 carrier % | Field 3T % |
|---|---|---|---|---|---|---|---|
| A4 | 6036 | 1250 | 4.83 | 1.7 | 16.6 | 60.3 | 100.0 |
| ADNI | 7234 | 2364 | 3.06 | 1.1 | 16.3 | 42.1 | 99.9 |
| HABS-HD | 5859 | 3673 | 1.6 | 2.0 | 13.4 | 27.5 | 100.0 |
| KBASE | 1170 | 602 | 1.94 | 2.0 | 11.4 | — | 100.0 |
| Knight ADRC | 1185 | 781 | 1.52 | 2.9 | 16.2 | 36.3 | 100.0 |
| LEARN | 917 | 535 | 1.71 | 1.4 | 16.8 | 22.2 | 100.0 |
| MAP | 2576 | 1032 | 2.5 | 2.1 | 15.9 | 23.1 | 76.4 |
| MARS | 649 | 287 | 2.26 | 2.5 | 15.5 | 33.1 | 99.5 |
| NACC | 13103 | 9048 | 1.45 | 1.6 | 15.8 | 38.3 | 82.8 |
| ROS | 193 | 95 | 2.03 | 2.1 | 19.5 | 17.5 | 77.2 |
| WHICAP | 1682 | 1682 | 1.0 | — | 11.7 | 27.7 | 60.8 |
| WRAP | 1865 | 735 | 2.54 | 3.0 | 16.6 | 37.4 | 100.0 |

#### F3. Between-cohort harmonisation: ComBat evaluation against cohort-adjusted modelling

Total WMH volume was harmonised with ComBat-GAM (neuroHarmonize v2.4.5), preserving a smooth age term and sex. Because ComBat assumes approximately Gaussian residuals and WMH volume is right-skewed with values near zero (S2.F1), harmonisation was performed on the log1p scale and back-transformed. As a single phenotype was harmonised, ComBat was run in its univariate form without empirical-Bayes shrinkage, which pools across features and is undefined for one; batch location and scale were estimated directly. Models were fitted on baseline scans and applied to all scans. Six batch definitions were compared: study; manufacturer; manufacturer × field strength; manufacturer × field × slice class (2D/3D); study × manufacturer × field; and study × manufacturer × field × scanner model. Batches with fewer than five baseline scans were folded into the next coarser level. Residual cohort effect was quantified as $\eta^2$ of cohort on age- and sex-adjusted log WMH. Associations with demographic, cognitive, vascular and biomarker variables were estimated before and after harmonisation using mixed models with a random intercept for cohort, adjusting for an age spline and sex.

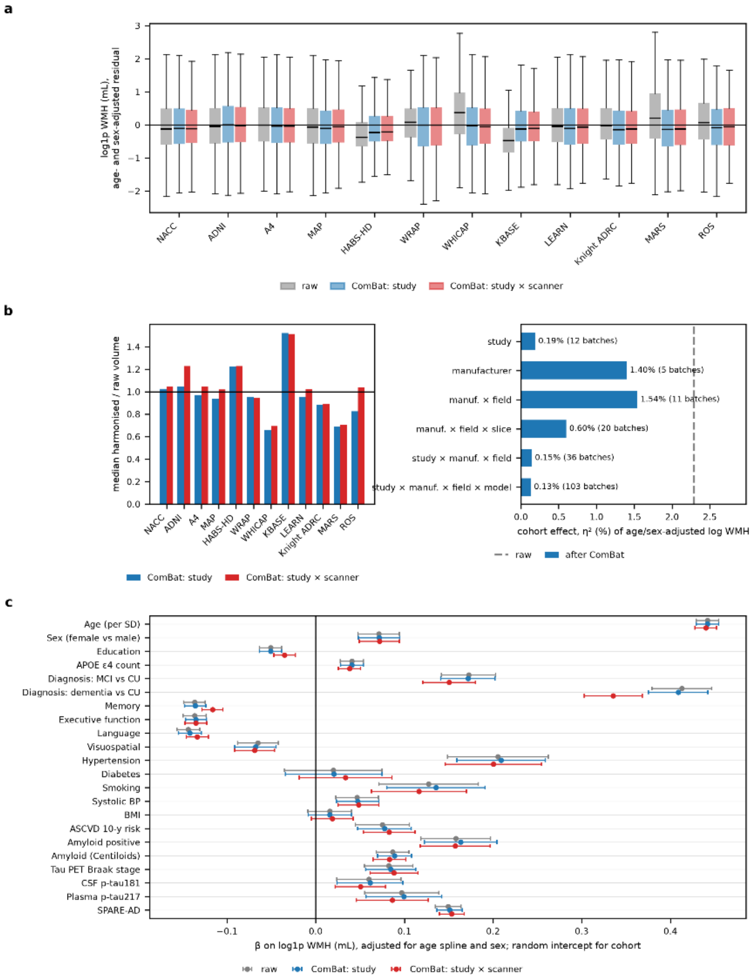

**S1.F3. | Between-cohort harmonisation: ComBat evaluation against cohort-adjusted modelling.** a, Age- and sex-adjusted log WMH by cohort for raw volumes (grey) and after ComBat with study (blue) or study × scanner (red) batches. Raw medians sit above zero in cohorts with high WMH for age (WHICAP, MARS, ROS, Knight ADRC) and below zero in cohorts with low WMH (KBASE, HABS-HD); both ComBat variants centre every cohort on zero while leaving within-cohort spread unchanged. b, Left: median per-cohort correction factor (harmonised/raw). Although cohort explained only 2.3% of adjusted variance, corrections ranged from ×0.66 (WHICAP) and ×0.69 (MARS) to ×1.23 (HABS-HD) and ×1.52 (KBASE); study and study × scanner batches diverge only where scanner composition differs within a cohort (ADNI, ROS). Right: residual cohort effect after harmonisation under each batch definition (dashed line, raw). Definitions containing study removed 92–94% of the cohort effect; scanner-only definitions removed 33–74%, indicating that cohort differences are not explained by scanner metadata alone. c, Standardised associations (β, 95% CI) with WMH before and after harmonisation. No association changed direction or significance. Study-level ComBat left all effect sizes within 7% of raw. Study × scanner ComBat attenuated effects of dementia diagnosis (−19%), MCI (−13%), memory (−15%), education (−31%) and plasma p-tau217 (−11%), while age, sex, hypertension, systolic blood pressure and amyloid were unchanged (<4%), consistent with scanner batches partly absorbing real clinical differences when diagnostic groups are unevenly distributed across scanners within a cohort. The cohorts most strongly corrected are those with known population differences in vascular burden, which ComBat cannot separate from acquisition effects. These results support retaining the cohort-adjusted quantile-regression reference as the primary model (S2.F5), with ComBat reported as a sensitivity analysis. Numerical values in Supplementary Tables S1.T3a–c.

#### S2. Normative modelling of WMH burden

##### F1. Distributional properties of WMH volume: zero fraction, skewness, kurtosis, dip test, mixture BIC.

The choice of modelling framework depends on the shape of the outcome distribution, so that shape was characterised before any reference curves were fitted. Log-transformed WMH volume was summarised in the cognitively unimpaired sample (n = 27,247) by zero fraction, skewness, kurtosis, Hartigan's dip test for unimodality and the Bayesian information criterion for one- and two-component Gaussian mixtures, separately for each compartment, sex and age band. Volumes are right skewed at younger ages and approach symmetry in the oldest band, and compartments with small absolute volumes carry a substantial proportion of exact zeros, reaching 17% for parietal WMH below 60 years. Both features argue against a Gaussian family model and motivate the distribution-free approach used throughout.

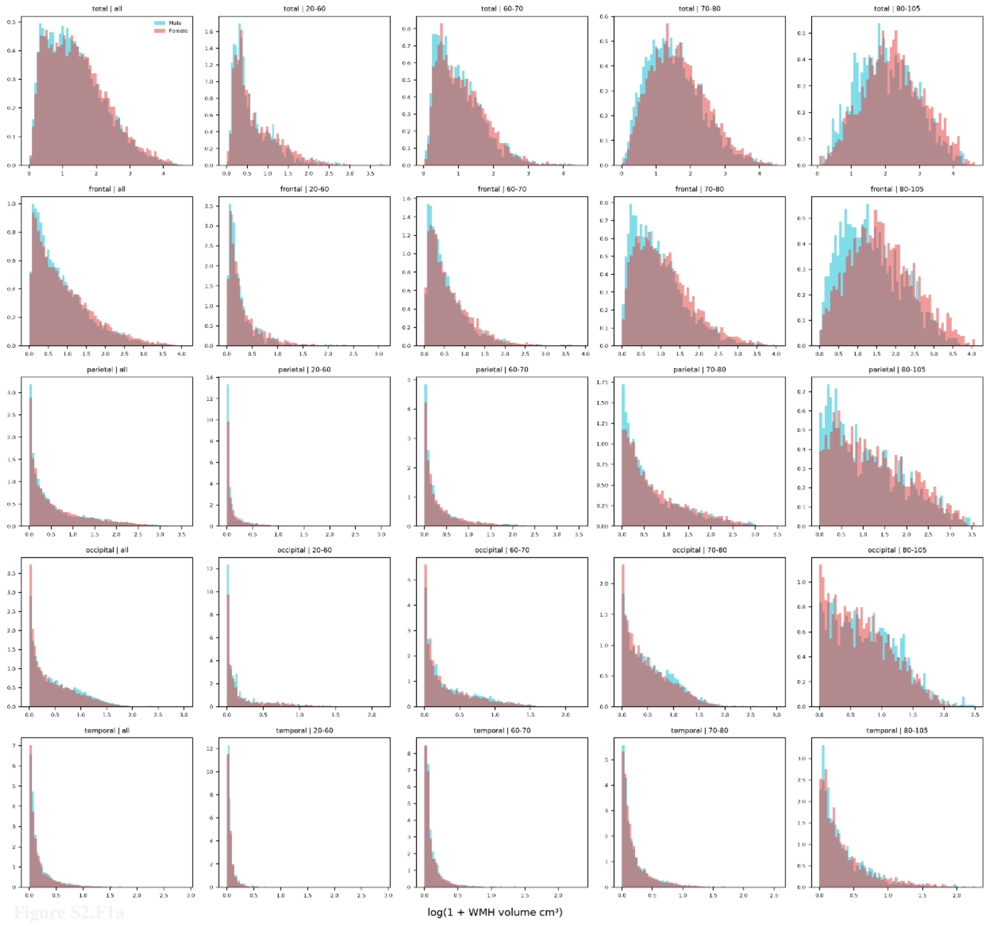

**Figure S2.F1 | Distribution of log-transformed WMH volume. Histograms of log(1 + WMH volume) in cognitively unimpaired participants (n = 27,247).** a, Total WMH and the four lobar compartments. b, The anterior, middle and posterior cerebral artery territories and the periventricular and ventricle-distal compartments. In both panels, columns show the pooled sample and four age bands, with male (red) and female (cyan) distributions overlaid. Distributions are right skewed at younger ages and approach symmetry in the oldest band. Corresponding zero fractions, skewness, kurtosis, Hartigan dip test statistics and Gaussian mixture BIC values are given in Supplementary Table S2.T1. Distributions for compartments are given in Supplementary Tables, S2.T1a.csv and S2.T1b.csv.

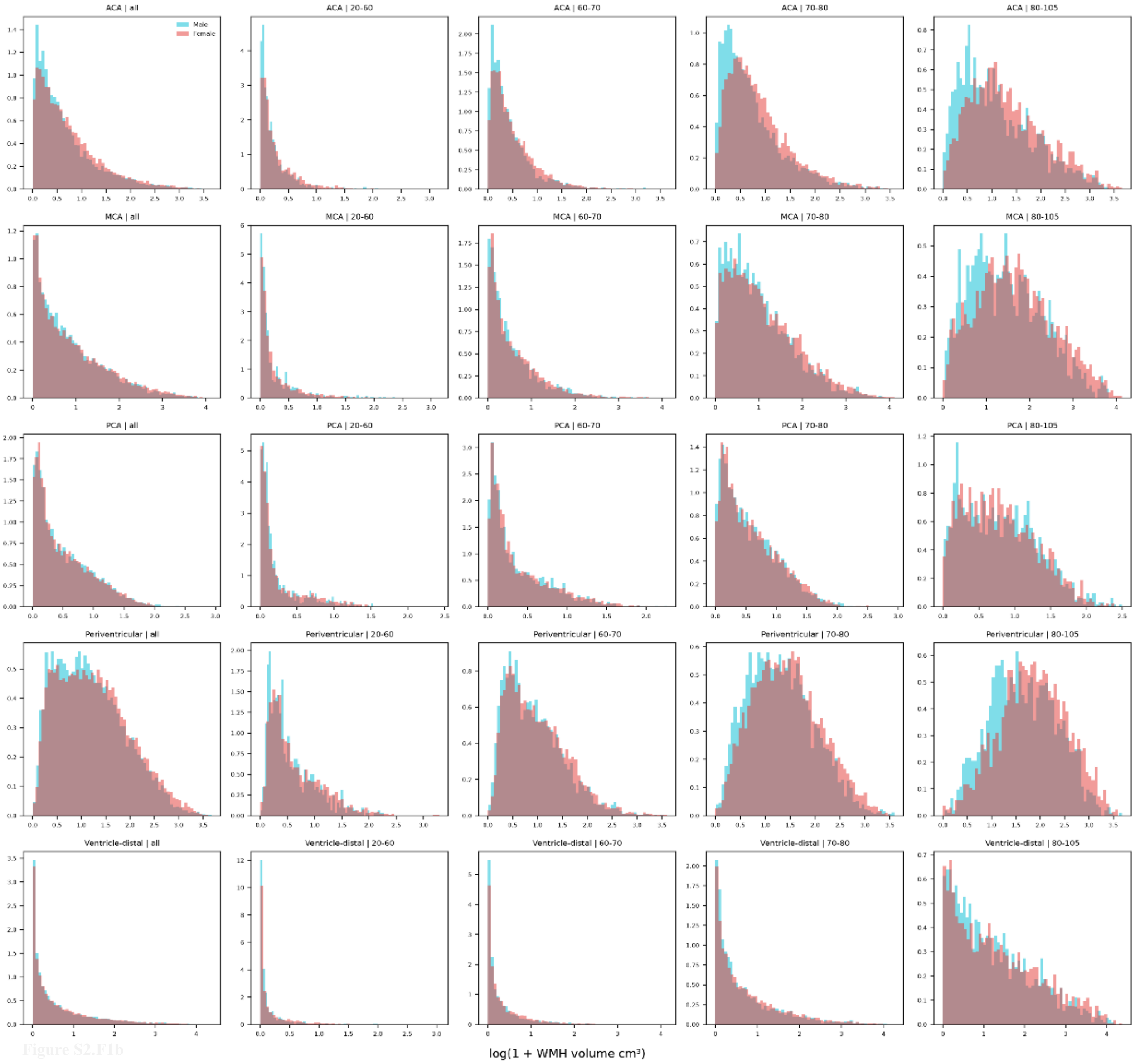

Figure S2.F1b

#### F2. Choice of quantile regression: comparison against Gaussian family GAM, total burden.

We compared quantile regression (QR) with a Gaussian-family generalized additive model (GAM) to determine which method provides more reliable percentile estimates of total WMH burden. QR estimates percentiles directly from the data, whereas GAM calculates them using assumptions about how the measurements are distributed. Both methods produced similar percentile rankings for most scans. However, QR better matched the observed data for scans with unusually high burden. GAM also showed unstable and potentially unrealistic changes in its fitted curves for individuals younger than 50 and older than 80 years. Overall, QR provided more reliable estimates in the regions where accurate percentile scoring is particularly important.

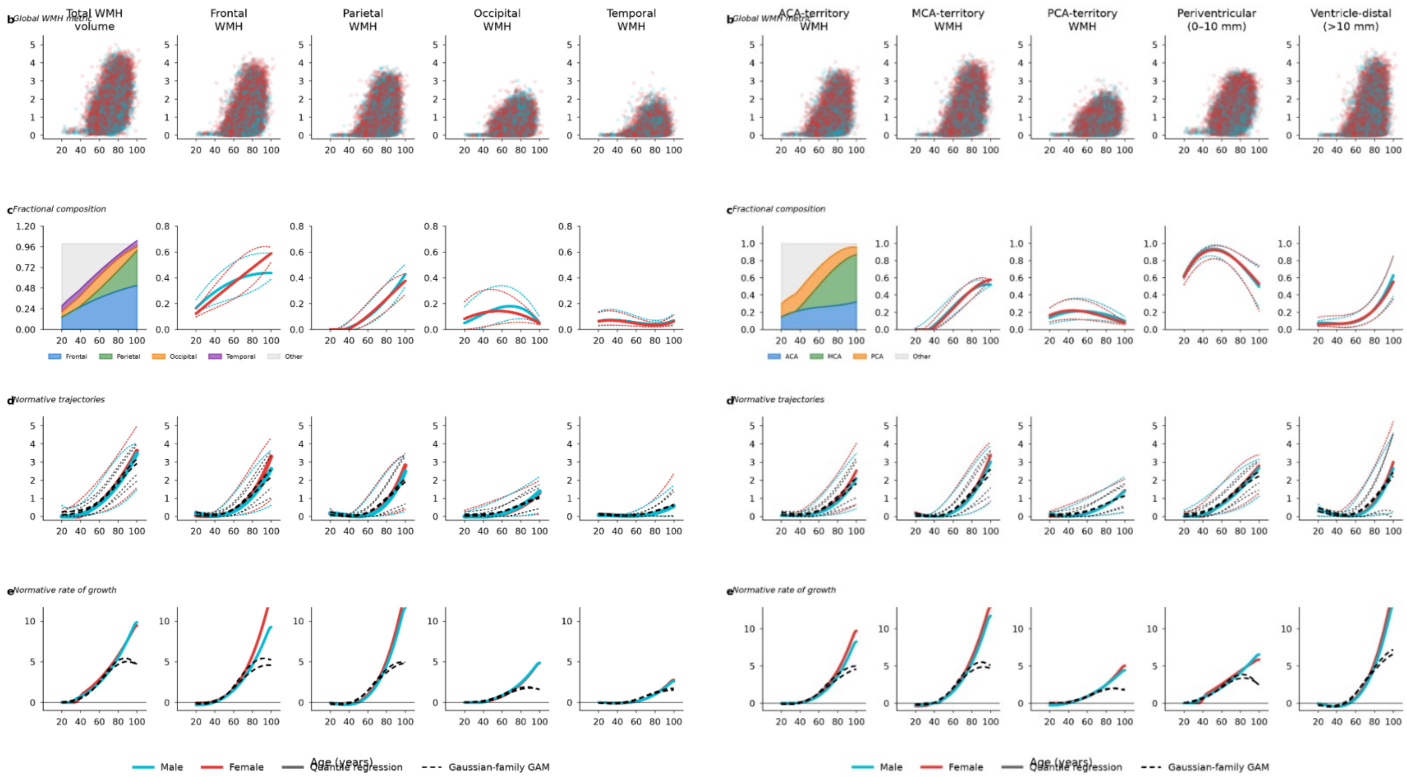

**Figure S2.F2 | Quantile regression compared with a Gaussian-family generalized additive model.** Normative modelling of WMH burden in cognitively unimpaired participants, using the same data and specification as main-text Figures 1 and 2. a, Total and lobar compartments. b, The anterior, middle and posterior cerebral artery territories and the periventricular (0–10 mm) and ventricle-distal (>10 mm) compartments. Within each panel: raw log(1 + WMH volume) against age, all participants, coloured by sex; fractional composition, showing stacked median fractions and fraction trajectories with 25$^{th}$, 50$^{th}$ and 75$^{th}$ percentiles, QR only; normative trajectories showing the 5$^{th}$, 50$^{th}$ and 95$^{th}$ percentiles, with solid coloured lines for cubic quantile regression and black dashed lines for the Gaussian-family GAM, in which mean and log-variance are each modelled as smooth functions of age and percentiles derived from the normal distribution; and rate of change of the median trajectory with respect to log age. QR curves remain stable where data are sparse at the extremes of the age range, whereas the GAM comparator shows unstable behaviour below 50 and above 80 years. Fitted curves for both methods are given in Supplementary Table S2.T2a and S2.T2b.

#### F3. Comparison against GAMLSS for regional compartments.

GAMLSS is widely used in published brain-chart references, so the reference was also fitted using this framework to assess whether the choice of quantile regression influenced the results. Models used the generalized gamma distribution with fractional polynomials in age, fitted in two variants: without cohort terms and with contributing study included in the location and scale parameters. Scan-level percentile assignments from both variants were compared with those from quantile regression, together with model selection results and fitted curves. The frameworks agree closely at the median but diverge at younger ages, where the generalized gamma distribution requires strictly positive values and exact zero volumes were replaced with half the voxel volume, and above age 75, where fractional polynomial terms are unconstrained and the sample size decreases.

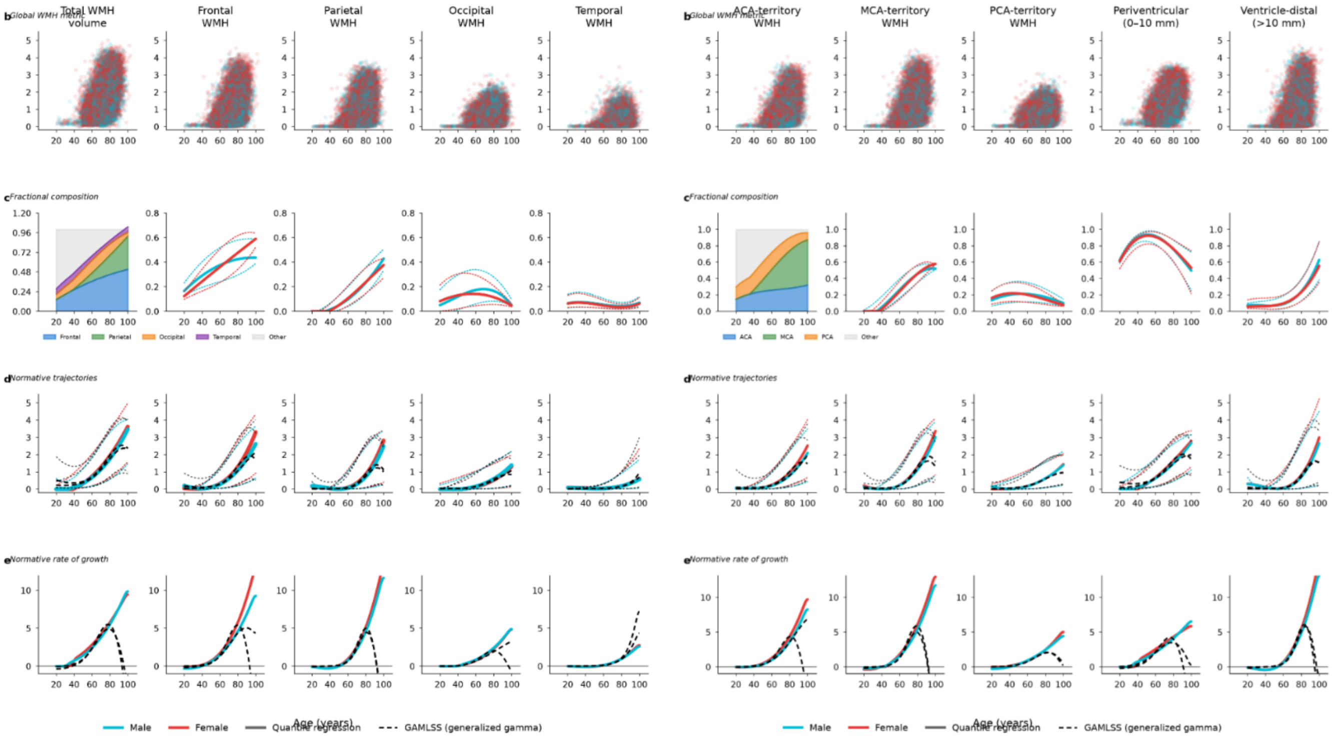

**Figure S2.F3 | Quantile regression compared with GAMLSS. Normative modelling of WMH burden in cognitively unimpaired participants, using the same data and specification as main-text Figures 1 and 2.** a, Total and lobar compartments. b, The anterior, middle and posterior cerebral artery territories and the periventricular (0–10 mm) and ventricle-distal (>10 mm) compartments. Within each panel: raw log(1 + WMH volume) against age, all participants, coloured by sex; fractional composition, showing stacked median fractions and fraction trajectories with 25$^{th}$, 50$^{th}$ and 75$^{th}$ percentiles, quantile regression only; normative trajectories showing the 5$^{th}$, 50$^{th}$ and 95$^{th}$ percentiles, with solid coloured lines for cubic quantile regression and black dashed lines for GAMLSS fitted with the generalized gamma distribution and fractional polynomials in age, without cohort terms; and rate of change of the median trajectory with respect to log age. GAMLSS trajectories are elevated at younger ages, where the requirement for strictly positive values necessitated replacing exact zero volumes with half the voxel volume, and decline above 75 years, where fractional polynomial terms are unconstrained and the sample thins. Fitted GAMLSS curves and model selection details are given in Supplementary Tables S2.T3a, S2.T3b, S2.T3c

#### F4. Empirical validation of fitted percentile curves against observed quantiles by decade.

A fitted percentile curve is only useful if it recovers the percentiles actually observed in the data. Within each decade of age, the 5th, 25th, 50th, 75th and 95th percentiles of log-transformed WMH volume were computed directly with no model, and each fitted curve evaluated at the mean age of the scans contributing to that decade. Deviation from the observed value was recorded for all three modelling frameworks. All three agree closely at the median. They differ in the tails, which is where percentile scoring is most consequential and where the sample is thinnest.

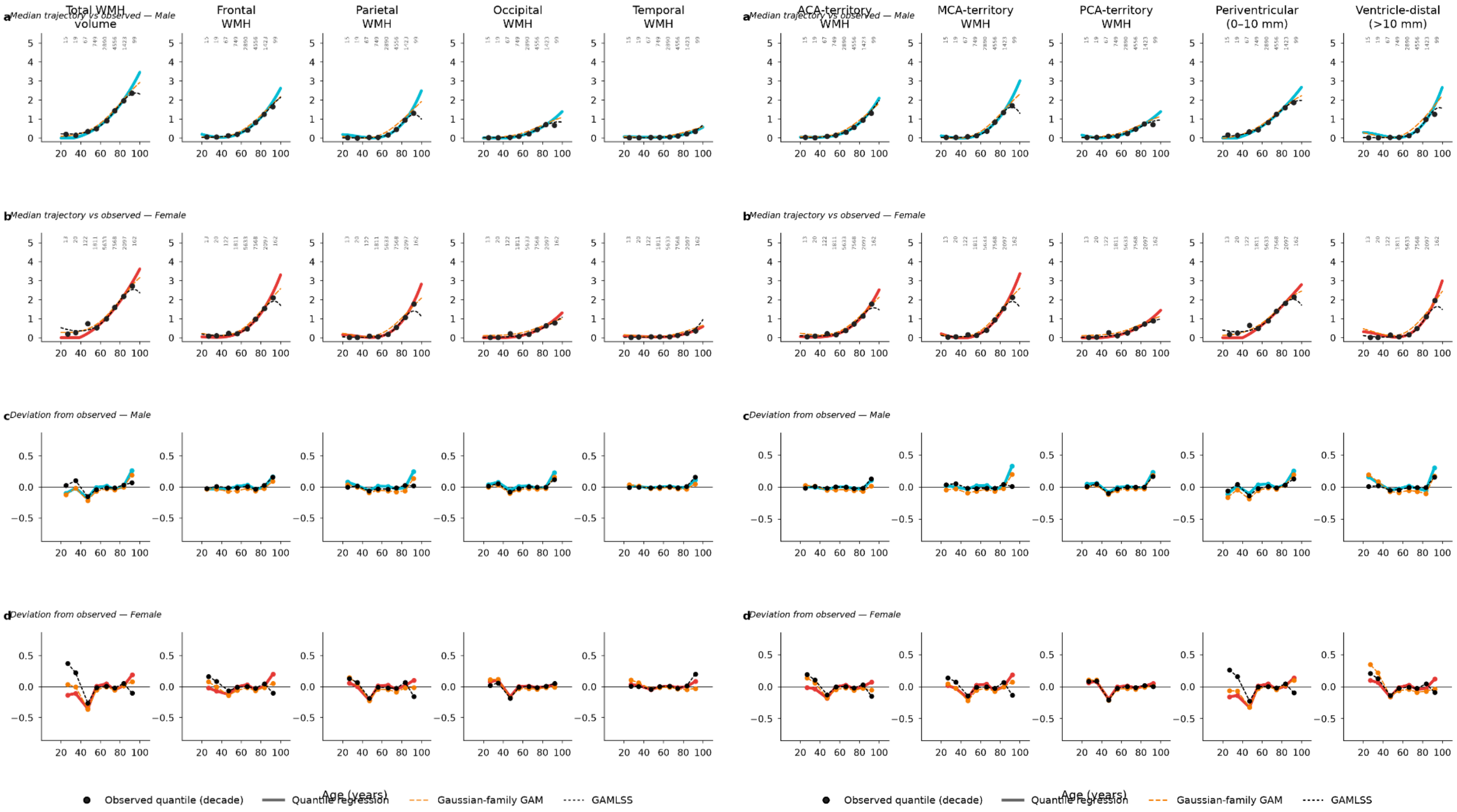

**Figure S2.F4 | Fitted percentile curves compared with observed quantiles by decade.** Empirical validation of the three modelling frameworks in cognitively unimpaired participants. Within each decade of age, the 5th, 25th, 50th, 75th and 95th percentiles of log(1 + WMH volume) were computed directly from the data with no model; fitted curves were then evaluated at the mean age of the scans contributing to that decade. a, Total and lobar compartments. b, The anterior, middle and posterior cerebral artery territories and the periventricular (0–10 mm) and ventricle-distal (>10 mm) compartments. Within each panel: median trajectories for males (cyan) and females (red), with solid coloured lines for cubic quantile regression, orange dashed lines for the Gaussian-family GAM and black dashed lines for GAMLSS, and points showing the observed decade-wise median with the number of contributing scans annotated above each band; and the deviation of each fitted curve from the corresponding observed quantile, averaged across all five percentiles, where values near zero indicate a curve that recovers the empirical distribution. All three methods agree closely at the median but differ in the tails, where percentile scoring is most consequential. At 80–100 years quantile regression recovers the 5th and 95th percentiles most closely, with mean absolute deviations of 0.03 and 0.11 log units for lobar compartments and 0.03 and 0.14 for arterial-territory and depth-distance compartments, whereas the Gaussian-family GAM departs substantially (0.32 and 0.44; 0.34 and 0.44). At 20–40 years GAMLSS shows the largest deviation at the 95$^{th}$ percentile (0.25 versus 0.14; 0.33 versus 0.13).

Mean absolute deviations by age range and percentile are given in Supplementary Table S2.T4, shown below for both compartment sets; the observed and fitted values underlying them are provided as separate files, S2.T4c.csv and S2.T4d.csv.

**a Lobar**

| band_group | n_comparisons | QR_mean_abs_dev | GAM_mean_abs_dev | GAMLSS_mean_abs_dev |
|---|---|---|---|---|
| all bands | 400 | 0.0666 | 0.1719 | 0.0639 |
| 20-40 (young) | 100 | 0.0735 | 0.0606 | 0.0775 |
| 40-80 (dense) | 200 | 0.0491 | 0.1859 | 0.0453 |
| 80-100 (old) | 100 | 0.0945 | 0.255 | 0.0877 |

**b Arterial-territory and depth-distance**

| band_group | n_comparisons | QR_mean_abs_dev | GAM_mean_abs_dev | GAMLSS_mean_abs_dev |
|---|---|---|---|---|
| all bands | 400 | 0.0739 | 0.1925 | 0.0705 |
| 20-40 (young) | 100 | 0.0792 | 0.0906 | 0.0982 |
| 40-80 (dense) | 200 | 0.0548 | 0.2029 | 0.0479 |
| 80-100 (old) | 100 | 0.1068 | 0.2736 | 0.088 |

#### F5. Cohort adjustment: QR ± cohort terms; GAMLSS with and without.

The published reference is fitted on the pooled sample without cohort indicators, so that scans from outside the contributing studies can be scored without requiring cohort membership. This analysis examines the consequences of that choice. Curves were refitted with contributing study included in both quantile regression and GAMLSS, and the cohort-specific and pooled trajectories compared. In quantile regression, cohort indicators enter as intercept shifts, so the cohort-specific curves share a common age dependence and differ in level. The vertical spread between cohort-specific curves illustrates the between-study variation not explicitly represented in the pooled reference.

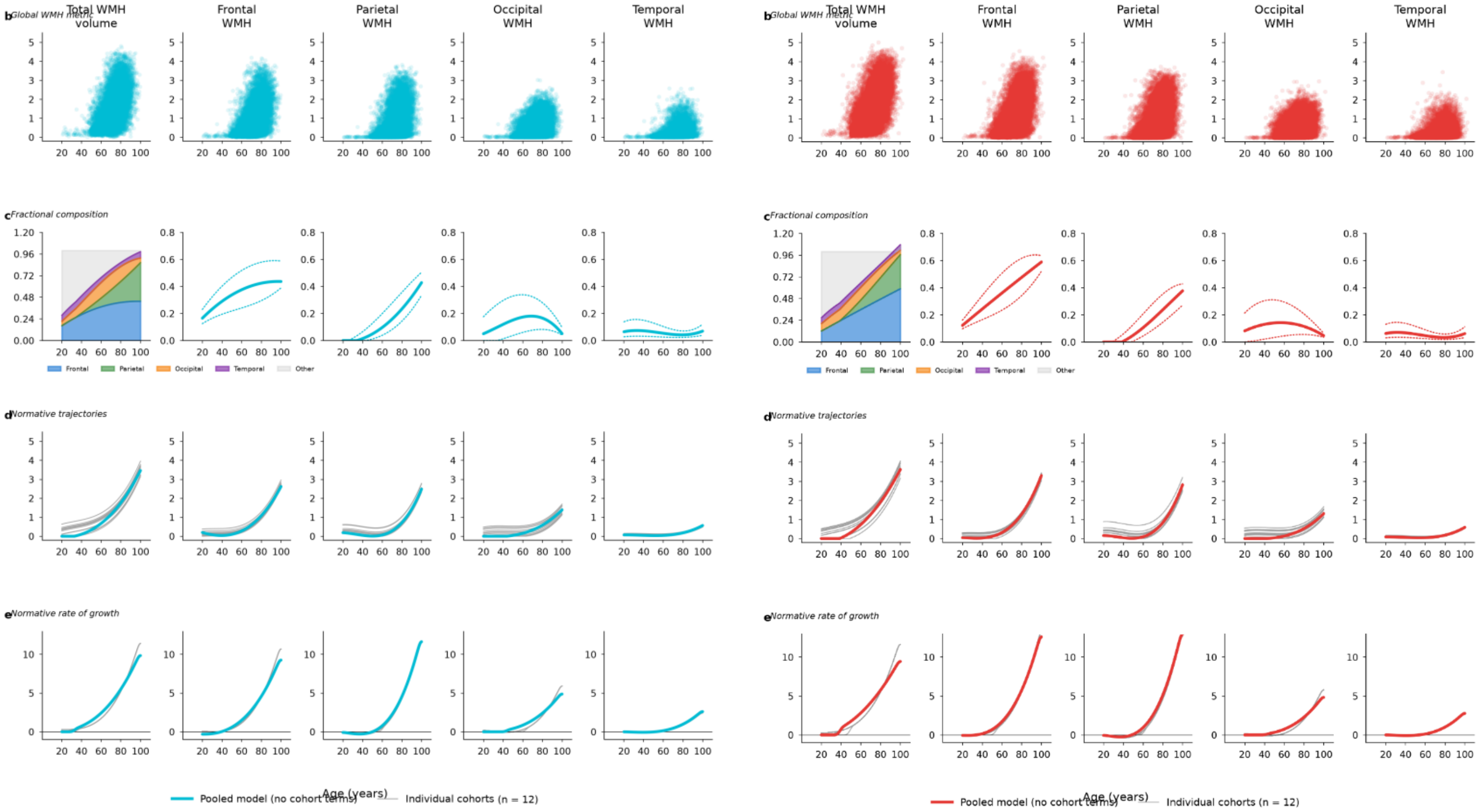

**Figure S2.F5 | Normative trajectories fitted with and without cohort terms across lobar compartments.** Models were fitted in cognitively unimpaired participants. Figures S2.F5a and S2.F5b show quantile regression in males (cyan) and females (red), respectively; Figures S2.F5c and S2.F5d (below) show GAMLSS in males and females, respectively.

Within each of the four figures, panels (b–e) show: (b) raw log(1 + WMH volume) versus age; (c) fractional composition, including stacked median lobar fractions and 25th, 50th, and 75th percentile trajectories from pooled quantile regression; (d) median normative trajectories; and (e) their rates of change with respect to log age. In Figures S2.F5a and S2.F5b, grey lines show cohort-specific medians from quantile regression with cohort indicators (one line per study), while coloured lines show the pooled model without cohort terms used in the main text. Cohort indicators act as intercept shifts, preserving age dependence while changing trajectory levels.

In Figures S2.F5c and S2.F5d, grey lines show cohort-specific medians from GAMLSS-B, with study included as a fixed factor in location and scale, while coloured lines show GAMLSS-A without cohort terms. Both use generalized gamma distributions with fractional polynomials in age. Zero volumes were replaced with half the voxel volume to ensure positivity. Vertical separation between cohort curves reflects between-study variation absorbed by cohort adjustment. Fitted curves are provided in Supplementary Table S2.T5.

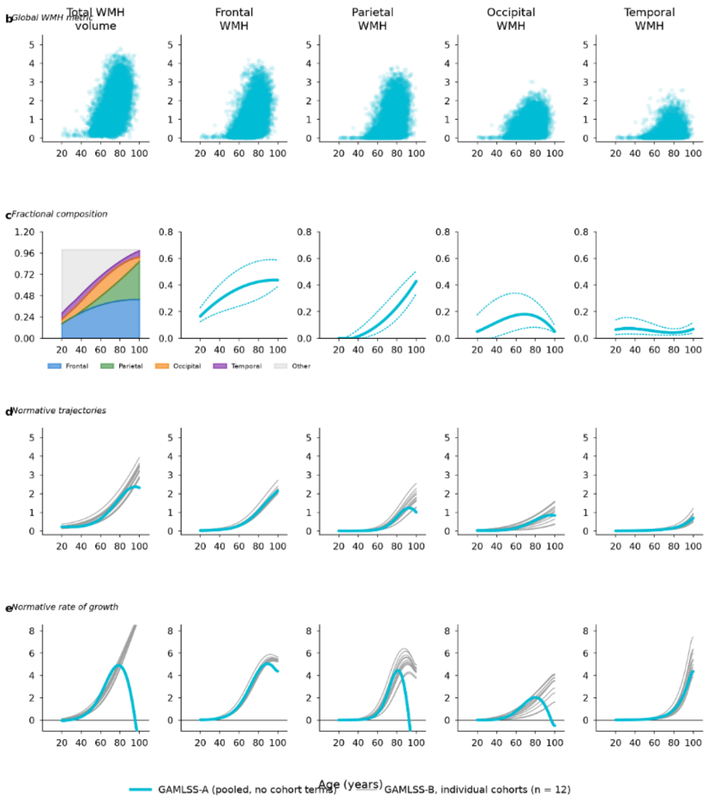

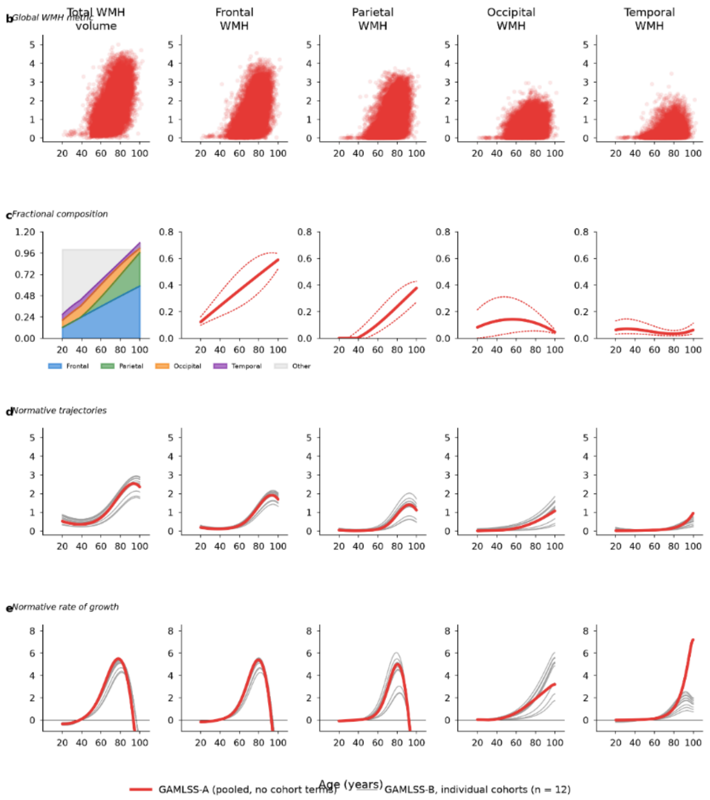

**Supplementary Table S2.T5 | Normative trajectories fitted with and without cohort terms**. a) Quantile regression percentile curves from models including cohort indicators (one curve per contributing study), alongside curves from the pooled model fitted without cohort terms, across the age range by compartment and sex. (b) GAMLSS model selection results, including polynomial degrees, Bayesian information criterion (BIC), and shape parameter for variant A (without cohort terms) and variant B (contributing study included as a fixed factor in the location and scale parameters), by compartment and sex. (c) Fitted GAMLSS curves from both model variants, across the age range by compartment and sex. Tables (a) and (c) are provided as separate CSV files (S2.T5a.csv and S2.T5c.csv), while Table (b) is presented below.

| compartment | sex | variant | n | npoly_mu | npoly_sigma | BIC | nu | minutes |
|---|---|---|---|---|---|---|---|---|
| frontal | Female | A | 17429 | 3 | 1 | 57196.8829 | 0.1602 | 42.7 |
| frontal | Female | B | 17429 | 3 | 1 | 56750.911 | 0.1552 | 65.6 |
| frontal | Male | A | 9818 | 2 | 1 | 29489.6461 | 0.153 | 0.4 |
| frontal | Male | B | 9818 | 2 | 1 | 29399.374 | 0.1675 | 0.7 |
| occipital | Female | A | 17429 | 1 | 2 | 18992.9088 | 0.6759 | 17.7 |
| occipital | Female | B | 17429 | 1 | 1 | 15525.8253 | 0.7222 | 12.1 |
| occipital | Male | A | 9818 | 2 | 2 | 13628.0721 | 0.7957 | 1.5 |
| occipital | Male | B | 9818 | 1 | 1 | 11551.9404 | 0.7523 | 0.7 |
| parietal | Female | A | 17429 | 3 | 1 | 34117.9507 | 0.1883 | 44.3 |
| parietal | Female | B | 17429 | 3 | 1 | 33025.9714 | 0.1809 | 35.7 |
| parietal | Male | A | 9818 | 3 | 2 | 17819.363 | 0.1708 | 2.1 |
| parietal | Male | B | 9818 | 2 | 2 | 17444.1457 | 0.1497 | 1.6 |
| temporal | Female | A | 17429 | 1 | 1 | -21708.3887 | 0.108 | 2.7 |
| temporal | Female | B | 17429 | 3 | 1 | -22536.5265 | 0.1151 | 35.4 |
| temporal | Male | A | 9818 | 1 | 2 | -11300.2536 | 0.1431 | 0.8 |
| temporal | Male | B | 9818 | 1 | 1 | -11619.764 | 0.1497 | 0.6 |
| total | Female | A | 17429 | 3 | 1 | 86107.4566 | 0.0479 | 41.2 |
| total | Female | B | 17429 | 3 | 1 | 84633.3956 | 0.0258 | 58.3 |
| total | Male | A | 9818 | 3 | 1 | 46793.5357 | 0.0316 | 0.9 |
| total | Male | B | 9818 | 1 | 1 | 46229.2442 | 0.0079 | 0.4 |

#### F6. Quantile crossing across the age range.

Quantile regression estimates each percentile independently, so nothing constrains the fitted curves to remain in order. Where they cross, percentile scoring is undefined. Curves were refitted using the same model specification as in the main analysis, and every adjacent percentile pair was tested at 300 points across the age range, recording the number of violations, the ages at which they occur, and their magnitude. Crossings were concentrated at the sparsely sampled age extremes, where several percentiles approached zero at younger ages. Elsewhere, the fitted curves remained ordered apart from a negligible local overlap.

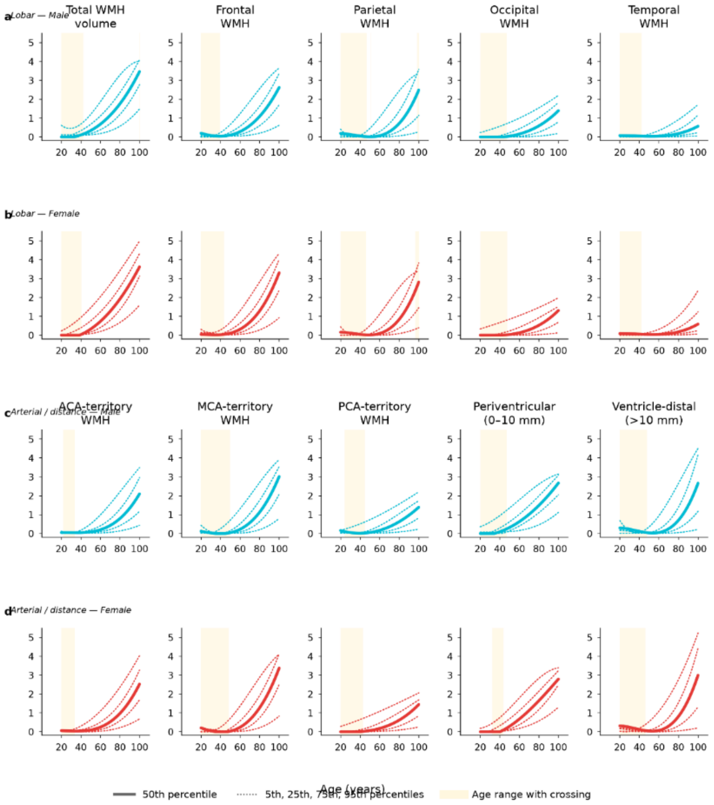

**Figure S2.F6 | Quantile crossing in the fitted normative curves. Quantile regression estimates each percentile independently, so the fitted curves are not constrained to remain ordered.** Curves were refitted exactly as in main-text Figures 1 and 2 (cubic in age, cognitively unimpaired participants, sex-stratified, no cohort terms) and every adjacent percentile pair was tested at 300 points across the age range. a, b, Lobar compartments in males and females. c, d, Arterial-territory and depth-distance compartments. Solid lines show the median; dotted lines the 5th, 25th, 75th and 95th percentiles. Shaded bands mark age ranges in which any adjacent pair is out of order. Crossings are concentrated below 40 years, where the cognitively unimpaired sample is sparse and several percentiles are close to zero. A few crossings also occur above 96 years, between the 75th and 95th percentiles in the parietal, total and MCA-territory compartments. Elsewhere, the curves remain ordered, apart from a negligible local overlap in parietal males at 50.5–51.0 years (0.0002 log units). The age ranges and magnitudes of these crossings are detailed in Supplementary Table S2.T6.

#### F7. Bootstrapped confidence bands on percentile curves.

The fitted curves are point estimates, and their precision varies with how densely each part of the age range is sampled. Cognitively unimpaired scans were resampled with replacement and the curves refitted 200 times, with bands spanning the 2.5th to 97.5th percentile of the refitted values at each age. Bands are narrow between approximately 50 and 90 years, where the sample is dense, and widen at both extremes where scans are sparse and the cubic term is least constrained. Mean band widths are reported by compartment, sex, percentile and age range.

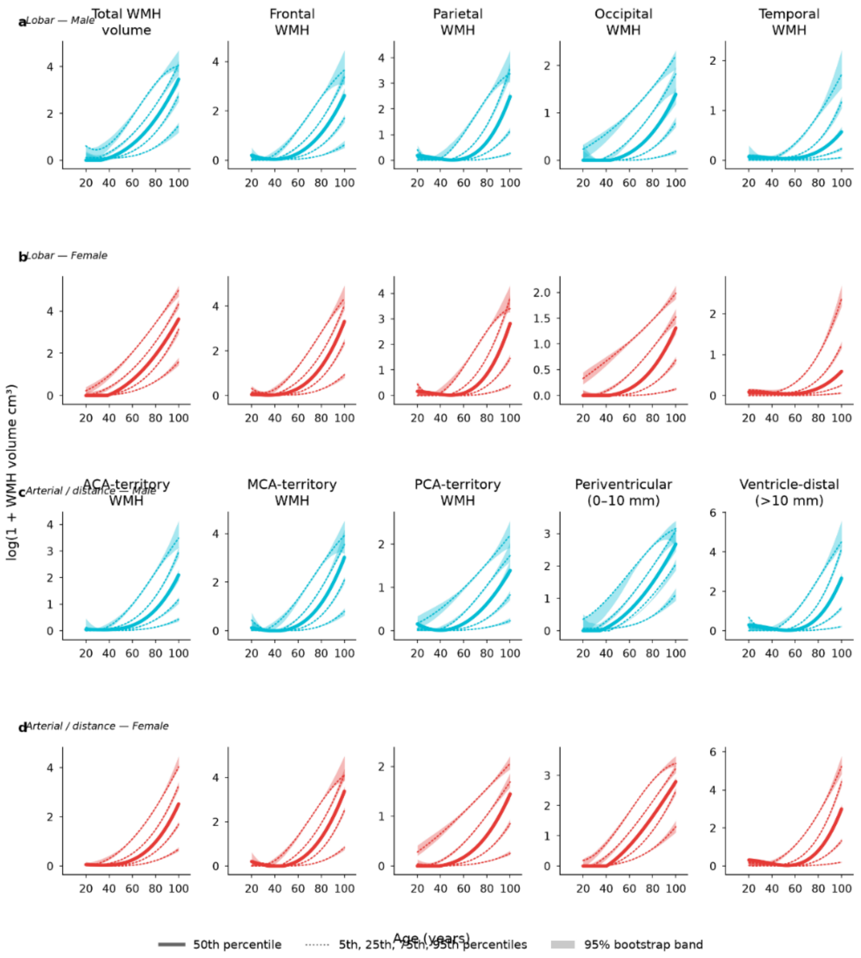

**Figure S2.F7 | Pointwise bootstrap 95% confidence bands for the normative percentile curves.** Cognitively unimpaired scans were resampled with replacement, and the quantile regression curves refitted 200 times, using the specification of main-text Figures 1 and 2 (cubic in age, sex-stratified, no cohort terms). Bands span the 2.5th to 97.5th percentile of the refitted values at each age. a, b, Lobar compartments in males and females. c, d, Arterial-territory and depth-distance compartments. Solid lines show the median; dotted lines the 5th, 25th, 75th and 95th percentiles. Vertical scales differ between panels so that band widths remain visible. Bands are narrow between approximately 50 and 90 years, where the sample is dense, and widen at both extremes where scans are sparse and the cubic term is least constrained. For the median curve, mean band widths range from 0.017–0.233 log units at 20–40 years, 0.007–0.063 at 40–80 years, and 0.054–0.185 at 80–100 years. Mean band widths by compartment, sex, percentile and age range are provided in Supplementary Table S2.T7a.csv, and the fitted values, band bounds, standard error and width across the age range in S2.T7b.csv.

##### F8. Out-of-sample validation: leave-one-cohort-out.

The reference is intended for scoring scans that did not contribute to it, so its performance was tested by holding out each contributing study in turn, refitting on the remaining eleven and scoring every held-out scan. Calibration was measured as the percentage of held-out scans falling below each nominal percentile curve. Three studies are well calibrated; the remainder show systematic deviations in a consistent direction across all five percentiles. These deviations reflect differences in percentile calibration but do not necessarily indicate differences in age-related trajectory shapes, which were evaluated separately in the cohort-adjusted analyses. Across studies, calibration deviations were associated with median WMH burden in cognitively unimpaired participants, while no statistically significant associations were observed with sample size, median age or sex ratio.

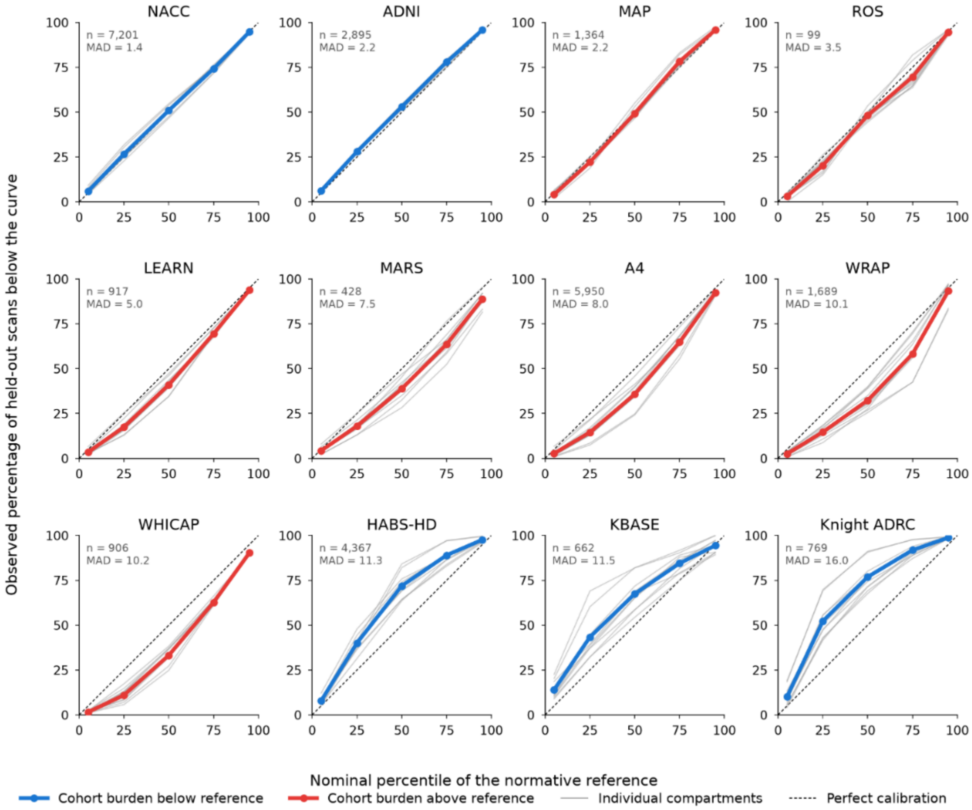

**Figure S2.F8 | Leave-one-cohort-out validation of the normative reference.** Each contributing study was held out in turn, quantile regression curves were refitted on the remaining eleven using the specification of main-text Figures 1 and 2, and every held-out scan was scored against the curve for its own sex. Panels show the percentage of held-out scans falling below each nominal percentile curve, ordered by mean absolute deviation; perfect calibration lies on the diagonal. Grey lines show individual compartments, colored lines their mean, with held-out n and mean absolute deviation (MAD, percentage points) annotated. NACC, ADNI and MAP are well calibrated (MAD 1.4–2.2 points). Remaining cohorts deviate in a single direction across all five percentiles; Knight ADRC, HABS-HD and KBASE carry lower burden than the pooled reference (mean deviation at the median +26.9, +21.9 and +17.4 points), while WRAP, WHICAP, A4 and MARS carry higher burden (−17.6, −16.8, −14.2 and −11.2). Deviations are largest at the median and smallest at the 5th and 95th percentiles, the signature of a shift in level rather than a difference in trajectory shape, consistent with the between-cohort spread shown in Supplementary Figures S5.F1–F4. Across cohorts, the calibration deviation is associated with median WMH burden of each study's cognitively unimpaired sample (Pearson $r = -0.78$, $P = 0.003$) and is unrelated to sample size ($r = 0.01$, $P = 0.99$), median age ($r = -0.33$, $P = 0.30$) or sex ratio ($r = -0.33$, $P = 0.30$); cohorts of similar median age differ in opposite directions, suggesting that differences in median age alone do not account for the observed calibration patterns. Cohort characteristics and mean absolute deviations are given in Supplementary Table S2.T8a, shown below; full coverage results are provided as a separate file, S2.T8b.csv.

| held_out_cohort | n_heldout | small | mean_abs_dev | max_abs_dev |
|---|---|---|---|---|
| NACC | 7201 | False | 1.42 | 6.76 |
| ADNI | 2895 | False | 2.18 | 4.44 |
| MAP | 1364 | False | 2.22 | 8.06 |
| ROS | 99 | False | 3.55 | 11.36 |
| LEARN | 917 | False | 5.03 | 15.54 |
| MARS | 428 | False | 7.54 | 22.66 |
| A4 | 5950 | False | 8.04 | 25.82 |
| WRAP | 1689 | False | 10.12 | 32.61 |
| WHICAP | 906 | False | 10.22 | 25.17 |
| HABS-HD | 4367 | False | 11.29 | 34.13 |
| KBASE | 662 | False | 11.54 | 44.03 |
| Knight ADRC | 769 | False | 16.05 | 44.83 |

#### F9. Longitudinal percentile tracking in participants with repeat scans.

A percentile is only interpretable as a property of an individual if it is stable within that person over time. Because absolute WMH volume rises substantially with age, a score that simply tracked volume would drift upward at every visit. Every scan from participants who were cognitively unimpaired at every visit and had two or more scans was scored against the reference, and first-to-last percentile change was measured alongside the rate of change per year of follow-up. Percentiles track the individual: first-to-last correlations are moderate to high, the median absolute change over the full follow-up is modest, and the distribution of annual change is centred on zero, indicating no systematic drift with age.

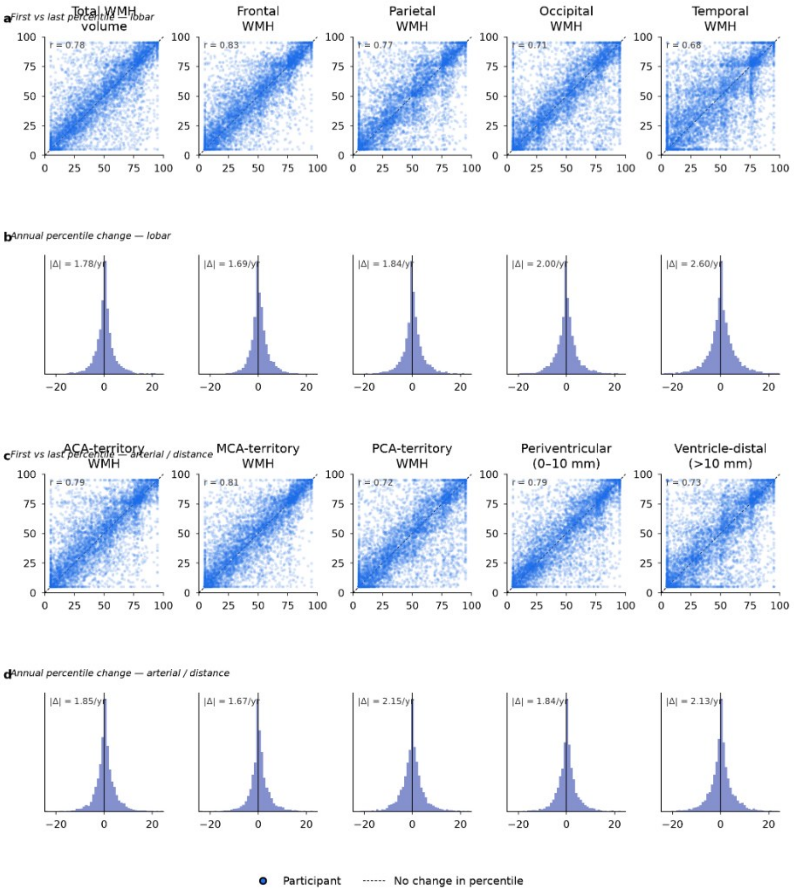

**Figure S2.F9 | Longitudinal stability of normative percentile scores.** Normative curves were fitted once on all 27,247 cognitively unimpaired scans using the specification of main-text Figures 1 and 2, and every scan from the 6,001 participants who were cognitively unimpaired at every visit and had two or more scans was scored against the curve for its own sex (19,945 scans; median 3 scans per participant, median follow-up 4.6 years). a, c, Each point is one participant, positioned by their percentile at the first scan (horizontal) and at the last (vertical), for lobar and for arterial-territory and depth-distance compartments. A participant whose standing relative to their peers is unchanged falls on the dashed diagonal; points above it moved up the distribution and points below moved down. Because absolute WMH volume rises substantially over follow-up, points clustering along the diagonal indicate that the percentile reflects the individual rather than their age. b, d, How far each participant's percentile moved per year of follow-up. A distribution concentrated at zero indicates stable scores, and its symmetry indicates no systematic drift up or down the scale with age. Percentiles track the individual: first-to-last correlations are 0.68–0.83 (highest for frontal, 0.83, and MCA territory, 0.81; lowest for temporal, 0.68), and the median absolute change over the full follow-up is 6.5–11.0 percentile points, equivalent to 1.7–2.6 points per year. Between 36% and 53% of participants move more than 10 percentile points and 11% to 21% more than 25 points over roughly 4.6 years. Summary statistics by compartment and sex are given in Supplementary Table S2.T9, shown below; per-participant results are provided as a separate file, S2.T9b.csv.

| compartment | n_participants | median_followup_yr | pearson_first_last | spearman_first_last | median_abs_change | median_within_sd | median_abs_annual_change | pct_moving_gt10 | pct_moving_gt25 |
|---|---|---|---|---|---|---|---|---|---|
| total | 6001 | 4.58 | 0.782 | 0.782 | 6.93 | 4.45 | 1.785 | 38.2 | 13.4 |
| frontal | 6001 | 4.58 | 0.828 | 0.828 | 6.71 | 4.14 | 1.686 | 36.2 | 11.1 |
| parietal | 6001 | 4.58 | 0.766 | 0.765 | 7.39 | 4.68 | 1.837 | 40.6 | 15.0 |
| occipital | 6001 | 4.58 | 0.71 | 0.709 | 7.56 | 5.08 | 2.001 | 41.3 | 15.9 |
| temporal | 6001 | 4.58 | 0.684 | 0.681 | 11.02 | 7.15 | 2.605 | 53.2 | 20.8 |
| ACA | 6001 | 4.58 | 0.787 | 0.787 | 7.49 | 4.73 | 1.855 | 40.2 | 13.5 |
| MCA | 6001 | 4.58 | 0.809 | 0.809 | 6.52 | 4.16 | 1.674 | 36.8 | 12.0 |
| PCA | 6001 | 4.58 | 0.722 | 0.723 | 8.21 | 5.29 | 2.146 | 43.3 | 15.9 |
| pv_total | 6001 | 4.58 | 0.786 | 0.786 | 7.08 | 4.57 | 1.844 | 39.2 | 13.6 |
| ventricle_distal | 6001 | 4.58 | 0.725 | 0.724 | 8.45 | 5.46 | 2.128 | 45.4 | 17.2 |

#### F10. One-scan-per-participant refit.

The main-text reference is fitted on all cognitively unimpaired scans, so participants with more visits carry more weight and the observations are not independent. Curves were refitted using only the earliest scan from each participant, giving one independent observation per person, and both samples were bootstrapped 200 times. The fitted curves are almost unchanged, so the reference is not distorted by repeated sampling of the same individuals. The confidence bands, however, are systematically wider in the one-scan-per-participant analysis than in the all-scan analysis.

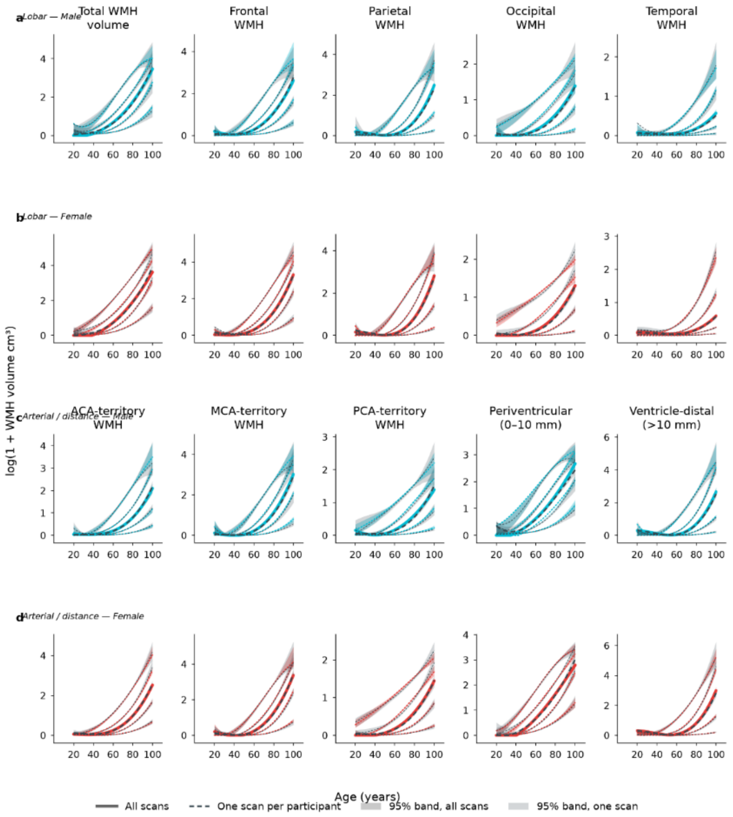

**Figure S2.F10 | Normative curves refitted using one scan per participant**. The main-text reference is fitted on 27,247 cognitively unimpaired scans from 13,303 participants, so repeated scans from the same person are not independent and participants with more visits carry more weight. Curves were therefore refitted using only the earliest scan from each participant, giving one independent observation per person, and both samples were bootstrapped 200 times. Colored lines and bands show the all-scan fit and its 95% bootstrap band; dark dashed lines and bands show the one-scan fit and its band. Solid and dashed lines mark the median; dotted lines the 5$^{th}$, 25$^{th}$, 75$^{th}$ and 95$^{th}$ percentiles. a, b, Lobar compartments in males and females. c, d, Arterial-territory and depth-distance compartments. Vertical scales differ between panels so that the bands remain visible. The fitted curves are almost unchanged: the median absolute difference between the two fits is 0.005–0.047 log units across compartments, under 5% in volume terms, so the reference is not distorted by repeated sampling of the same individuals. Confidence bands are wider in the one-scan-per-participant analysis, with a median width ratio of 1.33 (range 1.24–1.41 across compartments). Curve differences fitted curves and confidence band width comparisons are provided as separate CSV files: S2.T10a.csv, S2.T10b.csv and S2.T10c.csv, respectively.

#### F11. Transform sensitivity: log1p versus log(V + $1e^{-6}$).

The main-text reference curves are fitted on log(1 + V). The alternative log(V + $10^{-6}$) treats exact zero volumes very differently, mapping them far below the remainder of the distribution rather than to zero, and compartments containing many zeros could therefore be scored differently under the two. Curves were refitted under both transforms and every cognitively unimpaired scan scored against each. Percentile assignment is nearly unaffected overall; disagreement is concentrated in the compartments that contain exact zeros.

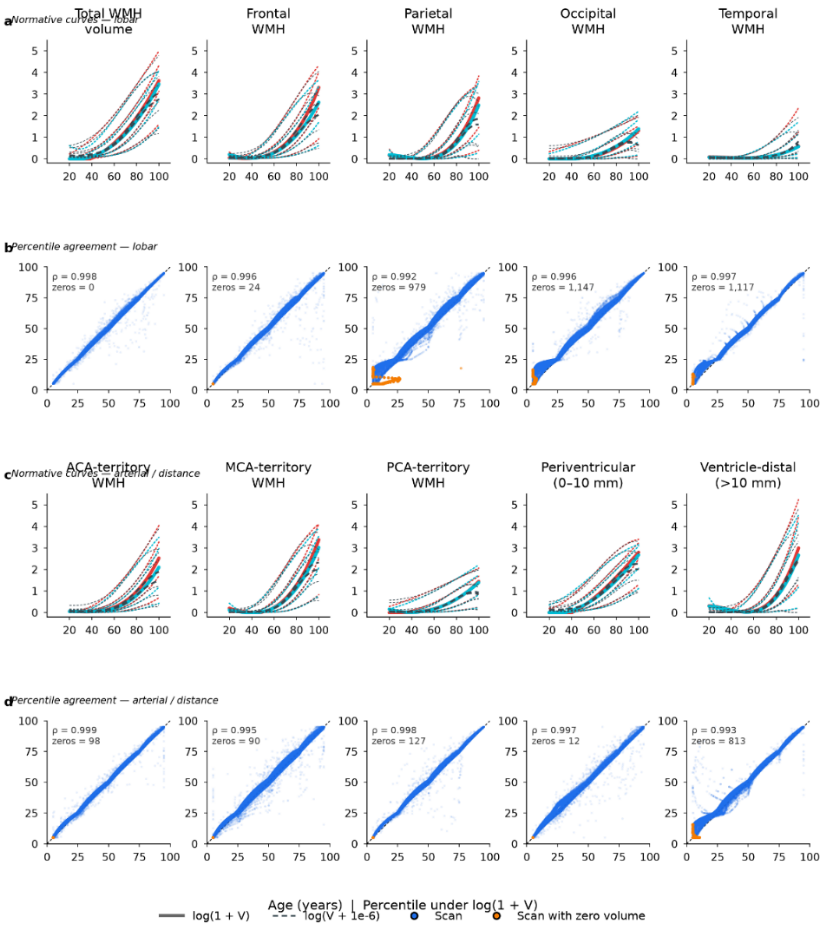

**Figure S2.F11 | Sensitivity of the normative reference to the choice of log transform.** The main-text reference curves are fitted on log(1 + V). The alternative log(V + $10^{-6}$) treats exact zero volumes very differently, mapping them to −13.8 rather than to zero and placing zero-volume scans far below the remainder of the distribution. Curves were refitted under both transforms with the specification of main-text Figures 1 and 2 and back-transformed to $cm^3$ for comparison. a, c, Fitted percentile curves for lobar and for arterial-territory and depth-distance compartments; colored lines, log(1 + V); dark dashed lines, log(V + $10^{-6}$). Solid and dashed lines mark the median, dotted lines the $5^{th}$, $25^{th}$, $75^{th}$ and $95^{th}$ percentiles. b, d, Percentile assigned to each cognitively unimpaired scan under each reference, with scans of exactly zero volume highlighted; the dashed line marks identical assignment. Percentile assignment is nearly unaffected by the choice of transform (Spearman ρ = 0.992–0.999; median absolute difference 0.6–1.8 percentile points). Disagreement is concentrated in compartments containing exact zeros: 15.8% of scans shift by more than five percentile points for parietal WMH (626 zero-volume scans) and 13.7% for occipital (804), compared with 0.8% for total WMH, which contains none. Summary statistics are given in Supplementary Table S2.T11a, shown below; full curve and percentile comparisons are provided as a separate file, S2.T11b.csv.

| compartment | n_zero | median_abs_diff_cm3 | max_abs_diff_cm3 | median_rel_diff | n_scans | spearman_rho | median_abs_pct_diff | pct_shift_gt5 | pct_shift_gt10 |
|---|---|---|---|---|---|---|---|---|---|
| ACA | 65 | 0.0222 | 14.2515 | 0.0887 | 27247 | 0.9985 | 0.88 | 0.7 | 0.2 |
| MCA | 61 | 0.0528 | 45.371 | 0.1483 | 27247 | 0.9952 | 1.39 | 3.9 | 0.8 |
| PCA | 86 | 0.0179 | 2.4428 | 0.0742 | 27247 | 0.9978 | 1.1 | 0.9 | 0.4 |
| frontal | 15 | 0.0437 | 33.6758 | 0.089 | 27247 | 0.9965 | 0.97 | 0.9 | 0.4 |
| occipital | 804 | 0.0197 | 3.6782 | 0.1598 | 27247 | 0.9964 | 1.34 | 13.7 | 3.0 |
| parietal | 626 | 0.0494 | 34.747 | 0.3315 | 27247 | 0.9925 | 1.65 | 15.8 | 3.5 |
| pv_total | 12 | 0.1065 | 9.9644 | 0.0638 | 27247 | 0.9968 | 0.95 | 4.5 | 0.4 |
| temporal | 707 | 0.0079 | 3.8678 | 0.1193 | 27247 | 0.9972 | 1.7 | 8.8 | 0.6 |
| total | 0 | 0.0923 | 30.1793 | 0.0534 | 27247 | 0.9984 | 0.6 | 0.8 | 0.3 |
| ventricle_distal | 510 | 0.054 | 65.3805 | 0.3707 | 27247 | 0.9929 | 1.77 | 11.8 | 1.4 |

#### F12. Sample-size convergence of the normative reference.

The reliability of a normative reference depends partly on sample size and whether the fitted curves remain stable as additional data become available. The cognitively unimpaired sample was subsampled at fractions from 2% to 75%, the curves refitted at each fraction with five replicates, and each refitted curve compared against the full-sample fit. Deviation was approximately consistent with inverse-square-root scaling across percentiles and age bands, providing a basis for model-based projections to larger sample sizes. The fitted scaling relationship suggests that additional scans would produce relatively small changes in the best-sampled age ranges, with deviation smallest between 60 and 80 years at the lower percentiles and largest at the extremes of the age range and in the upper tail, where the sample is thinnest. In contrast, leave-one-cohort-out validation (S2.8) reveals that the proportion of observations falling below the predicted median can deviate by up to 26.9 percentage points from the expected 50%. These findings suggest that cohort composition may be a more important constraint on reference calibration than further increases in sample size alone.

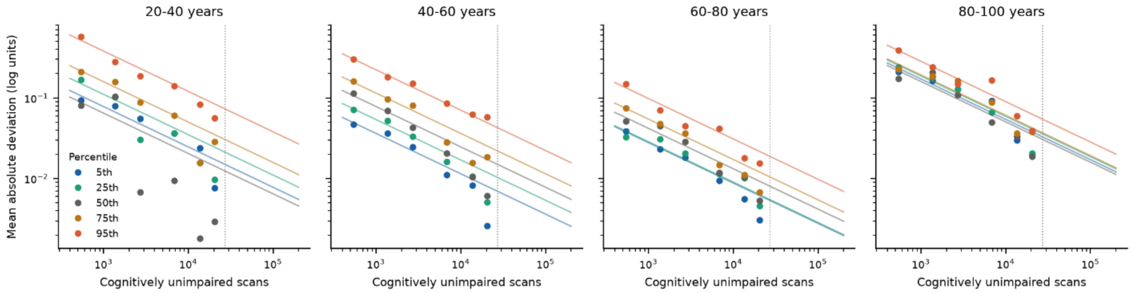

**Figure S2.F12 | Sample-size convergence of the normative reference.** The cognitively unimpaired sample was subsampled at fractions from 2% to 75%, the quantile regression curves refitted at each fraction with five replicates, and each refitted curve compared against the full-sample fit using the specification of main-text Figure 1. Points show the mean absolute deviation across replicates, by percentile and age band; lines show the fitted $1/\sqrt{n}$ relationship, and the dotted vertical line marks the current sample size of 27,247 scans. Both axes are logarithmic. Across the tested subsamples, deviations were approximately consistent with inverse-square-root scaling. At the current sample size, the fitted relationship predicts mean absolute deviations of 0.005–0.019 log units between 60 and 80 years, corresponding to 0.5–1.9% in volume terms. Predicted deviations are largest at the extremes of the age range and at the upper percentiles, reaching 0.073 log units (7.6%) at the $95^{\text{th}}$ percentile below 40 years and 0.031–0.055 log units (3.1–5.6%) above 80 years, where the sample is sparsest. Extrapolation of the fitted relationship to 200,000 scans predicts deviations of 0.002–0.027 log units. Fitted coefficients and projected deviations are given in Supplementary Table S2.T12a,b.

#### S3. WMH accumulation milestones

##### F1. Polynomial degree and smoothing sensitivity of growth-rate curves and milestone ages: quadratic versus cubic, σ = 2, 5, 8.

Growth rates are derivatives of the fitted median curves, so the polynomial degree in age determines the shape of the rate curves and influences the estimated milestone ages. Onset ages are determined by zero crossings, whereas crossover ages are determined by intersections between growth-rate curves. We therefore evaluated the milestone estimates across polynomial degrees and smoothing parameters. Curves were refitted for every combination of polynomial degree (quadratic, cubic) and Gaussian smoothing (σ = 2, 5, 8), and each milestone was re-estimated using 200 participant-clustered bootstrap replicates. The cubic places onsets several years earlier but estimates them far less precisely, with intervals two to three times wider, with greater variability in the sparsely sampled age range below 40 years; periventricular onset failed to resolve in almost every replicate under the cubic. Crossover ages, which fall in densely sampled age ranges, remain stable under both polynomial degrees. Smoothing had no material effect. Given the greater stability of onset estimates under the quadratic specification and the consistency of crossover ages across models, the quadratic specification was retained.

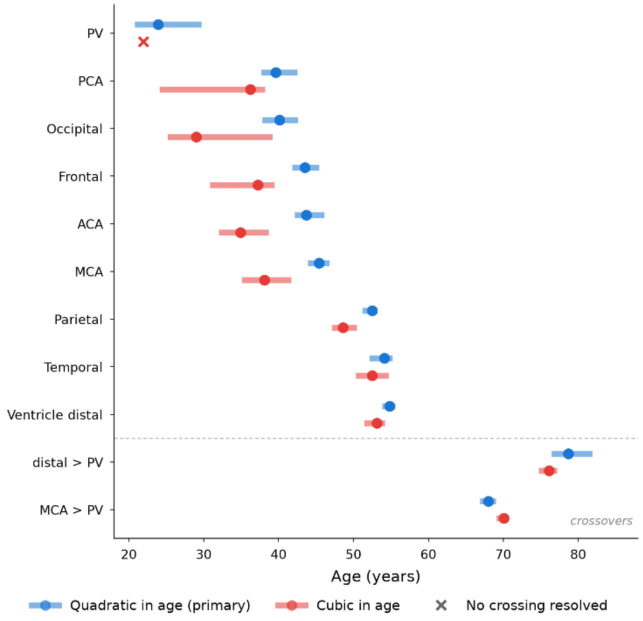

**Figure S3.F1 | Milestone ages are more precisely determined under a quadratic than a cubic model of age.** Growth rates are the derivative of the fitted median curve, so the polynomial degree influences the shape of the growth-rate curves and the estimated onset and crossover ages. Curves were refitted for every combination of degree (quadratic, cubic) and Gaussian smoothing (σ = 2, 5, 8), and each milestone re-estimated with a participant-clustered bootstrap of 200 replicates. Points show the estimated age and bars the 95% interval, at σ = 8. Onset ages, above the dashed line, are the ages at which each compartment's growth rate first becomes positive; crossover ages, below it, are the ages at which ventricle-distal and MCA-territory growth first exceed periventricular growth. The cubic places onsets 4 to 11 years earlier but estimates them far less precisely, with confidence intervals two to three times wider (occipital: 14.0 versus 4.8 years; PCA: 14.1 versus 4.8 years), consistent with the greater flexibility of the cubic model in the sparsely sampled age range below 40 years; periventricular onset failed to resolve in 199 of 200 replicates under the cubic. Crossover ages, which fall in densely sampled age ranges, are stable under both. Smoothing had no material effect: milestone ages varied by at most 0.3 years across $\sigma = 2$ to 8 for both degrees. The quadratic was therefore retained. Milestone ages under the primary specification are given in Supplementary Table S3.T1, shown below; full sensitivity results are provided as separate files, S3.T1a.csv and S3.T1b.csv.

| milestone | age | ci_lo | ci_hi | n_valid_replicates |
|---|---|---|---|---|
| onset_PV | 23.9 | 20.8 | 29.7 | 132 |
| onset_Ventricle_distal | 54.8 | 53.8 | 55.6 | 200 |
| onset_Frontal | 43.5 | 41.8 | 45.4 | 200 |
| onset_Parietal | 52.5 | 51.2 | 53.2 | 200 |
| onset_Occipital | 40.1 | 37.8 | 42.6 | 200 |
| onset_Temporal | 54.1 | 52.1 | 55.2 | 200 |
| onset_MCA | 45.4 | 43.9 | 46.8 | 200 |
| onset_PCA | 39.6 | 37.7 | 42.5 | 200 |
| onset_ACA | 43.7 | 42.1 | 46.1 | 200 |
| crossover_distal_over_PV | 78.7 | 76.4 | 81.9 | 200 |
| crossover_MCA_over_PV | 68.0 | 66.9 | 69.0 | 200 |

#### F2. Leave-one-cohort-out sensitivity of milestone ages

Milestone ages are single numbers derived from a pooled fit, so it matters whether anyone contributing study is driving them. Each study was removed in turn and the growth-rate curves and milestone ages re-estimated on the remaining eleven, using the primary specification. Most milestones vary by only a few years across folds. Occipital and PCA onsets are the exceptions; in both, the extreme fold is the removal of the study whose cognitively unimpaired participants carry the lowest WMH burden, with all other folds close to the full-sample value. Periventricular onset resolved in eight of twelve folds, reflecting accumulation already underway at the youngest ages sampled.

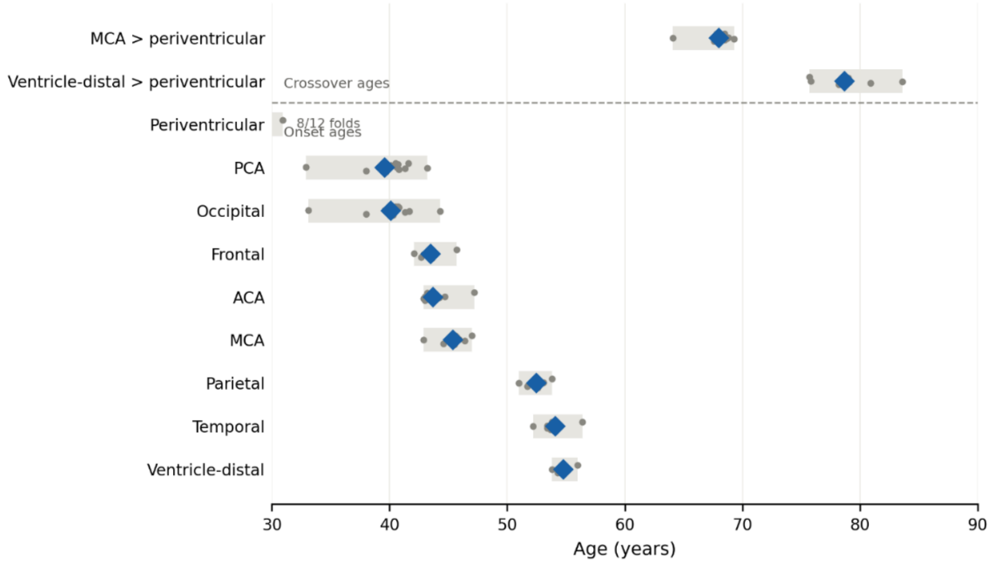

**Figure S3.F2 | Most milestone ages remain stable when individual studies are held out.** Each contributing study was removed in turn and the growth-rate curves and milestone ages re-estimated on the remaining eleven, using the primary specification. Diamonds show the full-sample estimate, grey points each leave-one-cohort-out fold, and the shaded bar the range across folds. Onset ages, above the dashed line, are the ages at which each compartment's growth rate first becomes positive; crossover ages, below it, are the ages at which ventricle-distal and MCA-territory growth first exceed periventricular growth. Most milestones vary little: ventricle-distal onset spans 2.2 years across folds, parietal 2.8, frontal 3.6 and MCA 4.1, and the two crossovers 5.2 and 7.9 years. Occipital and PCA onsets are the exceptions, spanning 11.2 and 10.3 years; in both, the extreme fold is the removal of HABS-HD, the cohort whose cognitively unimpaired participants carry the lowest WMH burden (Supplementary Fig. S2.F8), with all other folds within approximately two years of the full-sample value. Periventricular onset resolved in 8 of 12 folds, consistent with accumulation already underway at the youngest ages sampled; the fraction of folds resolved is annotated where fewer than all twelve.

The range of each milestone across folds is given in Supplementary Table S3.T2, shown below; milestone ages and growth rates by fold are provided as separate files, S3.T2b.csv and S3.T2c.csv.

| milestone | full_sample | fold_min | fold_max | range_yr | max_abs_dev | n_folds_resolved |
|---|---|---|---|---|---|---|
| crossover_MCA_over_PV | 68.0 | 64.1 | 69.3 | 5.2 | 3.9 | 12 |
| crossover_distal_over_PV | 78.7 | 75.7 | 83.6 | 7.9 | 4.9 | 12 |
| onset_ACA | 43.7 | 42.9 | 47.2 | 4.3 | 3.5 | 12 |
| onset_Frontal | 43.5 | 42.1 | 45.7 | 3.6 | 2.2 | 12 |
| onset_MCA | 45.4 | 42.9 | 47.0 | 4.1 | 2.5 | 12 |
| onset_Occipital | 40.1 | 33.1 | 44.3 | 11.2 | 7.0 | 12 |
| onset_PCA | 39.6 | 32.9 | 43.2 | 10.3 | 6.7 | 12 |
| onset_PV | 23.9 | 23.4 | 30.9 | 7.5 | 7.0 | 8 |
| onset_Parietal | 52.5 | 51.0 | 53.8 | 2.8 | 1.5 | 12 |
| onset_Temporal | 54.1 | 52.2 | 56.4 | 4.2 | 2.3 | 12 |
| onset_Ventricle_distal | 54.8 | 53.8 | 56.0 | 2.2 | 1.2 | 12 |

#### F3. Derived spatial indices.

The compartment-specific trajectories describe where WMH accumulates but not how the overall spatial pattern changes shape. Five indices summarise that redistribution as single quantities: three ratios contrasting anterior against posterior, periventricular against ventricle-distal, and MCA against PCA territory; a concentration index quantifying spatial unevenness, with higher values indicating greater concentration; and the proportion of burden lying in MCA territory. Trajectories were fitted by median quantile regression with a degree 4 polynomial in age and 200 bootstrap replicates, separately in each diagnostic group, allowing diagnostic-group differences to be evaluated relative to the cognitively unimpaired trajectory.

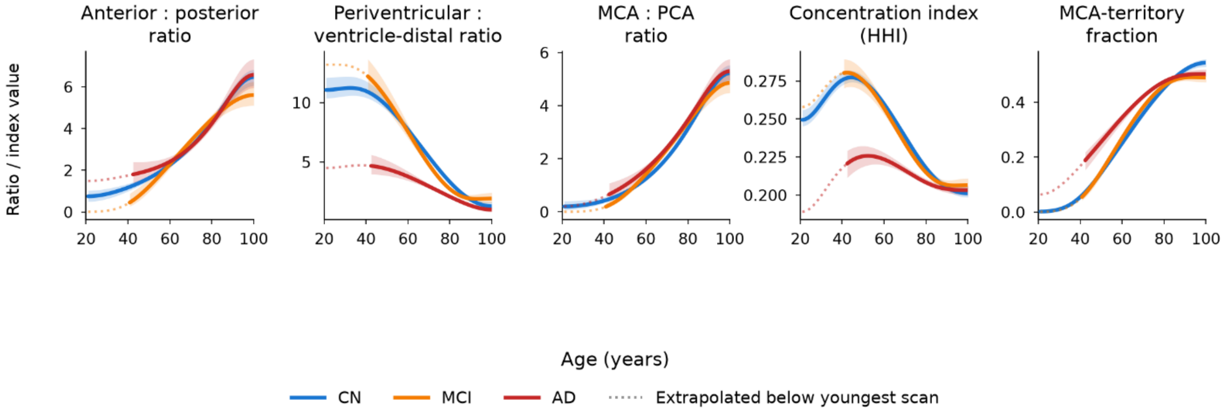

**Figure S3.F3 | Spatial redistribution of WMH with age in cognitively unimpaired, MCI and AD participants.** Main-text Figure 3 shows these indices in cognitively unimpaired participants only. Trajectories were fitted by median quantile regression with a degree 4 polynomial in age and 200 bootstrap replicates, separately in each diagnostic group. Solid lines with shaded 95% intervals span the observed age range for each group; dotted lines show model extrapolation below the youngest scan available (CN 21 years, MCI 41, AD 42). Sampling is uneven across the age range: in the fifth decade the groups contribute 189, 10 and 11 scans respectively, rising to 2,560, 492 and 251 in the sixth. The first three panels are ratios, so values above 1 indicate that the first-named region carries the greater share of lesion volume; the concentration index ranges from 1/7 = 0.14 when burden is spread evenly across the seven compartments to 1 when confined to one; the final panel is the proportion of total burden lying in MCA territory. All three groups exhibit broadly similar age-related spatial redistribution: from posterior to anterior, from periventricular toward ventricle-distal, and from PCA toward MCA territory. AD differs from cognitively unimpaired participants on three of the five measures: the periventricular:ventricle-distal ratio is lower at every decade (4.4 versus 9.7 at 50 years, non-overlapping intervals), and the concentration index and MCA-territory fraction differ through the seventh decade, with differences diminishing at older ages. The anterior:posterior and MCA:PCA ratios do not differ at any age, and MCI trajectories show no clear separation from cognitively unimpaired trajectories across the observed age range. The lower periventricular:ventricle-distal ratio in AD does not indicate reduced periventricular burden: median volumes at 70 years are higher in AD in both compartments (periventricular 3.83 versus 2.69 cm³; ventricle-distal 1.59 versus 0.58 cm³), so the ratio falls because the ventricle-distal excess is proportionally larger. Index values by decade and group are given in Supplementary Table S3.T3, provided as separate files, S3.T3a.csv and S3.T3b.csv.

#### F4. Age-prevalence of vascular, molecular and clinical anchors at the WMH milestone ages

The milestone ages describe when fitted regional WMH growth rates first become positive and when the fastest-growing compartment changes, but they carry no information about what else is happening at those ages. To place them in a biological and clinical context, we estimated the age-prevalence of seven measures available in different subsets of the study sample, grouped by the biological or clinical domain they represent: vascular risk factors (hypertension, diabetes), cerebrovascular disease (stroke), AD molecular pathology (amyloid positivity, tau Braak stage 3 or above) and clinical expression (mild cognitive impairment, Alzheimer's dementia). Diagnosis was taken from the scan-matched record rather than the participant's most recent diagnosis, so that a participant who converted later is not counted as impaired at earlier ages. Two features of the sample constrain what can be estimated. First, the anchors are measured in different subsets: vascular measures are available in 9,141 participants, amyloid in 10,070, tau in 5,284 and diagnosis in 19,463. Second, the studies contributing at each age differ, so a probability estimated by averaging across whichever studies happen to have data near a given age changes as that set changes. We therefore standardized each anchor over a fixed set of studies, selected as those with at least five participants within ±5 years of every milestone age that anchor could report. This holds cohort composition constant along the age axis, at the cost of restricting each anchor to the studies with the widest age coverage. The locally varying alternative is reported alongside as a sensitivity analysis. Because the fixed sets differ between anchors, the estimates describe how each anchor changes across the milestone ages, not how anchors compare with one another, and they are not population prevalence estimates.

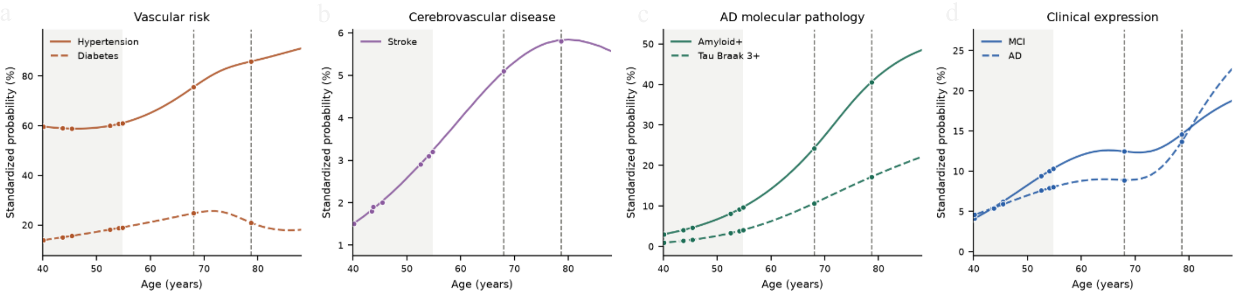

**Figure S3.F4 | Age-prevalence of vascular, molecular and clinical anchors across the WMH accumulation milestones.** Each anchor was modelled as a smooth function of age by generalized estimating equations with an exchangeable working correlation clustered on participant, adjusting for sex and contributing study, with spline complexity selected per anchor by QICu. Curves show probabilities standardized over a fixed set of contributing studies, chosen as those with at least five participants within ±5 years of every reportable milestone age, so that cohort composition is held constant across the age range; points mark the milestone ages. The shaded band spans the regional onset milestones (39.6–54.8 years) and dashed lines mark the two crossover milestones (68.0 and 78.7 years). a, Vascular risk (HABS-HD, NACC). b, Cerebrovascular disease (HABS-HD, NACC). c, AD molecular pathology (amyloid: HABS-HD, KBASE, Knight ADRC, NACC, WRAP; tau: HABS-HD, Knight ADRC, NACC). d, Clinical expression (KBASE, Knight ADRC, NACC, WRAP). Fixed cohort sets differ between anchors because data availability differs, so probabilities represent standardized estimates within the contributing studies and should not be interpreted as population prevalence or compared across anchors. Vertical scales differ between panels. The apparent convergence of MCI and AD probabilities above 80 years reflects equal cohort weighting applied to a small Knight ADRC subsample in that age range; unstandardized prevalence within the same cohorts shows MCI exceeding AD at every age band (Supplementary Table S3.T4). Estimates above 85 years rest on fewer than 600 participants for every anchor.

**Supplementary Table S3.T4 | Age-prevalence of vascular, molecular and clinical anchors at the WMH accumulation milestones**. Each anchor was modelled by generalized estimating equations with a binomial family, logit link and exchangeable working correlation clustered on participant, with a natural cubic spline in age, sex, and contributing study entered with sum-to-zero coding. Spline complexity was selected per anchor from 3, 4 and 5 degrees of freedom as the lowest value whose QICu fell within 2 of the minimum. Cohorts contributing no positive cases for an anchor were excluded from that model. Confidence intervals were obtained by 2,000 draws from the robust coefficient covariance. a, Descriptive prevalence by decade, unadjusted. b, QICu across all three spline specifications for every anchor, with the selected model indicated. c, Standardized probability at each milestone age under both the fixed and the locally varying cohort set, with the contributing studies listed. d, Standardized probability across a one-year age grid. e, Unadjusted prevalence within the fixed cohort set by age band, with participant and cohort counts. Provided as separate files, S3.T4a.csv to S3.T4e.csv.

| **Anchor** | **45.4 yr** | **52.5 yr** | **68.0 yr** | **78.7 yr** |
|---|---|---|---|---|
| AD | 5.9 (2.9-10.9) | 7.6 (4.2-13.3) | 8.9 (5.4-14.7) | 13.7 (8.6-21.1) |
| Amyloid+ | 4.7 (2.8-7.7) | 8.1 (6.4-10.3) | 24.2 (22.4-26.3) | 40.6 (38.1-43.2) |
| Diabetes | 15.6 (11.1-21.4) | 18.1 (15.9-20.8) | 24.7 (23.2-26.2) | 20.9 (19.3-22.6) |
| Hypertension | 58.8 (50.5-67.4) | 59.9 (56.6-63.6) | 75.5 (74.0-76.9) | 85.8 (84.5-87.1) |
| MCI | 6.2 (4.9-7.8) | 9.4 (8.2-10.8) | 12.5 (11.4-13.6) | 14.6 (13.3-16.0) |
| Stroke | 2.0 (1.3-3.1) | 2.9 (2.3-3.8) | 5.1 (4.6-5.7) | 5.8 (5.1-6.7) |
| Tau Braak 3+ | 1.7 (0.6-4.9) | 3.3 (1.3-8.1) | 10.6 (4.9-21.2) | 17.2 (8.4-30.9) |

#### S4. Burden percentile scoring

##### F1. Choice of spline bases for the Gaussian-family GAM comparator.

The GAM comparator constructs percentile curves from a mean and a variance, each fitted as a smooth function of age, and the number of basis functions controls how flexible those smooths are. Too few and the curves cannot follow the age trend; too many and they oscillate, producing percentile curves that fall with increasing age. A normative reference in which expected burden decreases with age is not interpretable, so the basis dimension was set to the highest value at which all seven fitted percentile curves remained non-decreasing across the observed age range, tested separately in men and women. The complete search across mean and variance bases is reported. Men are the binding constraint, and eight bases for the mean with four for the variance is the most flexible admissible setting; the violations at higher settings are numerically small and occur at the boundaries of the age range, reflecting a deliberately strict criterion rather than gross misfit.

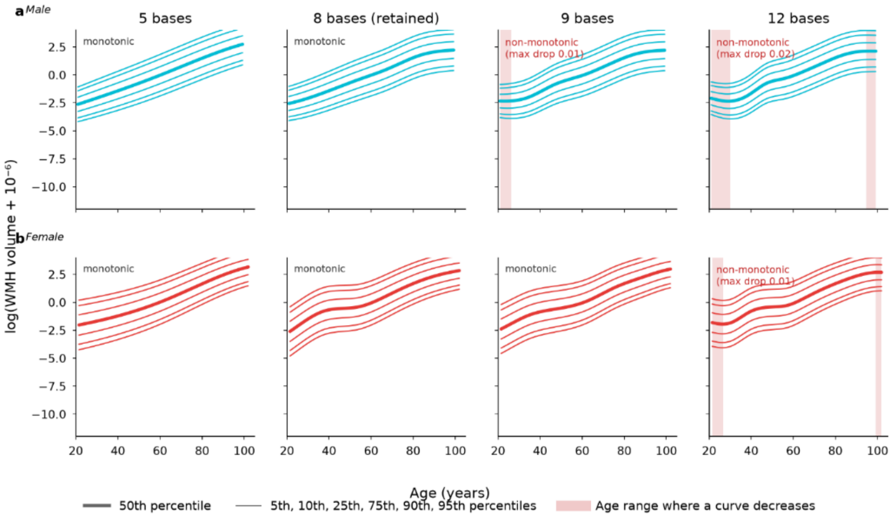

**Figure S4.F1 | Selection of the number of spline bases for the Gaussian family GAM comparator.** Percentile curves from the generalized additive model are constructed from a mean and a variance, each fitted as a smooth function of age, and are then read off a Gaussian conditional distribution. The number of basis functions controls how flexible those smooths are: too few and the curves cannot follow the age trend, too many and they begin to oscillate, producing percentile curves that fall with increasing age. A normative reference in which expected burden decreases with age is not interpretable, so the number of bases was set to the highest value at which all seven fitted percentile curves remained non-decreasing across the observed age range, tested separately in men (a, n = 10,030, ages 21–99) and women (b, n = 17,217, ages 22–102). Panels show the fitted curves at four settings for the mean, with the variance held at four bases throughout; thick lines mark the median and thin lines the $5^{th}$, $10^{th}$, $25^{th}$, $75^{th}$, $90^{th}$ and $95^{th}$ percentiles. Shaded bands mark age ranges in which at least one curve decreases, with the largest single decrease annotated. At five and eight bases every curve is non-decreasing in both sexes. At nine bases the male curves develop a small decrease near the youngest ages, while the female curves remain admissible; at twelve bases both sexes fail, at the youngest and oldest ages where the sample is sparsest. Men are therefore the binding constraint, and eight bases for the mean with four for the variance is the most flexible admissible setting. The violations are numerically small, with the largest being 0.02 log units, reflecting a deliberately strict criterion applied at the boundaries of the age range rather than substantial model misfit. The complete search across mean (3–20) and variance (3–10) bases is given in Supplementary Table S4.T1, provided as a separate file, S4.T1.csv.

#### F2. Concordance between quantile regression and GAM percentile assignments.

Having established the admissible GAM specification, the two references were fitted on the same outcome and structure and every cognitively unimpaired scan scored against both, to establish whether the choice of framework changes an individual's percentile. Assignments are closely concordant, with a median absolute difference under one percentile point and few scans differing by more than five. The residual difference is systematic rather than random: the GAM places scans marginally higher through the middle of the distribution and marginally lower above approximately the 75th percentile, consistent with differences between the Gaussian conditional model and the distribution-free QR reference.

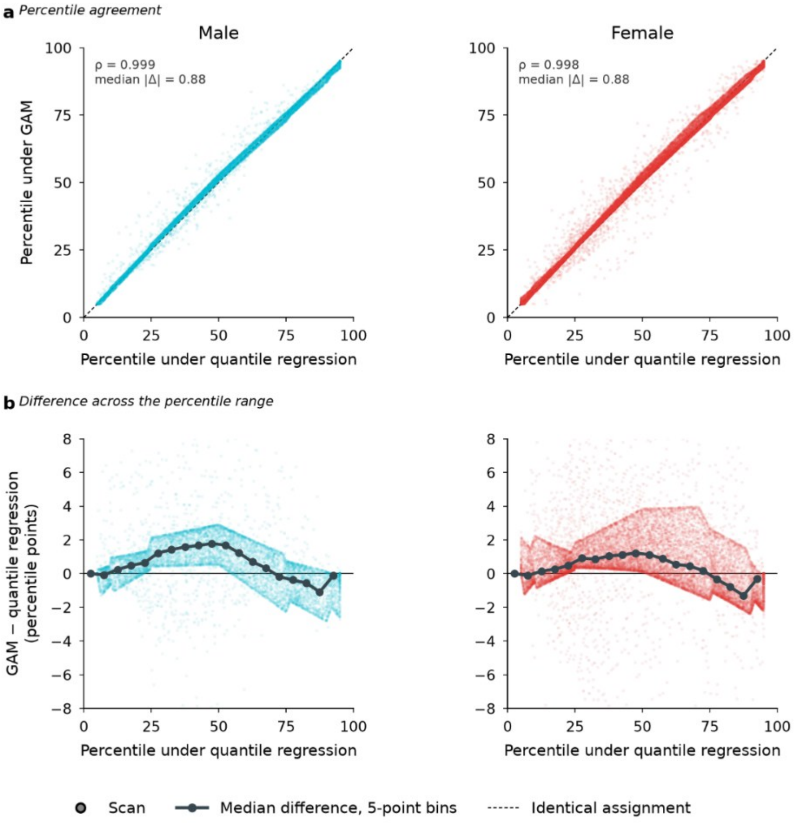

**Figure S4.F2 | Agreement between quantile regression and Gaussian-family GAM percentile assignments.** Both references were fitted on the cognitively unimpaired sample using the same outcome, $\log(V + 10^{-6})$ of harmonized total WMH volume, sex-stratified and without cohort terms so that the two models see the same structure; the GAM used eight basis functions for the mean and four for the variance (Supplementary Fig. S4.F1). Every cognitively unimpaired scan was then scored against both. a, Percentile assigned under each reference, one point per scan; the dashed line marks identical assignment. b, Difference between the two assignments plotted against the quantile regression percentile, with the median in five-point bins. Assignments are closely concordant overall: Spearman $\rho$ = 0.999 in men and 0.998 in women, median absolute difference 0.88 percentile points in both, with 1.7% and 2.4% of scans differing by more than five points. The residual difference is systematic rather than random. The GAM places scans marginally higher than quantile regression through the middle of the distribution, by up to 1.8 percentile points near the median, and marginally lower above approximately the 75th percentile, reaching −1.2 points by the 90th. This pattern is consistent with differences between the Gaussian conditional model and the distribution-free QR reference.

Summary statistics by percentile band are given in Supplementary Table S4.T2a, shown below; scan-level assignments are provided as a separate file, S4.T2b.csv.

| sex | band | n | spearman | pearson | median_abs_diff | mean_signed_diff | pct_gt5 | pct_gt10 |
|---|---|---|---|---|---|---|---|---|
| Female | all | 17836 | 0.9976 | 0.9973 | 0.88 | 0.54 | 2.4 | 0.6 |
| Female | 0-25 | 4458 | | | 0.44 | 0.45 | 1.6 | 0.4 |
| Female | 25-50 | 4472 | | | 1.17 | 1.26 | 4.4 | 1.3 |
| Female | 50-75 | 4447 | | | 1.06 | 0.96 | 2.5 | 0.7 |
| Female | 75-90 | 2677 | | | 1.21 | -0.34 | 1.5 | 0.2 |
| Female | 90-100 | 1782 | | | 0.43 | -0.71 | 0.3 | 0.0 |
| Male | all | 10030 | 0.9985 | 0.9981 | 0.88 | 0.5 | 1.7 | 0.4 |
| Male | 0-25 | 2503 | | | 0.39 | 0.38 | 1.6 | 0.5 |
| Male | 25-50 | 2505 | | | 1.67 | 1.52 | 2.9 | 0.4 |
| Male | 50-75 | 2512 | | | 1.16 | 0.8 | 2.0 | 0.7 |
| Male | 75-90 | 1504 | | | 0.73 | -0.72 | 0.2 | 0.0 |
| Male | 90-100 | 1006 | | | 0.13 | -0.62 | 0.0 | 0.0 |

#### F3. Associations between burden percentile and cognitive, biomarker and vascular measures.

A percentile is a descriptive position within a reference distribution and carries no meaning on its own; its value depends on whether it relates to measures of brain health. Every scan was assigned a burden percentile against the pooled cognitively unimpaired reference and related to the cognitive, biomarker and vascular measures available in the dataset, with associations reported overall, by diagnostic group, and with and without cohort terms in the reference model. Higher burden percentile is associated with poorer performance in all four cognitive domains and with higher values on every biomarker and vascular measure examined. Effect sizes are small, consistent with WMH burden being one of several contributors to cognitive and structural variation.

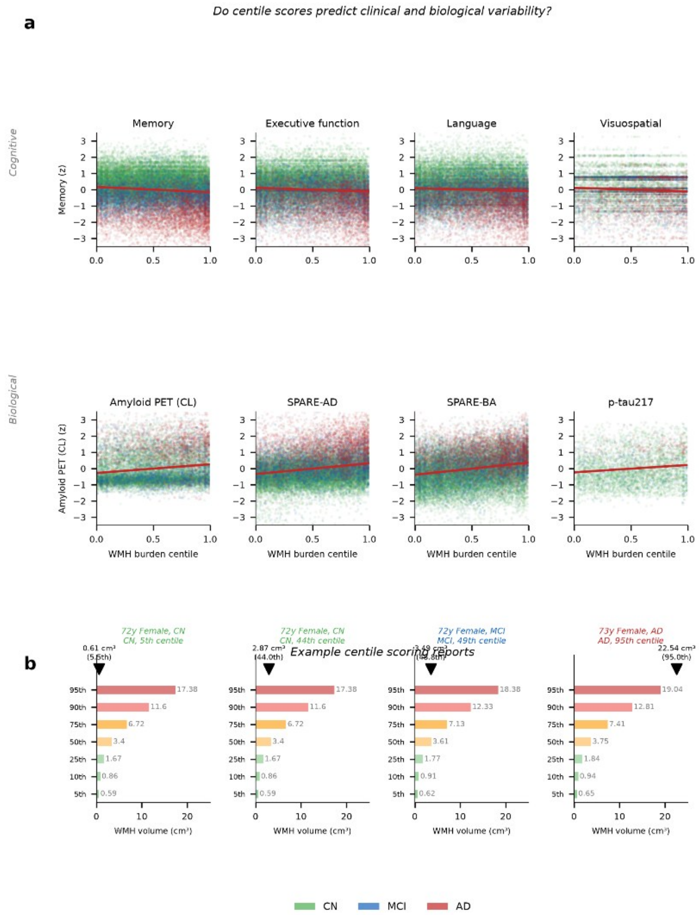

**Figure S4.F3 | Associations between burden percentile and cognitive, biomarker and vascular measures, and single-scan interpretation.** a, Every scan was assigned a burden percentile against the cognitively unimpaired reference (quantile regression on $\log(V + 10^{-6})$ of harmonized total WMH volume, age and age squared, sex-stratified, with contributing study included) and related to available cognitive, biomarker and vascular measures, as partial correlations adjusted for age and sex; correlations involving binary measures (hypertension, smoking) are point-biserial and their magnitude is bounded by the prevalence of the measure. Higher burden percentile was associated with poorer performance in all four cognitive domains — memory ($r = -0.134$, 95% CI −0.143 to −0.124, $n = 38{,}358$), executive function (−0.125, $n = 31{,}392$), language (−0.121, $n = 31{,}387$) and visuospatial ability (−0.096, $n = 12{,}114$) — and with higher values on every biomarker and vascular measure examined: SPARE-BA ($r = 0.208$, $n = 29{,}130$),

SPARE-AD (0.183, n = 29,130), plasma p-tau217 (0.139, n = 4,849), hypertension (0.115, n = 9,059), amyloid PET Centiloids (0.094, n = 16,860), ASCVD risk score (0.093, n = 8,809), systolic blood pressure (0.072, n = 8,875) and smoking (0.053, n = 9,057). All associations were significant at $P < 10^{-6}$. Correlations range from 0.05 to 0.21 in absolute value, corresponding to 0.3% to 4.3% of shared variance, consistent with WMH burden being one of several contributors to cognitive and structural variation. Direction and significance are unchanged when the percentile is constructed without study terms, when study is adjusted for in the association model, and when correlations are estimated within each study separately (Supplementary Table S4.T4). b, Interpretation of an individual scan against the exported reference curves, showing reference WMH volumes at the 5th, 25th, 50th, 75th and 90th percentiles for a given age and sex, and the percentile assigned to an observed volume. Correlations, confidence intervals and sample sizes are given in Supplementary Table S4.T3, shown below; associations by diagnostic group and the effect of cohort terms are provided as separate files, S4.T3b.csv and S4.T3c.csv.

| outcome | r | ci_lo | ci_hi | p | n |
|---|---|---|---|---|---|
| Memory | -0.1336 | -0.1434 | -0.1237 | 3.70e-152 | 38358 |
| Executive | -0.125 | -0.1356 | -0.1143 | 1.90e-109 | 31392 |
| Language | -0.1212 | -0.1317 | -0.1105 | 4.85e-103 | 31387 |
| Visuospatial | -0.0963 | -0.1146 | -0.0786 | 2.29e-26 | 12114 |
| Amyloid PET | 0.0936 | 0.0788 | 0.1081 | 3.77e-34 | 16860 |
| SPARE-AD | 0.1825 | 0.172 | 0.1936 | 1.75e-216 | 29130 |
| SPARE-BA | 0.2079 | 0.1975 | 0.219 | 6.06e-282 | 29130 |
| p-tau217 | 0.1387 | 0.1101 | 0.1657 | 2.86e-22 | 4849 |
| ASCVD risk | 0.0927 | 0.0714 | 0.1127 | 2.88e-18 | 8809 |
| Smoking | 0.0533 | 0.0331 | 0.074 | 3.89e-07 | 9057 |
| Hypertension | 0.1151 | 0.0943 | 0.1357 | 4.27e-28 | 9059 |
| Systolic BP | 0.0724 | 0.0503 | 0.0927 | 8.69e-12 | 8875 |

#### F4. Sensitivity of burden associations to cohort handling.

The burden percentile can be constructed in two ways, and they answer slightly different questions. A pooled reference is fitted on all cognitively unimpaired scans with no study indicators, so every scan is compared against one common curve; this is what allows a scan from outside the contributing studies to be scored. A cohort-adjusted reference includes an indicator for each study, so a scan is compared against others in its own study. Studies differ in the WMH burden of their cognitively unimpaired participants by up to 27 percentile points, so the choice affects individual percentiles and therefore the associations computed from them. Each association was therefore estimated under both references, crossed with four adjustment sets in the association model: age and sex; age, sex and contributing study; age, sex and log WMH volume; and all four terms together. Associations retain their direction under every specification. Their magnitude is sensitive to two choices: for the cognitive outcomes, scoring against the pooled reference without adjusting the association model for study weakens the association, and adjusting for log WMH volume distinguishes the persistent cognitive associations from the substantially attenuated structural associations.

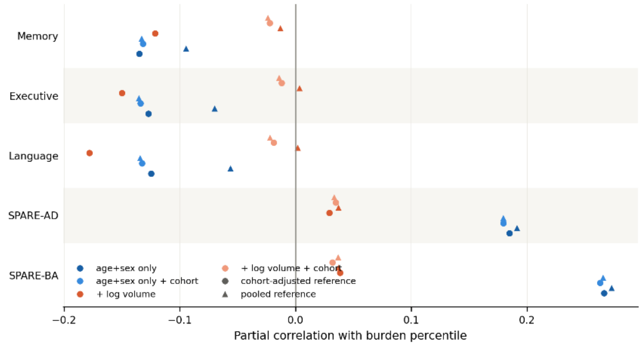

**Figure S4.F4 | Sensitivity of burden associations to cohort handling.** The burden percentile can be scored against a cohort-adjusted reference, which includes an indicator for each contributing study, or against a pooled reference fitted without study indicators; the pooled version is what allows a scan from outside the contributing studies to be scored. Studies differ substantially in the WMH burden of their cognitively unimpaired participants, so the choice affects individual percentiles and therefore the associations computed from them. Each association was estimated under both references, crossed with four adjustment sets in the association model: age and sex; age, sex and contributing study; age, sex and log WMH volume; and all four terms together. Marker shape distinguishes the reference and colour the adjustment set. Associations retain their direction under every specification, but their magnitude depends on the specification in two distinct ways. First, for the three cognitive outcomes the pooled reference adjusted only for age and sex gives weaker associations than the cohort-adjusted reference (memory −0.095 against −0.135; executive function −0.069 against −0.127), and adding contributing study to the association model recovers the difference. Second, adjusting for log WMH volume separates the cognitive outcomes from the structural indices: cognitive associations persist and in some cases strengthen under the cohort-adjusted reference, whereas SPARE-AD and SPARE-BA fall from approximately +0.18 and +0.26 to below +0.05. This adjustment should be read with care, since the burden percentile is itself a function of volume, age and sex, so conditioning on volume removes much of the variable's own content.

**Supplementary Table S4.T4 | Sensitivity of burden associations to cohort handling.** Partial correlations between burden percentile and each cognitive, biomarker and vascular measure, evaluated under two percentile references (cohort-adjusted and pooled), each crossed with four adjustment sets: age and sex; age, sex and contributing study; age, sex and log WMH volume; and age, sex, contributing study and log WMH volume. Columns give the specification, sample size, number of contributing studies, correlation, 95% confidence interval by Fisher z transformation, and P value. Within-study estimates combine studies contributing at least 200 scans for that outcome, weighted by sample size; no P value is reported for the combined estimate. Provided as a separate file, S4.T4.csv.

#### S5. Voxelwise topographic reference modelling (vTRM)

##### F1. Consistency of the population Frequency maps.

The vTRM score uses frequency maps derived from cognitively unimpaired participants, making it important to establish that WMH frequency maps represent a spatial pattern shared across contributing studies rather than one dominated by any single cohort. A frequency map records, for every voxel, the proportion of contributing scans with a lesion at that location, and two maps can therefore be compared by correlating their values across voxels; a high correlation meaning the same regions are commonly affected in both, irrespective of how much lesion burden each group carries overall. Each study's map was correlated against the map built from the full analytic sample, male and female maps were correlated within age band, and a map built from one scan per participant was compared against the full-sample map. The spatial distribution is closely shared across studies even where total burden and peak frequency differ substantially, the sexes show near-identical spatial patterns from the sixth decade onward, and repeated imaging of the same participants does not materially alter the spatial patterns.

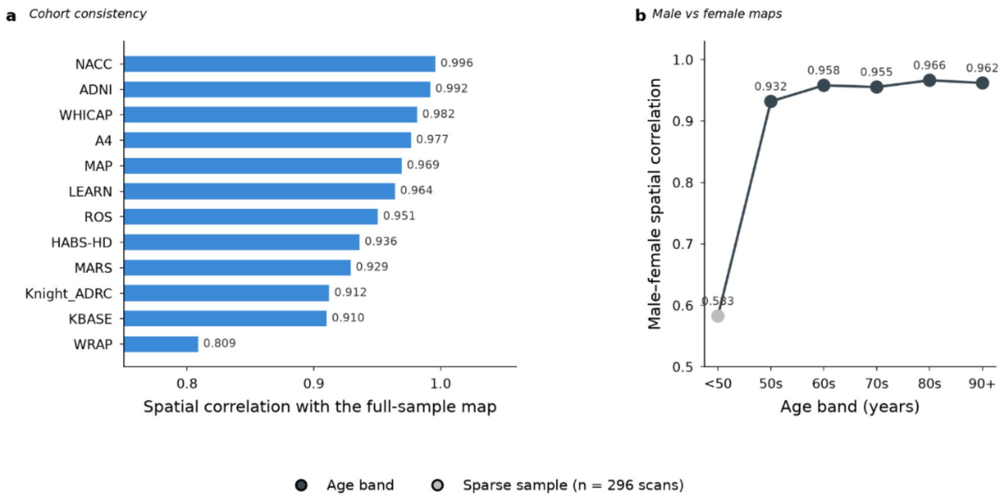

**Figure S5.F1 | Consistency of the population WMH frequency maps.** A frequency map records, for every voxel in MNI152 2 mm space, the proportion of contributing scans with a lesion at that location. Two maps can therefore be compared by correlating their values across voxels: a high correlation means the same regions are commonly affected in both, irrespective of how much lesion burden each group carries overall. a, Each contributing study's frequency map correlated against the map built from the full analytic sample, over the 165,018 voxels lesioned in at least one scan. Correlations range from 0.809 to 0.996, so the spatial distribution of WMH is closely shared across studies even though total burden and peak frequency differ substantially between them (peak frequency 17.8% in Knight ADRC to 46.2% in MARS). The lowest value, WRAP, is the youngest contributing cohort, in which lesions remain concentrated close to the ventricles and the more distributed pattern seen at older ages has not yet emerged. b, Male and female frequency maps correlated within each age band. From the sixth decade onward the two sexes show near-identical spatial patterns (r = 0.932 to 0.966), indicating that where WMH occur does not differ appreciably by sex even though women reach higher peak frequencies at every age. The value below 50 years is not interpretable: only 296 scans contribute to that band, and the male map contains 8,654 non-zero voxels against 20,557 in the female map, so the two are compared over largely non-overlapping territory. Repeated imaging of the same participants does not shape the maps: a map built from one scan per participant is near-identical to the full-sample map (Pearson r = 0.998,

Spearman ρ = 0.969), with peak frequency 23.8% against 25.2% and 151,996 against 165,018 non-zero voxels. Cohort correlations, sex correlations by age band, the repeated-scan comparison and peak frequency by age band are given in Supplementary Table S5.T1, shown below; mean frequency by anatomical compartment for each study is provided as a separate file, S5.T1c.csv.

**a Cohort consistency**

| cohort | pearson_r | spearman_rho | n_voxels_compared | peak_pct | nonzero_voxels |
|---|---|---|---|---|---|
| NACC | 0.996 | 0.9318 | 165018 | 23.2 | 143536 |
| ADNI | 0.9919 | 0.8662 | 165018 | 28.0 | 98154 |
| WHICAP | 0.9816 | 0.8086 | 165018 | 39.1 | 81486 |
| A4 | 0.9767 | 0.8095 | 165018 | 34.6 | 81692 |
| MAP | 0.9694 | 0.8174 | 165018 | 41.8 | 83321 |
| LEARN | 0.9641 | 0.7054 | 165018 | 34.8 | 48704 |
| ROS | 0.9506 | 0.6371 | 165018 | 42.4 | 34487 |
| HABS-HD | 0.9361 | 0.7972 | 165018 | 22.9 | 87171 |
| MARS | 0.9291 | 0.7234 | 165018 | 46.2 | 54928 |
| Knight_ADRC | 0.9122 | 0.6822 | 165018 | 17.8 | 44880 |
| KBASE | 0.9102 | 0.6632 | 165018 | 23.2 | 43605 |
| WRAP | 0.8088 | 0.6614 | 165018 | 30.8 | 54866 |

**b Sex comparison by age band**

| age_band | pearson_r | spearman_rho | n_voxels_compared | peak_male_pct | peak_female_pct | nonzero_male | nonzero_female | mean_diff_pct | peak_both_pct |
|---|---|---|---|---|---|---|---|---|---|
| lt50 | 0.5827 | -0.027 | 24223 | 16.7 | 19.9 | 8654 | 20557 | 0.505 | 15.2 |
| 50s | 0.9319 | 0.3869 | 69742 | 14.7 | 18.0 | 42121 | 57258 | 0.008 | 15.9 |
| 60s | 0.958 | 0.6937 | 118611 | 20.3 | 23.7 | 85351 | 104488 | 0.026 | 21.7 |
| 70s | 0.9553 | 0.7876 | 140240 | 25.7 | 34.6 | 108490 | 126808 | 0.064 | 29.6 |
| 80s | 0.9663 | 0.7592 | 122909 | 41.7 | 46.8 | 95067 | 107586 | 0.213 | 42.4 |
| 90plus | 0.9619 | 0.6846 | 70923 | 52.7 | 56.7 | 44154 | 63479 | 1.122 | 54.2 |

**d One-scan-per-participant comparison**

| comparison | pearson_r | spearman_rho | max_freq_all | max_freq_baseline | nonzero_all | nonzero_baseline |
|---|---|---|---|---|---|---|
| all_scans_vs_baseline_only | 0.9984 | 0.9694 | 25.2 | 23.8 | 165018 | 151996 |

**e Peak frequency by age band**

| age_band | n_scans | n_male | n_female | peak_frequency_pct |
|---|---|---|---|---|
| lt50 | 296 | 120 | 176 | 15.2 |
| 50s | 3456 | 1139 | 2317 | 15.9 |
| 60s | 11702 | 4339 | 7363 | 21.7 |
| 70s | 18116 | 7513 | 10603 | 29.6 |
| 80s | 7126 | 3040 | 4086 | 42.4 |
| 90plus | 742 | 275 | 467 | 54.2 |

#### F2. Construct checks on the vTRM score: self-inclusion, choice of summary statistic, and repeatability.

We evaluated three properties of the vTRM score: self-inclusion bias, sensitivity to the summary statistic, and within-participant reproducibility. First, every cognitively unimpaired scan contributes to the frequency maps it is then scored against, raising the frequency at the voxels it occupies; because the maps store lesion counts divided by contributing scans, the value with that scan removed can be computed exactly. Second, the score begins as the mean reference frequency across a scan's lesion voxels, and a mean can be pulled down by a few voxels in very rare locations, so three alternative summaries were tested against the same burden-and-age adjustment. Third, because the score is what remains after burden and age are removed, its reproducibility within a person should be established rather than assumed and was measured as the intraclass correlation among participants with repeat scans. The self-inclusion effect is negligible, the mean is the summary the adjustment explains best, and the score reaches an intraclass correlation close to that obtained for directly measured WMH volume.

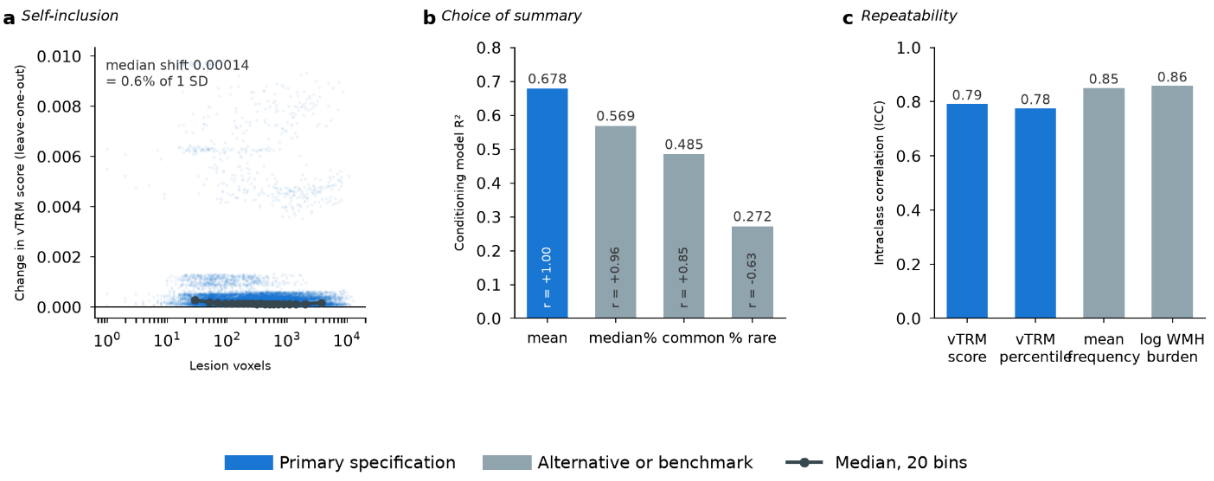

**Figure S5.F2 | Construct checks on the vTRM score.** Three checks on how the vTRM score is built, all in the cognitively unimpaired reference sample. a, Does a scan inflate its own score? Every cognitively unimpaired scan helped build the frequency maps it is then scored against, so each one raises the frequency at the voxels it occupies. The maps store lesion counts divided by the number of contributing scans, so the value with that scan removed can be calculated exactly. The panel shows how much each vTRM score changes when this is done, against the size of the lesion mask; the dark line is the median in twenty bins. The typical change is $1.4 \times 10^{-4}$ score units, about 0.2% of a scan's mean lesion-location frequency. Scans from participants with mild cognitive impairment or dementia do not contribute to the maps and are unchanged. b, Is the mean the right summary? Each scan's score starts as the average reference frequency across its lesion voxels. An average can be pulled down by a few voxels in very rare locations, so three other summaries recorded by the same pipeline were tested: the median, the proportion of lesion voxels in rare locations, and the proportion in common locations. Each was put through the same burden-and-age adjustment, and the bars show how much of each summary that adjustment accounts for. The mean is the summary the adjustment explains best, and the median gives almost the same final score (r = 0.96), indicating that the resulting scores are largely insensitive to the choice between mean and median. c, Is the vTRM score stable within a person? Because the vTRM score is what remains after burden and age are removed, its reproducibility should be established rather than assumed. Among 6,001 participants who were cognitively unimpaired at every visit and had two or more scans, the intraclass correlation shows how much of the total variation lies between people rather than between repeat scans of the same person. The vTRM score reaches 0.79, close to the 0.86 obtained for WMH volume, a directly measured quantity included as a benchmark. The score therefore shows substantial within-participant reproducibility. Full results are given in Supplementary Table S5.T2, shown below.

**a Self-inclusion**

| group | n | median_freq_incl | median_freq_excl | median_abs_freq_diff | median_rel_diff_pct | median_abs_vtrm_diff | max_abs_vtrm_diff |
|---|---|---|---|---|---|---|---|
| CN (in the maps) | 27247 | 0.075 | 0.0748 | 0.0001 | 0.2017 | 0.0001 | 0.0099 |
| MCI/AD (not in the maps) | 14191 | 0.0834 | 0.0834 | 0.0 | | 0.0 | 0.0 |

**b Choice of summary statistic**

| variant_a | variant_b | pearson_r | spearman_rho |
|---|---|---|---|
| mean | median | 0.9591 | 0.9576 |
| mean | pct_rare | -0.6311 | -0.6611 |
| mean | pct_common | 0.851 | 0.8819 |
| median | pct_rare | -0.6275 | -0.6731 |
| median | pct_common | 0.8397 | 0.8913 |
| pct_rare | pct_common | -0.773 | -0.7684 |

**c Repeatability**

| measure | column | n_participants | pearson_first_last | spearman_first_last | within_participant_var | between_participant_var | prop_variance_between | median_within_sd |
|---|---|---|---|---|---|---|---|---|
| vTRM score | vtrm_raw | 6001 | 0.736 | 0.7439 | 0.0001 | 0.0005 | 0.7918 | 0.0062 |
| vTRM percentile | vtrm_pct | 6001 | 0.7104 | 0.7111 | 194.385 | 672.884 | 0.7759 | 7.6421 |
| mean lesion-location frequency | mean_freq | 6001 | 0.862 | 0.8509 | 0.0003 | 0.0017 | 0.8513 | 0.0101 |
| log WMH burden | log_wmh | 6001 | 0.8466 | 0.8433 | 0.2556 | 1.5513 | 0.8586 | 0.301 |

#### F3. Age-interpolated lookup.

The reference frequency maps are built in six age bands, but age is continuous. Assigning each scan the map for its band would give the same reference to everyone within a band and change it abruptly at each boundary, so each scan's reference instead blends the two maps whose median cognitively unimpaired ages bracket its own age. This item quantifies what that choice avoids, comparing median lesion-location frequency by year of age under both lookups and measuring the discontinuity at each band boundary. Under band assignment, people one year apart in age differ substantially at the boundaries; interpolation reduces this to the change expected over a year of age.

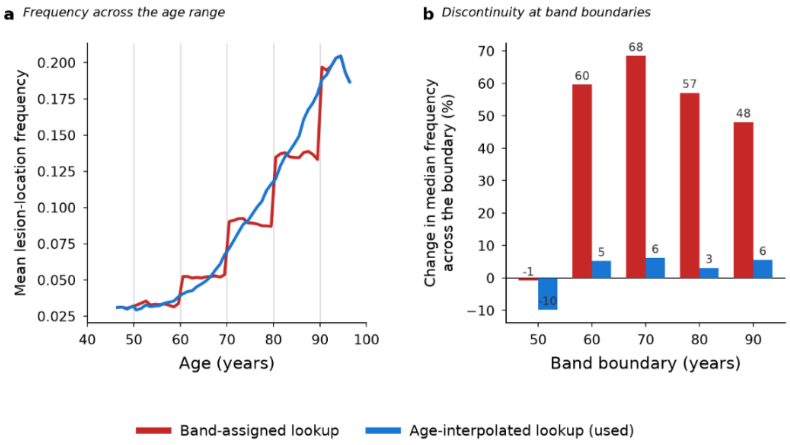

**Figure S5.F3 | Age-interpolated frequency lookup**. The reference frequency maps are built in six age bands, but age is continuous. Assigning each scan the map for the band it falls in would give the same reference to everyone within a band and change it abruptly at each boundary, so each scan's reference instead blends the two maps whose median cognitively unimpaired ages bracket its own age. a, Median mean lesion-location frequency by year of age under each lookup. The band-assigned version rises in steps at the boundaries marked in grey, whereas the interpolated version increases smoothly. b, Median frequency measured in scans aged within one year below and one year above each boundary, expressed as the change across the boundary. Under band assignment, people one year apart in age differ by 48% to 68% in mean lesion-location frequency at the boundaries at 60, 70, 80 and 90 years; interpolation reduces this to 3% to 6%, which is the change expected over a year of age. Across all scans the two lookups differ by a median of 9.6% in mean frequency, largest in the seventh and eighth decades where the bands are widest relative to the rate of change, and near zero below 50 and above 90 where interpolation reverts to the nearest map. Full results are given in Supplementary Table S5.T3, shown below.

| group | n | n_finite | median_abs_rel_diff_pct | p95_abs_rel_diff_pct | max_abs_rel_diff_pct | view | edge_age | n_below | n_above | band_jump_pct | interp_jump_pct |
|---|---|---|---|---|---|---|---|---|---|---|---|
| all scans | 41438.0 | 41434.0 | 9.587 | 38.398 | 218.483 | by band | | | | | |
| lt50 | 296.0 | 296.0 | 0.051 | 11.679 | 79.196 | by band | | | | | |
| 50s | 3456.0 | 3455.0 | 5.905 | 29.386 | 85.834 | by band | | | | | |
| 60s | 11702.0 | 11700.0 | 10.498 | 37.493 | 107.343 | by band | | | | | |
| 70s | 18116.0 | 18115.0 | 11.816 | 43.158 | 218.483 | by band | | | | | |
| 80s | 7126.0 | 7126.0 | 7.554 | 32.716 | 85.449 | by band | | | | | |
| 90plus | 742.0 | 742.0 | 0.0 | 6.059 | 11.089 | by band | | | | | |
| | | | | | | band edge | 50.0 | 62.0 | 161.0 | -0.8 | -9.85 |
| | | | | | | band edge | 60.0 | 580.0 | 628.0 | 59.59 | 5.21 |
| | | | | | | band edge | 70.0 | 1821.0 | 2034.0 | 68.44 | 6.18 |
| | | | | | | band edge | 80.0 | 1321.0 | 1239.0 | 57.09 | 3.09 |
| | | | | | | band edge | 90.0 | 290.0 | 228.0 | 48.02 | 5.54 |

#### F4. Burden-and-age conditioning

Mean lesion-location frequency falls as lesion masks grow, because larger masks necessarily extend into less commonly affected voxels. The vTRM score removes this by subtracting the frequency expected for a person's WMH burden and age, so the specification of that expectation determines whether the score reflects lesion location or residual lesion quantity. Two models were compared: an additive model, in which expected frequency is a function of burden plus a function of age and the burden adjustment is a single fixed curve applied at every age; and a tensor-product model, in which the two spline expansions are multiplied, allowing the shape of the burden curve to differ with age. Under the additive model the residual is uncorrelated with burden across the sample as a whole, but a relationship remains within age bands, negative in the sixth decade and positive in the ninth and tenth. That reversal of sign is the signature of a burden–frequency relationship that changes with age, which a single fixed curve cannot fit. The tensor model removes it, explains slightly more variance, and was adopted. The two scores nonetheless agree closely, so individual scores change only marginally; the distinction matters because only the tensor specification supports the claim that the score reflects where lesions are located rather than how many lesion voxels are present.

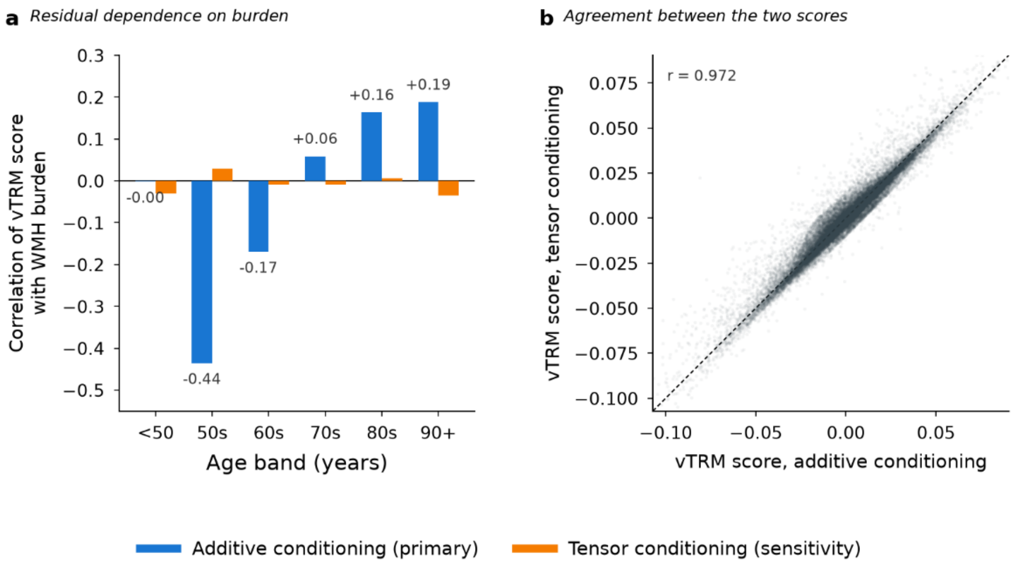

**Figure S5.F4 | Conditioning the vTRM score on lesion burden and age.** Mean lesion-location frequency falls as lesion masks grow, because larger masks necessarily extend into less commonly affected voxels. The vTRM score removes this by subtracting the frequency expected for a person's WMH burden and age, estimated by a spline regression fitted in cognitively unimpaired participants; the score is what remains, observed minus expected. Two ways of modelling the expected frequency were compared. In the additive model, expected frequency is a function of burden plus a function of age, so the burden adjustment is a single fixed curve applied at every age: the burden-specific adjustment for 5 $cm^3$ is the same at ages 55 and 85, although the age-specific contribution differs. In the tensor-product model, the two spline expansions are also multiplied together, allowing the shape of the burden curve to differ with age. a, Correlation between the vTRM score and log WMH volume, calculated separately within each age band. Under the additive model the residual is uncorrelated with burden across the sample as a whole, but within age bands a relationship remains, negative in the sixth decade ($r = -0.44$) and positive in the ninth and tenth ($r = +0.16$ and $+0.19$). The reversal of sign is the signature of a burden–frequency relationship that changes with age: at younger ages higher burden leaves a lower residual, at older ages a higher one, and a single fixed curve cannot fit both, so it splits the difference and is wrong in opposite directions at each end. The tensor model, which contains the interaction, fits both: no age band exceeds $r = 0.04$ and none is statistically significant. It also explains slightly more variance in mean frequency ($R^2 = 0.698$ against 0.679), and was adopted. b, The two scores nonetheless agree closely ($r = 0.972$ in cognitively unimpaired scans), so individual scores change only marginally. The distinction matters because only the tensor specification supports the statement that the score reflects where lesions are located rather than how many lesion voxels are present. Full results are given in Supplementary Table S5.T4; correlations by age band are shown below, and the tensor-versus-additive downstream comparison and scan-level burden independence results are provided as separate files, S5.T4b.csv and S5.T4c.csv.

| age_band | n | r_before_conditioning | r_after_conditioning | p_after | spearman_after | r_squared_after_pct | view | decile | median_wmh_cm3 | mean_vtrm | median_vtrm |
|---|---|---|---|---|---|---|---|---|---|---|---|
| <50 | 256 | 0.0706 | -0.0019 | 0.976 | -0.0172 | 0.0 | age band | | | | |
| 50s | 2560 | -0.1521 | -0.4354 | 0.0 | -0.4783 | 18.96 | age band | | | | |
| 60s | 8523 | 0.0309 | -0.1694 | 0.0 | -0.2029 | 2.87 | age band | | | | |
| 70s | 12124 | 0.0781 | 0.0589 | 0.0 | 0.0426 | 0.35 | age band | | | | |
| 80s | 3520 | 0.0239 | 0.1633 | 0.0 | 0.1309 | 2.67 | age band | | | | |
| 90+ | 264 | -0.0655 | 0.1881 | 0.0022 | 0.0873 | 3.54 | age band | | | | |
| | 2738 | | | | | | burden decile | 1.0 | 0.2433 | 0.0006 | 0.0029 |
| | 2721 | | | | | | burden decile | 2.0 | 0.592 | 0.001 | 0.0027 |
| | 2715 | | | | | | burden decile | 3.0 | 1.0371 | -0.001 | -0.0001 |
| | 2731 | | | | | | burden decile | 4.0 | 1.549 | -0.0009 | 0.0004 |
| | 2726 | | | | | | burden decile | 5.0 | 2.2 | -0.0013 | -0.0008 |
| | 2717 | | | | | | burden decile | 6.0 | 3.032 | 0.0006 | 0.0002 |
| | 2725 | | | | | | burden decile | 7.0 | 4.1811 | 0.0005 | 0.0 |
| | 2724 | | | | | | burden decile | 8.0 | 5.9878 | -0.0003 | -0.0011 |
| | 2726 | | | | | | burden decile | 9.0 | 9.352 | 0.0005 | 0.0007 |
| | 2724 | | | | | | burden decile | 10.0 | 19.6026 | 0.0003 | 0.0019 |

#### F5. Cohort coding and out-of-sample scoring

The vTRM percentile model includes an indicator for each contributing study, but a scan from outside the reference sample has no such indicator. The indicators are therefore coded to sum to zero, so that the intercept is the unweighted mean across studies rather than the value for one arbitrarily chosen reference study, and a scan with all cohort terms set to zero is compared against that average. This item establishes what an external scan loses by being scored that way. Every scan in the reference sample was scored twice, once at its own study and once with the cohort terms set to zero, and the two percentiles compared. Shifts are modest and balanced around zero, and omitting the cohort terms altogether, which weights studies by sample size rather than equally, gives a nearly identical result.

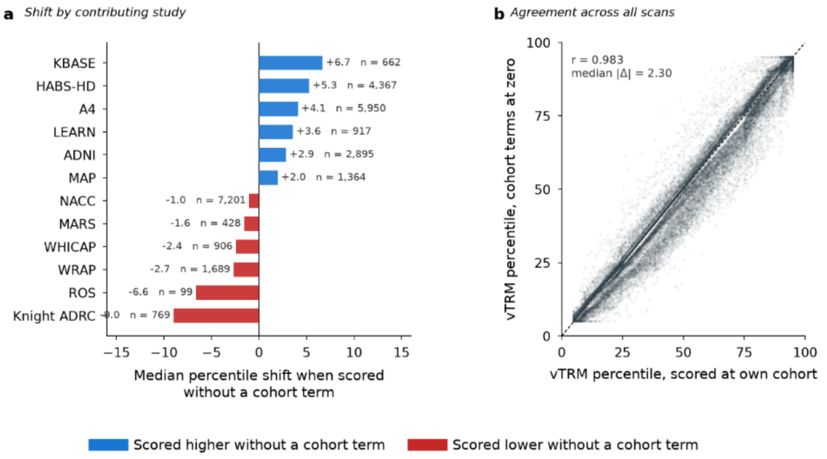

**Figure S5.F5 | Cohort coding and scoring of scans from outside the reference sample.** The vTRM percentile model includes an indicator for each contributing study, coded so that the indicators sum to zero. The intercept is then the unweighted mean across studies rather than the value for one arbitrarily chosen reference study, and a scan with all cohort terms set to zero is compared against that average. This is how a scan from outside the reference sample is scored, since no cohort term is available for it. a, For each study, the median difference between the percentile a scan receives when scored at its own study and the percentile it receives with the cohort terms set to zero, with the number of cognitively unimpaired scans that study contributes to the reference. Shifts are modest and balanced around zero, ranging from −9.0 percentile points for Knight ADRC to +6.7 for KBASE; the two largest are among the smallest contributors. b, The two scorings agree closely across all scans (r = 0.983, median absolute difference 2.30 percentile points, 9.2% differing by more than 10). Omitting the cohort terms altogether, which weights studies by their sample size rather than equally, gives a nearly identical result (r = 0.985, median absolute difference 2.45 points), so the choice of centering has little practical consequence. A scan scored externally therefore receives a percentile within a few points of the value it receives within the reference sample. Full results are given in Supplementary Table S5.T5, shown below.

| comparison | n | pearson_r | spearman_rho | median_abs_diff | pct_gt10 | view | cohort | n_scans | n_CN_in_reference | median_own_minus_zero | median_abs_own_minus_zero | pct_gt10_own_vs_zero | median_own_minus_pooled |
|---|---|---|---|---|---|---|---|---|---|---|---|---|---|
| own cohort vs cohort terms at zero | 41409.0 | 0.9831 | 0.9839 | 2.3 | 9.2 | overall | | | | | | | |
| own cohort vs no cohort terms | 41409.0 | 0.9849 | | 2.45 | 4.2 | overall | | | | | | | |
| | | | | | | by cohort | Knight ADRC | 908.0 | 769.0 | -8.99 | 8.99 | 46.5 | -10.2 |
| | | | | | | by cohort | ROS | 170.0 | 99.0 | -6.63 | 6.63 | 30.0 | -7.9 |
| | | | | | | by cohort | WRAP | 1865.0 | 1689.0 | -2.69 | 2.73 | 0.0 | -5.02 |
| | | | | | | by cohort | WHICAP | 1475.0 | 906.0 | -2.43 | 2.43 | 0.0 | -4.76 |
| | | | | | | by cohort | MARS | 584.0 | 428.0 | -1.57 | 1.72 | 0.2 | -3.46 |
| | | | | | | by cohort | NACC | 13021.0 | 7201.0 | -1.03 | 1.03 | 0.0 | -2.65 |
| | | | | | | by cohort | MAP | 2293.0 | 1364.0 | 1.99 | 1.99 | 0.0 | 0.78 |
| | | | | | | by cohort | ADNI | 7198.0 | 2895.0 | 2.86 | 2.86 | 10.0 | 1.16 |
| | | | | | | by cohort | LEARN | 917.0 | 917.0 | 3.57 | 3.57 | 8.2 | 2.01 |
| | | | | | | by cohort | A4 | 5950.0 | 5950.0 | 4.09 | 4.09 | 20.9 | 2.64 |
| | | | | | | by cohort | HABS-HD | 5858.0 | 4367.0 | 5.27 | 5.27 | 15.8 | 2.94 |
| | | | | | | by cohort | KBASE | 1170.0 | 662.0 | 6.68 | 6.68 | 32.3 | 5.34 |

#### F6. Associations between vTRM and cognitive, biomarker and vascular measures

As with burden percentile, the score's value depends on whether it relates to measures of brain health, and specifically on whether it carries information that lesion volume does not. Every scan was assigned a vTRM score and related to the available cognitive, biomarker and vascular measures, with partial correlations adjusted for age, sex and total WMH volume so that the association reflects lesion location rather than lesion quantity. The corresponding burden-percentile associations are reported alongside, adjusted for age and sex only; because the burden percentile is itself a rescaling of log volume given age and sex, adjusting it for volume would remove the variable's own content, so the two columns are not directly comparable.

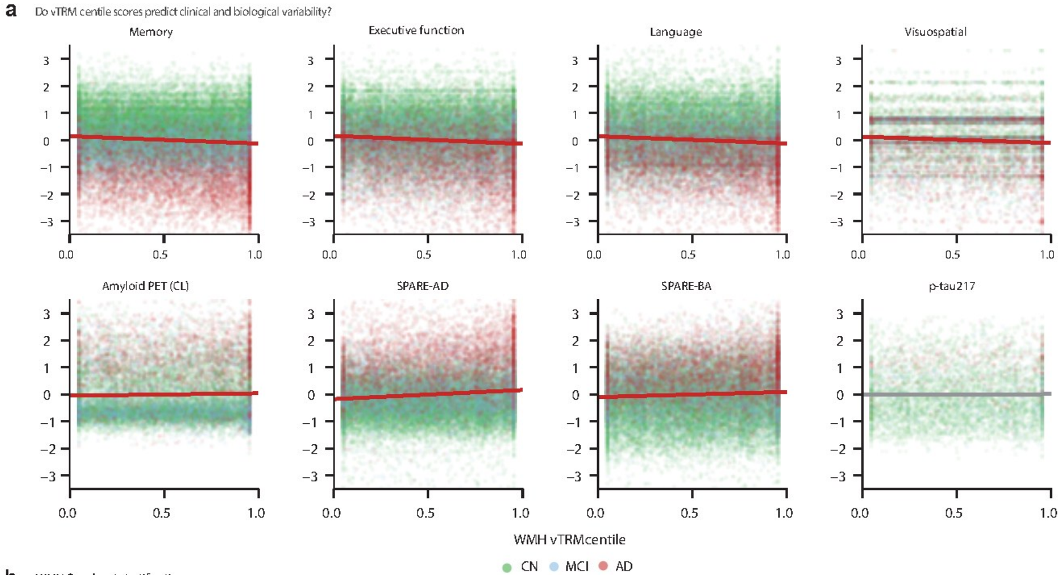

**Figure S5.F6 | Associations between vTRM and cognitive, biomarker and vascular measures.** Every scan was assigned a vTRM score against the age- and sex-matched cognitively unimpaired frequency maps, conditioned on total WMH burden and age, and related to available cognitive, biomarker and vascular measures. Each panel plots the vTRM percentile against one outcome, with outcomes z-scored. Top row: memory, executive function, language and visuospatial performance. Bottom row: amyloid PET Centiloids, SPARE-AD, SPARE-BA and plasma p-tau217. After adjustment for age, sex and total WMH volume, higher vTRM was associated with greater SPARE-AD ($r = 0.113$, 95% CI 0.102 to 0.124, $n = 29{,}644$) and SPARE-BA (0.073, 0.062 to 0.084, $n = 29{,}644$), and with poorer memory (−0.087, −0.097 to −0.077, $n = 38{,}648$), executive function (−0.100, −0.111 to −0.089, $n = 31{,}761$), language (−0.093, −0.104 to −0.082, $n = 31{,}754$) and visuospatial performance (−0.061, −0.079 to −0.043, $n = 11{,}781$). Associations were also present with amyloid PET Centiloids (0.030, 0.015 to 0.045, $n = 16{,}910$) and CSF Aβ42 (−0.044, −0.075 to −0.013, $n = 3{,}920$), whereas vascular burden showed no association (0.010, −0.010 to 0.030, $P = 0.35$, $n = 9{,}192$). All associations except vascular burden were statistically significant. Correlations range from 0.01 to 0.11 in absolute value, so the vTRM score accounts for a small proportion of variance in each measure while remaining independent of total lesion volume by construction. Partial correlations, confidence intervals and sample sizes, reported alongside the corresponding burden-percentile associations, are given in Supplementary Table S5.T6, shown below.

| outcome | n | vtrm_r | vtrm_lo | vtrm_hi | vtrm_p | burden_r | burden_lo | burden_hi | burden_p |
|---|---|---|---|---|---|---|---|---|---|
| SPARE_AD | 29644 | 0.113 | 0.1017 | 0.1242 | 4.1e-85 | 0.19 | 0.179 | 0.2009 | 1.4e-239 |
| SPARE_BA | 29644 | 0.073 | 0.0617 | 0.0843 | 1.5e-36 | 0.273 | 0.2624 | 0.2835 | 0.0e+00 |
| Memory | 38648 | -0.087 | -0.0969 | -0.0771 | 9.4e-66 | -0.094 | -0.1039 | -0.0841 | 2.9e-76 |
| Executive | 31761 | -0.1 | -0.1109 | -0.0891 | 2.4e-71 | -0.069 | -0.0799 | -0.058 | 1.1e-34 |
| Language | 31754 | -0.093 | -0.1039 | -0.0821 | 1.6e-61 | -0.055 | -0.066 | -0.044 | 1.3e-22 |
| Visuospatial | 11781 | -0.061 | -0.079 | -0.043 | 4.4e-11 | -0.065 | -0.083 | -0.047 | 1.8e-12 |
| Amyloid_CL | 16910 | 0.03 | 0.0149 | 0.0451 | 1.2e-04 | 0.136 | 0.1212 | 0.1508 | 1.4e-70 |
| CSF_AB42 | 3920 | -0.044 | -0.0752 | -0.0127 | 6.0e-03 | -0.177 | -0.2072 | -0.1465 | 7.5e-29 |
| Vascular_PC1 | 9192 | 0.01 | -0.0104 | 0.0304 | 3.5e-01 | 0.021 | 0.0006 | 0.0414 | 4.0e-02 |

**Supplementary Table S5.T6 | Associations between vTRM and burden percentile and cognitive, biomarker and vascular measures.** Partial correlations with 95% confidence intervals, P values and sample sizes for each outcome. The vTRM score is adjusted for age, sex and total WMH volume. The burden percentile is adjusted for age and sex only, because it is itself a rescaling of log volume given age and sex and adjusting it for volume would remove the variable's own content. The two columns are therefore reported side by side for reference and are not directly comparable.

#### F7. Age dependence of vTRM variability.

The vTRM percentile charts show a normative band that is wide at the youngest ages, narrows through middle adulthood and widens again thereafter, and the main text describes this shape. Because the reference is fitted with age and age squared, the width of the band between any two percentile curves is itself a quadratic function of age, so a U shape is what that specification will produce whenever the fitted percentiles fan out at both ends of the range. This analysis distinguishes model-derived patterns from those supported by the observed data. The fitted band width was computed analytically from the exported quantile regression coefficients and compared against the dispersion observed directly in sliding age windows. The two agree above the trough, where the sample is dense; below it the observed dispersion is flat while the fitted band continues to widen, so the left arm of the U is a property of the quadratic form in a sparsely sampled age range rather than a measured increase in variability.

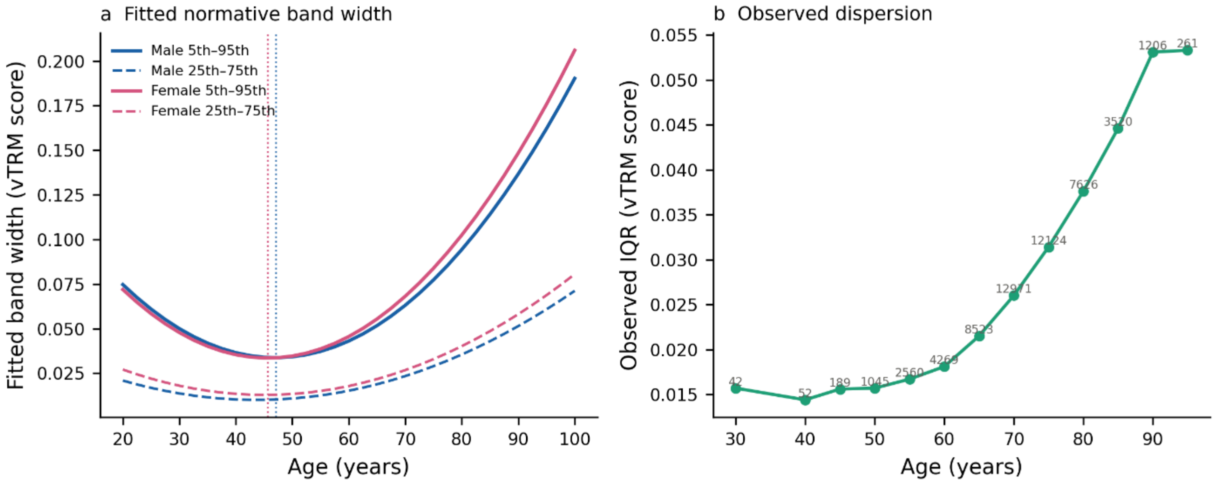

**Figure S5.F7 | The U-shaped age dependence of vTRM variability.** a, Width of the fitted 5th–95th and 25th–75th percentile bands across age, in men and women; dotted lines mark the minimum. The band narrows 2.2-fold from age 20 to a minimum at 47.1 years in men and 45.6 years in women, then widens 5.6-fold by age 100. At younger ages, fitted percentile bands are wider, although this widening is not supported by the observed dispersion in the sparsely sampled younger age range; the band is narrowest in middle adulthood, where lesion burden is sufficient to stabilise the estimate and the sample is dense; and it widens thereafter as lesion topography diverges between individuals. b, Interquartile range of the observed vTRM score in sliding 10-year windows, with the number of contributing scans annotated. The observed dispersion reproduces the widening above the trough but is flat below age 50, where only 42 to 189 scans contribute per window. Fitted band widths at 2.5-year intervals from 20 to 100 by sex, computed from the exported quantile regression coefficients with cohort terms at zero, together with the observed scan count, interquartile range and median lesion-voxel count per window, are provided as a separate file, S5.T7.csv.

#### S6. Two-axis stratification

##### F1. Independence of the burden and vTRM axes, and group occupancy.

The two-axis framework only adds information if the axes measure different things. Burden percentile quantifies how much lesion volume a scan carries for its age and sex; vTRM percentile quantifies how atypical the lesion locations are given that volume. This item establishes that they are close to independent and that the resulting four groups are populated as independence would predict. The two percentiles were correlated across all scans with both, overall and within diagnostic group, and group sizes compared against those expected if the two dichotomies were independent. The axes share less than one percent of their variance, group sizes are close to expectation, and the ratio of AD to cognitively unimpaired scans rises monotonically across the four groups.

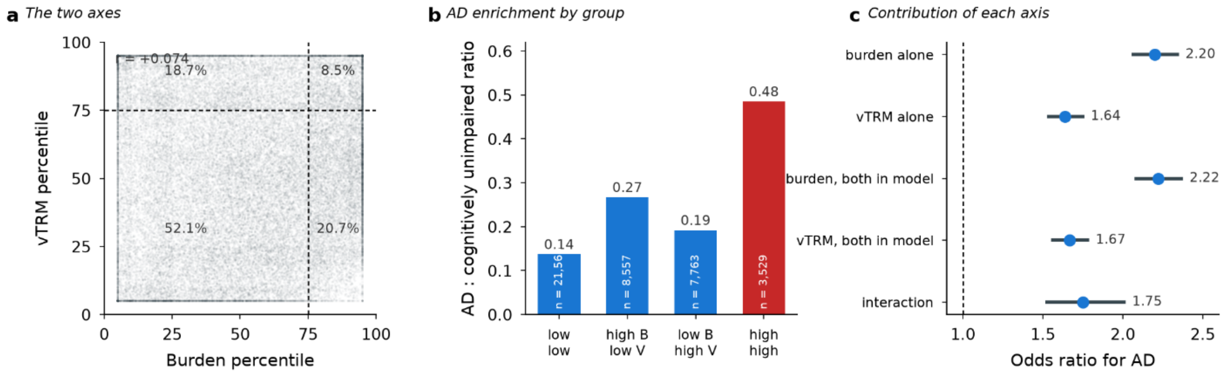

**Figure S6.F1 | Burden and vTRM percentiles capture largely distinct dimensions of WMH.** The two-axis framework treats WMH burden percentile, which measures how much lesion volume a scan carries for its age and sex, and vTRM percentile, which measures how atypical the lesion locations are, as separate dimensions. a, The two percentiles plotted against each other for all 41,409 scans with both, with the upper-quartile boundaries marked and the percentage of the sample falling in each group annotated. The axes are weakly correlated (r = 0.074 across all scans; 0.051 in cognitively unimpaired), so less than 1% of the variance in either is shared with the other. Group sizes are close to those expected if the two dichotomies were independent, with observed-to-expected ratios of 0.97 to 1.06, and the two off-diagonal groups contain 8,557 and 7,763 scans. b, The ratio of AD to cognitively unimpaired scans in each group, rising from 0.14 where both axes are low to 0.49 where both are high, a 3.5-fold enrichment. Full results are given in Supplementary Tables S6.T1a-d.

#### F2. Stability, cost of dichotomisation, and outcomes at matched burden.

Splitting two continuous axes at the upper quartile creates interpretable groups but discards information, and three properties needed establishing before the grouping could be used. First, whether group membership is stable within a person, measured among participants with repeat scans. Second, what the dichotomisation costs in discrimination, by comparing the four groups against the two percentiles entered as continuous predictors. Third, and most important, whether lesion location carries information at matched lesion volume: the two high-burden groups differ only in whether lesion locations are atypical, so comparing them on outcomes that took no part in constructing either axis isolates the contribution of location. Group membership is largely stable, dichotomisation costs a small amount of discrimination, and atypical location at matched burden is associated with poorer cognition and higher structural neurodegeneration indices while amyloid and plasma p-tau217 do not differ appreciably.

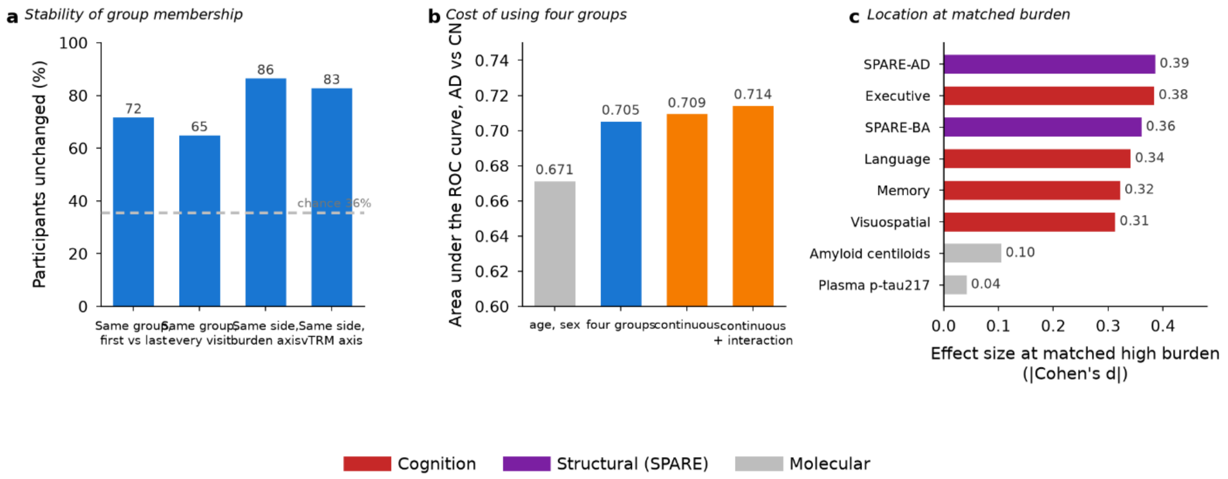

**Figure S6.F2 | Utility of the four-group framework.** Three properties were examined beyond the validity checks of Supplementary Figure S6.F1. a, Stability of group membership among the 6,001 participants who were cognitively unimpaired at every visit and had two or more scans. 71.7% were in the same group at their first and last scan, against 35.6% expected if groups were assigned at random given their sizes, and each axis separately placed participants on the same side of its cut in 83% to 87% of cases. b, Discrimination of AD from cognitively unimpaired status by logistic models adjusted for age and sex. The four groups achieve an area under the ROC curve of 0.705, above the 0.671 obtained from age and sex alone, but below the 0.709 obtained from the two percentiles entered as continuous predictors. Dichotomizing at the upper quartile therefore costs a small amount of discrimination, which is the price of an interpretable grouping. c, The comparison that isolates lesion location: scans in the two groups with high WMH burden, distinguished by their vTRM percentiles and differ only in whether lesion locations are atypical, compared on outcomes that took no part in constructing either axis. Atypical location is associated with poorer performance in all four cognitive domains and with higher structural neurodegeneration indices; within AD scans alone the differences are larger. Amyloid Centiloids and plasma p-tau217 do not differ appreciably between the two groups, indicating that the location axis relates to cognitive and structural outcomes rather than to molecular AD pathology. Stability, discrimination and outcome-comparison results are given in Supplementary Table S6.T2, shown below; outcome comparisons between the two high-burden groups are provided as a separate file, S6.T2c.csv.

**a Stability**

| measure | value | n |
|---|---|---|
| identical group at every visit | 64.9 | 6001.0 |
| same group at first and last visit | 71.7 | 6001.0 |
| same side of the burden cut, first vs last | 86.5 | 6001.0 |
| same side of the vTRM cut, first vs last | 82.8 | 6001.0 |
| expected agreement if assigned at random | 35.6 | |

**b Cost of dichotomisation**

| model | n | n_terms | pseudo_r2 | auc | aic |
|---|---|---|---|---|---|
| groups (both + interaction) | 32530 | 5 | 0.0829 | 0.7051 | 26481.2 |
| continuous (both) | 32530 | 4 | 0.0896 | 0.7092 | 26286.9 |
| continuous (both + interaction) | 32530 | 5 | 0.0928 | 0.714 | 26197.0 |
| age and sex only | 32530 | 2 | 0.057 | 0.6712 | 27224.9 |

**d Prognosis**

| group | n | median_dlog_per_year | mean_dlog_per_year | median_followup_yr |
|---|---|---|---|---|
| LowB/LowV | 3201 | 0.1078 | 0.1116 | 4.4 |
| HighB/LowV | 1189 | 0.0574 | 0.0401 | 5.2 |
| LowB/HighV | 1195 | 0.0798 | 0.0752 | 4.5 |
| HighB/HighV | 250 | 0.0628 | 0.0248 | 5.0 |

#### F3. Quadrant characterisation: clinical, biomarker and vascular profiles

Having established that the four groups are populated and stable, this item asks what distinguishes the people in them. Each group was profiled on demographic, cognitive, biomarker and vascular measures, with continuous variables compared by Kruskal–Wallis test and categorical variables by chi-squared test. Profiling was performed across all scans and repeated within AD scans alone, so that disease-related variation can be distinguished from normative variation. The comparison of interest remains the two high-burden groups, which are matched on lesion volume and differ only in whether lesion locations are atypical. Supplementary Table S6.T3 reports medians and group sizes for every measure, across all diagnoses and within AD. The comparison of interest is between the two high-burden groups, which are matched on lesion volume and differ only in whether lesion locations are atypical: median WMH volume is 14.9 $cm^3$ in HighB/LowV against a comparable value in HighB/HighV, while cognitive scores and the SPARE indices differ substantially between them. Amyloid positivity, APOE4 dose and the diagnostic composition of each group are given in the same table. Provided as separate files, S6.T3a.csv and S6.T3b.csv.

#### F4. Spatial composition by quadrant

The two axes are defined by burden and by a single summary of lesion-location typicality, neither of which specifies where the lesions of each group actually lie. Regional WMH fractions were therefore compared across the four groups to establish where the lesions of each group are located. Supplementary Table S6.T4 gives the mean percentage of each scan's lesion volume falling in the periventricular and ventricle-distal compartments, the four lobar compartments, and the three arterial territories, across all diagnoses and within AD scans. The two axes separate spatially in different ways. High burden is accompanied by a shift away from periventricular white matter, from 84.9% in LowB/LowV to 69.5% in HighB/LowV, and toward parietal and MCA-territory locations. High atypicality at low burden shows a different pattern, with a comparable periventricular reduction to 71.2% but an increase in temporal involvement, from 5.4% to 8.4%, and no accompanying rise in MCA territory.

**a All diagnoses**

| quadrant | N | PV_mean_pct | Distal_mean_pct | Frontal_mean_pct | Parietal_mean_pct | Occipital_mean_pct | Temporal_mean_pct | MCA_mean_pct | PCA_mean_pct | ACA_mean_pct |
|---|---|---|---|---|---|---|---|---|---|---|
| LowB/LowV | 21028 | 84.9 | 15.1 | 44.3 | 18.4 | 19.4 | 5.4 | 36.5 | 21.2 | 29.1 |
| HighB/LowV | 8262 | 69.5 | 30.5 | 45.1 | 30.6 | 14.7 | 4.9 | 46.9 | 16.4 | 30.9 |
| LowB/HighV | 7501 | 71.2 | 28.8 | 38.7 | 18.5 | 18.1 | 8.4 | 34.8 | 22.2 | 27.2 |
| HighB/HighV | 3325 | 50.6 | 49.4 | 48.7 | 26.9 | 14.6 | 5.5 | 50.7 | 16.1 | 28.4 |

**b AD only**

| quadrant | N | PV_mean_pct | Distal_mean_pct | Frontal_mean_pct | Parietal_mean_pct | Occipital_mean_pct | Temporal_mean_pct | MCA_mean_pct | PCA_mean_pct | ACA_mean_pct |
|---|---|---|---|---|---|---|---|---|---|---|
| LowB/LowV | 2106 | 77.8 | 22.2 | 46.4 | 22.8 | 18.5 | 5.2 | 42.9 | 19.3 | 29.5 |
| HighB/LowV | 1364 | 63.2 | 36.8 | 46.5 | 30.7 | 14.6 | 5.2 | 48.4 | 15.8 | 31.3 |
| LowB/HighV | 981 | 62.5 | 37.5 | 41.4 | 21.4 | 18.1 | 8.8 | 40.4 | 21.1 | 28.6 |
| HighB/HighV | 832 | 45.7 | 54.3 | 49.0 | 25.9 | 16.4 | 5.9 | 50.8 | 16.9 | 28.8 |

#### S7. Applying the framework to new scans

The published reference model includes an indicator for each contributing study. A scan from outside the reference sample has no such indicator, so the distributed scoring package uses a cohort-free version of the model in which those terms are omitted. To establish what this costs, every scan in the reference sample was scored twice, once against the full model at its own study and once against the cohort-free package, and the two percentiles compared. Supplementary Table S7.T1 gives the mean difference and standard deviation for each axis, together with the percentage of scans that cross the 75$^{th}$ percentile threshold under one scoring but not the other, by contributing study and by age stratum. Differences are modest across studies: mean burden shifts range from 4.99 percentile points in MARS to 5.75 in ROS, with 3.3% to 7.1% of scans crossing the burden threshold and 2.9% to 5.7% the vTRM threshold. Shifts are larger at the youngest ages, where few cognitively unimpaired scans contribute to the reference: mean burden differences reach 40.2 percentile points in the third decade and 32.7 in the fourth, falling to 13.0 in the fifth. Scores for scans below approximately 50 years should therefore be interpreted with caution. A worked example of the scoring output for a single test scan is provided as a separate file, S7.T1a-c.csv.

[72] Bao YW, Ji YQ, Wang LN, Zhou Y, Wang F, Shen DM, Pylypenko D, Tong Q, Mak KF, Guo LL. Topography of Juxtaventricular white matter hyperintensities and cognitive associations in Alzheimer's disease: a dual-cohort study.

Eur Radiol Exp. 2026 Jul 29;10(1):109. doi: 10.1186/s41747-026-00776-1. PMID: 42525186; PMCID: PMC13421530.

[73] Phuah CL, Chen Y, Strain JF, Yechoor N, Laurido-Soto OJ, Ances BM, Lee JM; Alzheimer's Disease Neuroimaging Initiative. Association of Data-Driven White Matter Hyperintensity Spatial Signatures With Distinct Cerebral Small Vessel Disease Etiologies. Neurology. 2022 Dec 5;99(23):e2535-e2547. doi: 10.1212/WNL.0000000000201186. PMID: 36123127; PMCID: PMC9754646.
